# Deep Mutational Scanning in Disease-related Genes with Saturation Mutagenesis-Reinforced Functional Assays (SMuRF)

**DOI:** 10.1101/2023.07.12.548370

**Authors:** Kaiyue Ma, Shushu Huang, Kenneth K. Ng, Nicole J. Lake, Soumya Joseph, Jenny Xu, Angela Lek, Lin Ge, Keryn G. Woodman, Katherine E. Koczwara, Justin Cohen, Vincent Ho, Christine L. O’Connor, Melinda A. Brindley, Kevin P. Campbell, Monkol Lek

## Abstract

Interpretation of disease-causing genetic variants remains a challenge in human genetics. Current costs and complexity of deep mutational scanning methods hamper crowd-sourcing approaches toward genome-wide resolution of variants in disease-related genes. Our framework, Saturation Mutagenesis-Reinforced Functional assays (SMuRF), addresses these issues by offering simple and cost-effective saturation mutagenesis, as well as streamlining functional assays to enhance the interpretation of unresolved variants. Applying SMuRF to neuromuscular disease genes *FKRP* and *LARGE1*, we generated functional scores for all possible coding single nucleotide variants, which aid in resolving clinically reported variants of uncertain significance. SMuRF also demonstrates utility in predicting disease severity, resolving critical structural regions, and providing training datasets for the development of computational predictors. Our approach opens new directions for enabling variant-to-function insights for disease genes in a manner that is broadly useful for crowd-sourcing implementation across standard research laboratories.

## Introduction

Recent advancement in Next-generation sequencing (NGS) and population studies^1–4^ together with establishment of large biobanks have improved disease-associated variant detection and interpretation^5–7^. However, for many genetic disease patients, it remains a challenge to pinpoint their specific disease-causing variant(s), hampering access to approved treatments, participation in clinical trials, options for disease management and family planning^8,9^. The emergence of gene therapies has further heightened the importance of finding diagnoses for genetic disease patients as an initial and crucial prerequisite towards clinical trial readiness. The variants discovered in patients that are difficult to interpret are classified as variants of uncertain significance (VUS), which is often a result of insufficient evidence to determine pathogenicity^10^. The rate of VUS observed in disease-associated genes is typically higher in individuals of non-European ancestry due to limited diversity in biomedical databases^11^.

To provide an additional line of evidence for the interpretation of variants, deep mutational scanning (DMS) was proposed to unbiasedly generate functional scores for all possible variants^12^. DMS employs a pooled model cell population consisting of large amounts of variants, which are subsequently characterized with appropriate high-throughput functional assays^13^. The collective functional read-out from each variant in the gene of interest gives rise to the generation of an “Atlas of Variant Effects”^14^. However, despite its promise, the high cost and technical complexity of performing DMS discourage many researchers, especially those in modestly funded rare disease labs.

In saturation mutagenesis, well-established methods include the insertion of variant-carrying tiles^15^, saturation genome editing (multiplex homology-directed repair)^16^, reversibly-terminated inosine mutagenesis^17^, and recently emerged saturation prime editing^18^. However, these methods often come with limitations like intensive labor requirements, high expenses, disparate variant representation, and limited spanning regions. Hence, there is a need for a simple and cost-effective saturation mutagenesis method that provides comprehensive and unbiased coverage. Programmed Allelic Series (PALS) was previously developed as a saturation mutagenesis method with relatively simple steps and unbiased coverage^19^. PALS had the potential to reduce costs by using minimized-length oligos instead of long DNA fragments to introduce variants. However, its reliance on special reagents has limited its widespread adoption in DMS studies. A previous optimization of PALS removed the need for special reagents but transformed the method from requiring only a single-tube reaction to necessitating independent reactions for each targeted codon, thus significantly increasing the labor involved^20^.

Besides saturation mutagenesis, there are challenges in the development of robust and accurate functional assays. Typical functional assays used in DMS to enrich variants include growth assays and flow cytometry assays^21^. Growth assays are primarily suitable for genes significantly affecting cell viability or growth rates and may not apply to other genes. Flow cytometry assays convert gene function-related biological signals into quantifiable fluorescence. However, they are relatively costly due to cell sorter reliance, particularly when saturation mutagenesis efficiency is low thereby requiring an increased number of cells to be sorted. Second, precisely controlling protein expression levels to mimic physiological relevance poses another challenge, as overexpression of the targeted gene potentially mitigates variant pathogenic effects in certain cases^22^, thereby diminishing assay sensitivity for variant interpretation. Lastly, barcoding is a common practice in DMS studies to facilitate variant detection using short-read NGS. However, this process typically necessitates an additional round of NGS to assign the barcodes to the variants, incurring extra expenses^23^.

Addressing these challenges is crucial for applying DMS to disease-related genes. Dystroglycanopathies are a set of rare autosomal recessive diseases with clinical heterogeneity ranging from brain malformation in Walker-Warburg syndrome (WWS) to milder muscular symptoms in Limb-Girdle Muscular Dystrophies (LGMDs)^24^. Pathogenic variants in *DAG1*, the gene that encodes alpha-dystroglycan (α-DG), and genes encoding enzymes involved in α-DG glycosylation disrupt the binding between α-DG and extracellular matrix ligands, which compromises the muscle cell integrity and leads to dystroglycanopathies^25^. The most severe dystroglycanopathy cases can lead to miscarriage and neonatal deaths, highlighting the critical need for a better understanding of the clinical significance of variants in genetic testing^26–28^.

Saturation Mutagenesis-Reinforced Functional assays (SMuRF) was developed as a flexible DMS framework to address challenges that were identified in both saturation mutagenesis and functional assays. We built upon the core concept of PALS but achieved saturation mutagenesis without the need for specialized reagents, equipment, or labor-intensive steps, resulting in a method named Programmed Allelic Series with Common procedures (PALS-C). Since our method only requires minimal-length oligos, a regular plasmid template, restriction enzymes, PCR polymerase, and a Gibson assembly kit, PALS-C is ready to apply in most molecular biology laboratories. We demonstrated the versatility of SMuRF by applying it to two key enzyme genes, *FKRP* and *LARGE1*, involved in the glycosylation of α-DG. For this implementation of SMuRF, we employed lentiviral delivery to introduce variants into cells for the flow cytometric functional assay, achieving high variant coverage. In addition, we utilized the weak UbC promoter^29^, and streamlined the workflow to control gene expression levels, ensuring precise variant assessment. Lastly, we implemented a block-by-block strategy throughout the saturation mutagenesis and functional assay, eliminating the need for barcoding and significantly reducing costs.

## Results

### Establishing the functional assay in SMuRF

SMuRF was developed with the aim of creating a DMS framework that is adaptable for various genes by employing appropriate assays. Initially, the functional assay was established to characterize all possible coding single nucleotide variants (SNVs) in genes related to dystroglycanopathies. Most variants of α-DG glycosylation enzymes, including FKRP and LARGE1, lack clinical reports and those reported remain poorly interpreted (Figure S1A), with many unique to the families^30^. FKRP adds the second ribitol-5-phosphate (Rbo5P) to the Rbo5P tandem^31^ while LARGE1 is responsible for adding the repeated disaccharide units of matriglycan^32^ (Figure S1B). New drugs^33^, gene therapies^34,35^ and cell therapies^36^ are being actively developed for dystroglycanopathies related to FKRP or LARGE1, emphasizing the need for improved variant interpretation to facilitate patient enrollment in these gene-specific trials.

To develop a functional assay suitable for DMS, we initially identified α-DG hypoglycosylation as the molecular phenotype associated with dystroglycanopathies and the focal point for assessment. The IIH6C4 antibody, extensively used in α-DG-related research and clinical diagnoses, specifically binds to the matriglycan chain of glycosylated α-DG Core M3, enabling the quantification of α-DG glycosylation levels^37–50^. Fibroblasts from patients with dystroglycanopathies have been characterized using the IIH6C4 antibody in a fluorescence flow cytometry (FFC) assay^51^. The human haploid cell line, HAP1, has become a widely utilized platform in α-DG-related research, covering various areas such as dystroglycanopathy gene discovery^52^, enzymatic functions^42^, and α-DG binding properties and functions^53^. Additionally, previous studies have established the compatibility of HAP1 cells with FFC^54^. We modified the IIH6C4 FFC assay originally designed for fibroblasts to be suitable for HAP1 cells while improving its sensitivity to differentiate between variants (Figures S1C and S1D). Subsequently, this IIH6C4 assay was adapted for fluorescence-activated cell sorting (FACS) and applied to generate functional scores for all possible coding SNVs in *FKRP* and *LARGE1* (Figure 1A).

**Figure 1:**
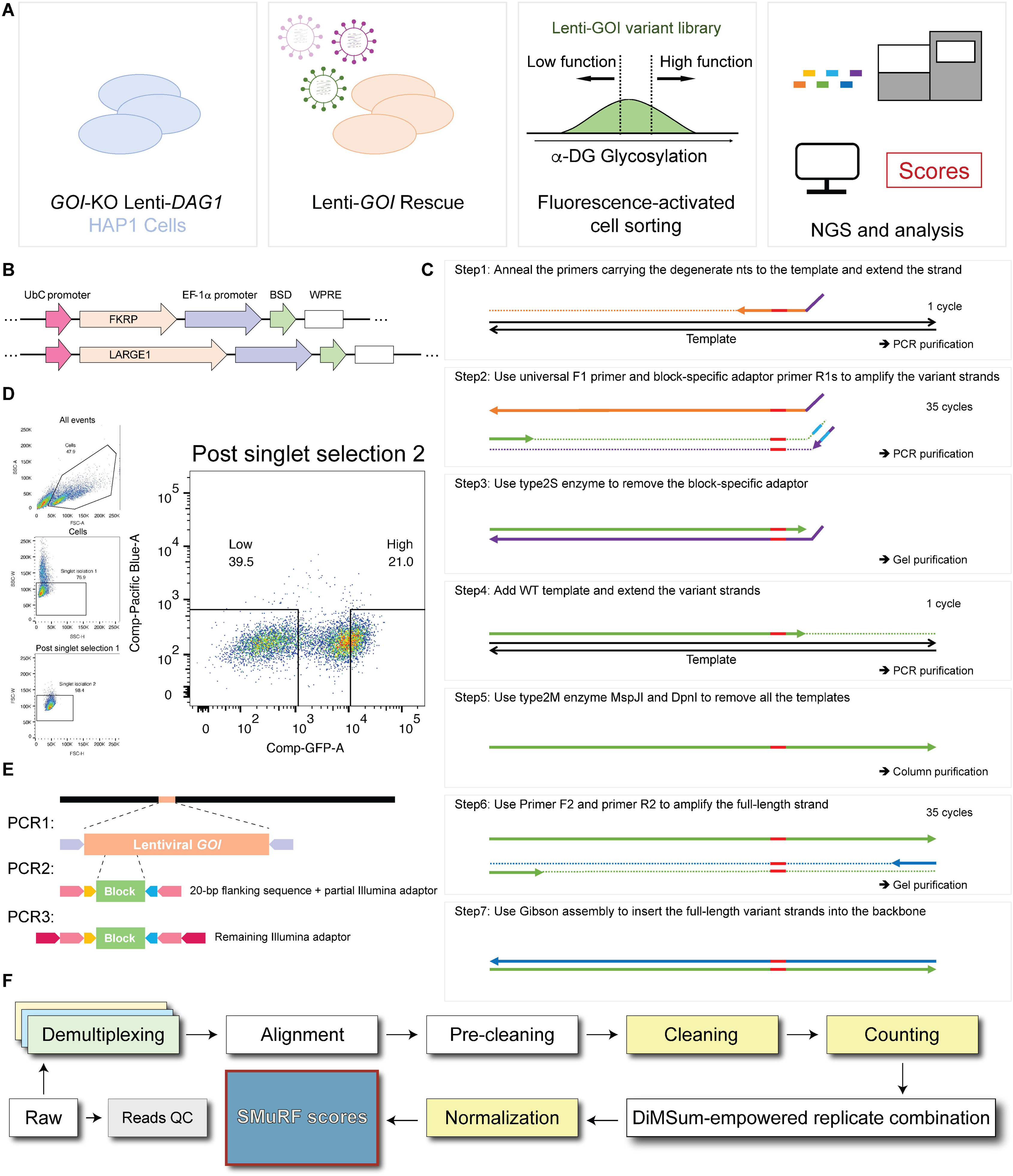
**Streamlining the saturation mutagenesis and FACS assay in SMuRF.** (A) A universal workflow of SMuRF. SMuRF accompanies saturation mutagenesis with functional assays. Here, saturation mutagenesis is achieved by delivering variant lentiviral particles to the engineered HAP1 platform where the endogenous gene of interest (GOI) was knocked out and stable *DAG1* overexpression was established through lentiviral integration. A fluorescence-activated cell sorting (FACS) assay was employed to separate the high-function population and the low-function population. (B) Lenti-*GOI* constructs used for the saturation mutagenesis. The *GOI* CDS expression is driven by a weak promoter UbC. (C) PALS-C is simple and accessible to most molecular biological laboratories. To accommodate the requirements of downstream short-read NGS, the *GOI* variants were separated into multiple blocks (6 blocks for *FKRP* and 10 blocks for *LARGE1*). PALS-C generates block-specific lentiviral plasmid pools from 1 oligo pool per GOI. The steps are massively multiplexed: Step 1 requires only a single-tube reaction; the following steps can be done in a single-tube reaction for each block. Step 8 (not shown) uses electrotransformation to deliver the assembled plasmid pools to bacteria for amplification. (D) A representative example shows the gating strategy; 20k flow cytometry events of FKRP block1 were recorded and reanalyzed with FlowJo. (E) A 3-round PCR strategy to build the NGS library. Samples from the high and low glycosylation groups were barcoded differently in PCR2. PCR2 products of all samples were multiplexed for a single PCR3 reaction. (F) A universal pipeline to generate SMuRF scores from raw NGS data. Steps colored yellow indicate employment of customized scripts. Cleaning is a critical step where the reads carrying co-occurred variants are filtered out.

### Streamlining the saturation mutagenesis and FACS assay in SMuRF

To engineer the cell line platforms for the saturation mutagenesis, the endogenous gene of interest (GOI), *FKRP* or *LARGE1*, was knocked out to make *GOI*-KO HAP1 lines, which lack the endogenous *GOI* function (Figures S1F and S1G). Since α-DG glycosylation level is the biological signal quantified by the functional assay, α-DG overexpression was achieved by Lenti-*DAG1* transduction, as a way to increase assay sensitivity. (Figures S1H-J). Subsequent experiments were performed using monoclonal *GOI*-KO Lenti-*DAG1* HAP1 lines. Since most known dystroglycanopathy cases are caused by missense variants^55^, we performed saturation mutagenesis for all possible coding SNVs. We first constructed the plasmids carrying the wild-type (WT) GOIs. To better control the expression level (Figures S1K and S1L), we employed the weak promoter UbC for GOI expression^29^, creating Lenti-UbC-*FKRP*-EF1α-*BSD* and Lenti-UbC-*LARGE1*-EF1α-*BSD* (Figure 1B).

Next, we performed PALS-C to introduce all possible coding SNVs using the WT plasmids as templates (Figure 1C). The variants were introduced initially as 64-nt reverse PCR primers. The short length of the oligos allowed for a low cost in synthesizing the oligo pool. We adopted a block-by-block strategy where we divided the GOI variants into multiple non-overlapping blocks (6 for *FKRP* and 10 for *LARGE1*). For each GOI, PALS-C initiates with one single pool of oligos and eventually generates an isolated lentiviral plasmid pool for each block. PALS-C steps are massively multiplexed: Step1 requires only a single-tube reaction; the following steps can be done in a single-tube reaction for each block (STARl1lMethods). All downstream experiments following PALS-C, up to the NGS, were conducted individually for each block. This block-by-block strategy allowed us to employ short-read NGS to examine variant enrichment while avoiding the requirement of an additional NGS to assign barcodes to variants spanning the entire CDS and the expenses associated with it^20,56,57^.

Variant representation in the lentiviral plasmid pools generated by PALS-C was evaluated using a shallow NGS service, according to which, more than 99.6% of all possible SNVs were represented (Figures S2A-D). The plasmid pools were packaged into lentiviral particles, which were subsequently delivered into the platform cells through transduction. Once transduced cells were sufficiently expanded following drug selection, they were sorted using FACS to isolate cells with either high or low glycosylation levels. Staining conditions and gating parameters in FACS were optimized with mini-libraries (Figure S2E). α-DG glycosylation level was quantified by IIH6C4-FITC signal (Figure 1D, and Figures S2F-G). The FACS events of each group achieved a minimum of ∼1000 × coverage.

Genomic DNA from each group of each block was used to build the multiplexed sequencing library using a 3-round PCR strategy (Figure 1E). Raw NGS datasets were analyzed with our customized analytical pipeline “Gargamel-Azrael” to generate SMuRF scores for all variants (Figure 1F). Essentially, the SMuRF score is the normalized relative enrichment of a variant in the FACS groups (STARl1lMethods). High SMuRF scores indicate normal functionality of variants in α-DG glycosylation while low scores indicate deleterious effects.

### SMuRF recapitulated and expanded the knowledge gained from variant databases

Three biological transduction replicates were performed for each gene to confirm the reproducibility of the workflow (Figure S2H and Table S1). After combining biological replicates using DiMSum^58^, SMuRF scores were generated for all possible coding SNVs of *FKRP* (4455) and *LARGE1* (6804) (Figures S3A and S3B, Tables S2 and S3), and those with high confidence were used for downstream analysis (*FKRP*: 4325, 97.1%; LARGE1: 6473, 95.1%). SMuRF scores align with the anticipated patterns of different variant types (Figures 2A and 2B). *FKRP* synonymous variants have a median of -0.05, with a 95% confidence interval (CI) of -0.07∼-0.02, while *LARGE1* synonymous variants have a median of -0.07 (95% CI: -0.09∼-0.04). The nonsense variants consistently exhibit low SMuRF scores, with sparse outliers observed. *FKRP* nonsense variants have a median of -2.34 (95% CI: -2.43∼-2.22), while *LARGE1* nonsense variants have a median of -2.51 (95% CI: -2.57∼-2.45). Two noteworthy outliers among the nonsense variants are *FKRP* c.1477G>T (p.Gly493Ter) (SMuRF = -0.44) and *LARGE1* c.2257G>T (p.Glu753Ter) (SMuRF = 0.15). These two are the nonsense variants positioned closest possible to their respective canonical stop codons. The relatively high SMuRF scores of these two variants suggest that their impact on the enzymatic function is negligible in the context of the CDS constructs. Furthermore, since both variants are in the last exon of their respective transcripts, it is also unlikely for them to be substantially influenced by nonsense-mediated decay (NMD) *in vivo*^59^.

**Figure 2:**
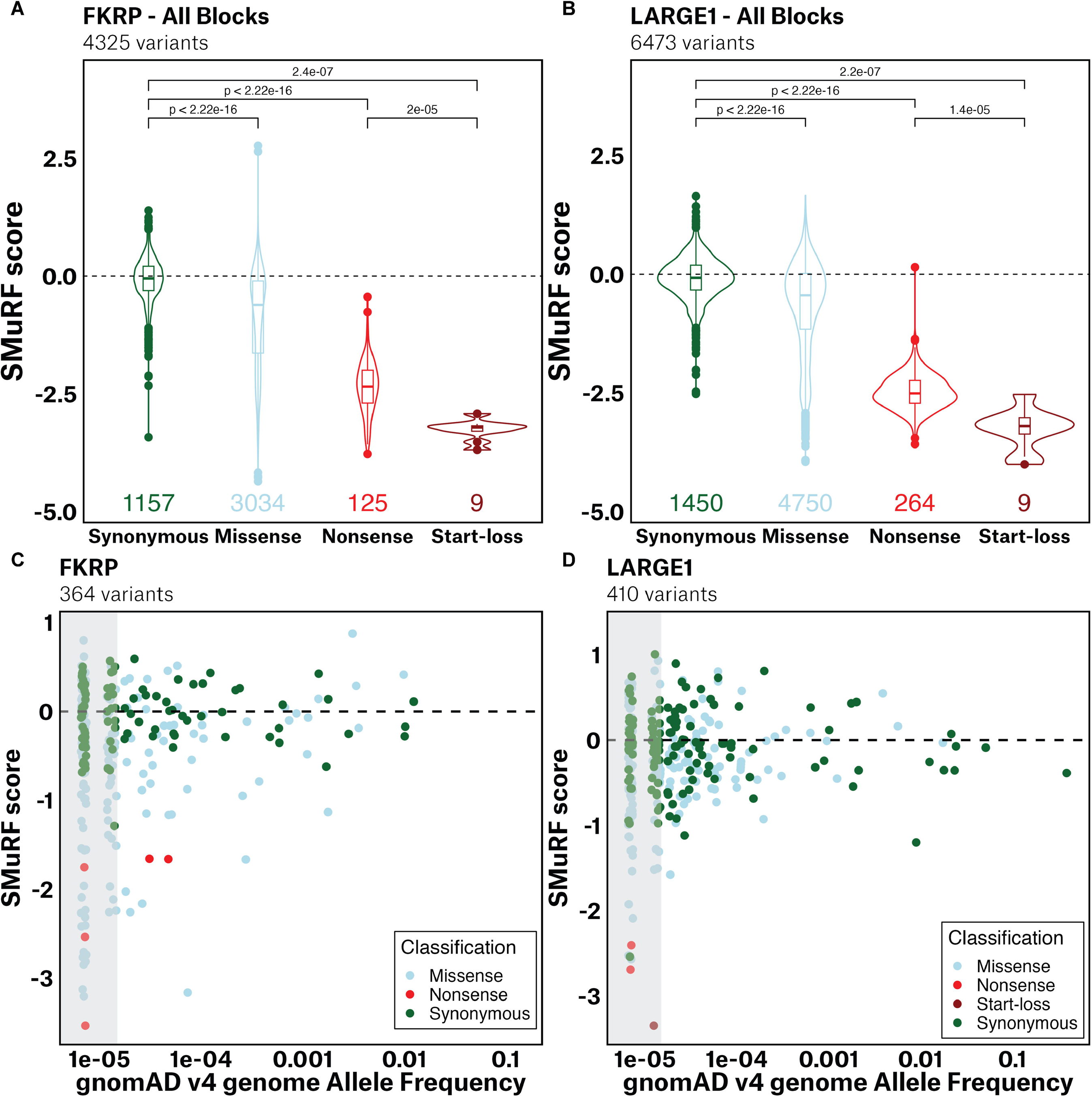
**SMuRF recapitulated and expanded the knowledge gained from variant databases.** (A and B) High confidence SMuRF scores align with variant types (A, *FKRP*; B, *LARGE1*). The mean of synonymous variants of each block was used to normalize the scores across blocks. The box boundaries represent the 25th/75th percentiles, with a horizontal line indicating the median and a vertical line marking an additional 1.5 times interquartile range (IQR) above and below the box boundaries. p-values were calculated using the two-sided Wilcoxon test. Counts of variants were labeled below the boxes. (C and D) SMuRF revealed functional constraints based on variants reported in gnomAD v4.0.0 genome sequencing data (C, *FKRP*; D, *LARGE1*): Low allele frequency variants had diverse functional scores, while high allele frequency variants converged towards wild-type (WT) due to selection pressures (Gray box: AF < 1.5e-05). Dashed lines represent WT functional score. Dots were jittered with geom_jitter (width = 0.05, height = 0.05).

Interestingly, the start-loss variants exhibit markedly low SMuRF scores, significantly lower than those observed in most nonsense variants (p-value = 2e-5, *FKRP*; 1.4e-5, *LARGE1*). *FKRP* start-loss variants have a median of -3.21 (95% CI: -3.52∼-3.14), while *LARGE1* start-loss variants have a median of -3.19 (95% CI: -3.84∼-2.99). While this observation does not necessarily indicate stronger negative effects of the start-loss variants, compared to that of the nonsense variants, it indicates that, at least in the context of the *FKRP* and *LARGE1* SMuRF CDS constructs, there is a lack of effective genetic compensation to counter the start-loss variants, such as functional downstream alternative start codons^60^. The homozygous start-loss variant *FKRP* c.1A>G (SMuRF = -3.29) has been reported to be associated with WWS, the most severe FKRP-related disorder. This variant has been documented in two cases, with one resulting in the unfortunate death of a child at the age of 6 days and the other leading to a terminated pregnancy^28^. Therefore, the start-loss variants may warrant increased attention in genetic testing protocols.

When compared to allele frequency (AF) data in The Genome Aggregation Database (gnomAD, v4.0.0)^6,61^, SMuRF scores aligned with the selection against pathogenic variants: low allele frequency variants (AF < 1.5e-05, representing allele count of 1 or 2 in gnomAD v4 genome sequencing data), exhibited a wide range of functional scores, while variants with higher frequency showed functional scores converging towards the WT score as common variants are depleted of pathogenic variants (Figures 2C, 2D, S3C and S3D).

The α-DG glycoepitope, as well as the enzymes involved in its glycosylation, are largely conserved within Metazoa^62^. Evolutionary conservation scores, such as PhyloP scores, indicate the degree of conservation of a variant derived from multiple sequence alignments across species^63^. When compared with PhyloP scores calculated from 100 vertebrates, SMuRF demonstrated the evolutionary tolerance of relatively harmless variants and the selection against damaging variants in both *FKRP* and *LARGE1* (Figures S3E and S3F), with a tendency for missense variants to be more disruptive at the more conserved sites (rho = -0.44, *FKRP*; -0.25, *LARGE1*).

### SMuRF is sensitive and accurate in predicting variant pathogenicity and severity

ClinVar is a public archive of reports of human variants^64^, where the variants were classified according to clinical supporting evidence into different categories including Benign (B), Benign/likely benign (B/LB), Likely benign (LB), Likely pathogenic (LP), Pathogenic/likely pathogenic (P/LP), Pathogenic (P), and Variants of Uncertain Significance (VUS). SMuRF scores correlate well with clinical classification in ClinVar (Figures 3A, 3B, S3G, and S3H). Furthermore, dystroglycanopathies associated with *FKRP* encompass a spectrum of diseases with varying severity, including severe cases like WWS and muscle-eye-brain disease (MEB), intermediate cases like congenital muscular dystrophies (CMD), and relatively mild cases like LGMDR9 (LGMD2I)^24,28^. To explore whether SMuRF scores reflect the disease severity, we employed a naive additive model where the functional scores of the variants on both alleles were summed to calculate the biallelic functional score^65,66^. We aggregated data from 8 well-curated cohorts and compared them with SMuRF scores (Table S4)^28,47,55,67–71^. The combined functional scores of the mild cases were significantly higher compared to those of the intermediate and severe cases, as expected (Figure 3C). Another strategy to evaluate the biallelic variant effect is using the higher allelic score to represent the compound heterozygous variant pair^56^. This model produces similar results and conclusions as the naive additive model (Figure S3I). Additionally, SMuRF scores showed a correlation with the reported disease onset age (Figure 3D), where high-function variants were associated with later onset (rho = 0.72; 0.70, male; 0.73, female). Cox proportional hazards analysis revealed a statistically significant association between the combined SMuRF scores and the onset (p-value < 0.003). The estimated hazard ratio was 0.28 (95% CI: 0.12 - 0.64), indicating that higher functional scores are correlated with a substantially lower risk of earlier disease onset. Specifically, the negative coefficient (-1.27) suggests that as the functional score decreases, the likelihood of early onset increases. The model’s concordance index of 0.67 reflects a moderate level of predictive accuracy.

**Figure 3:**
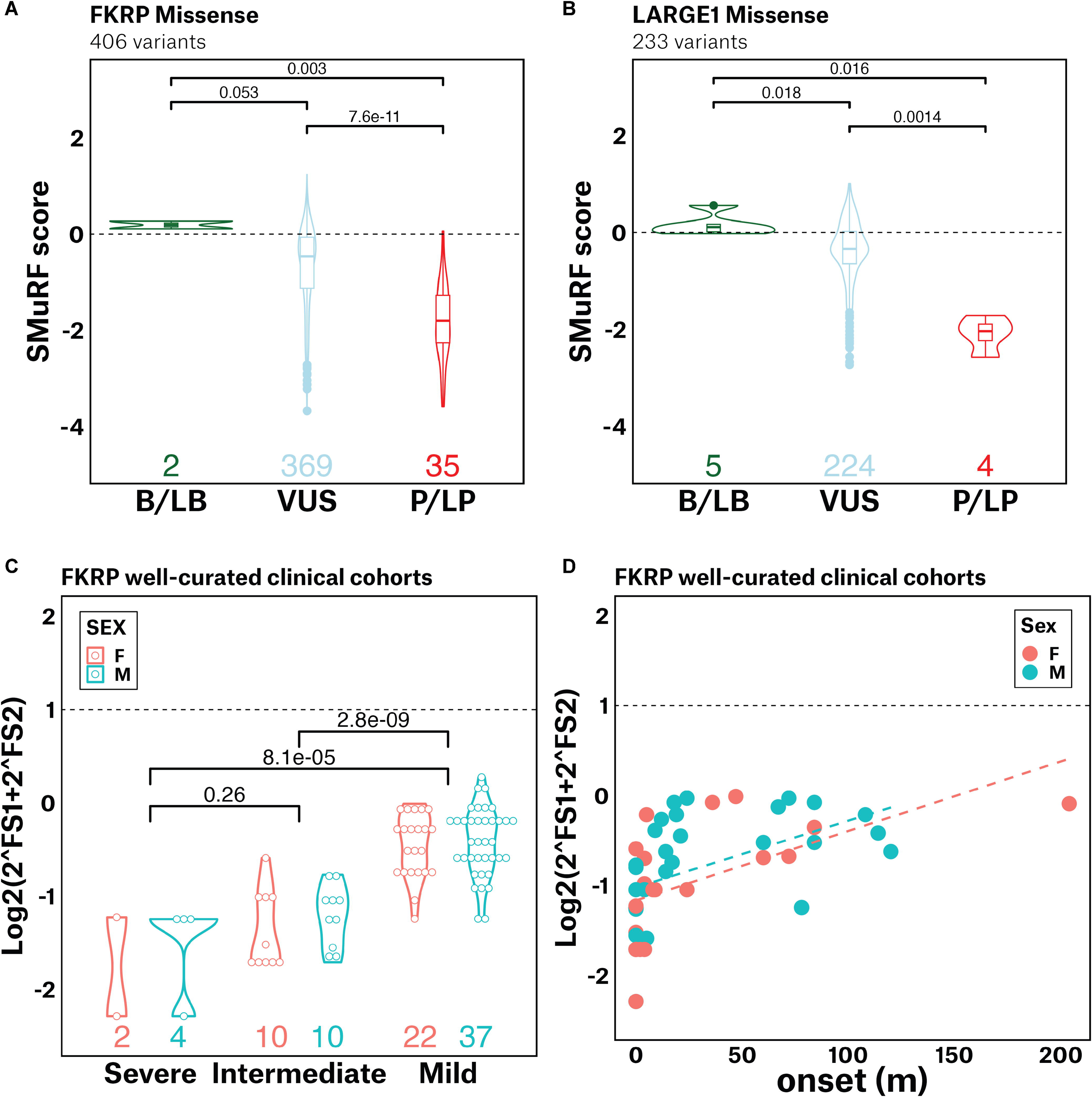
**SMuRF improved the scope of clinical interpretation of genetic variants.** (A and B) SMuRF scores correlate well with clinical classification in ClinVar (**A,** *FKRP*; **B,** *LARGE1*). (B/LB: Benign, Benign/Likely benign or Likely benign in ClinVar; VUS: Uncertain significance in ClinVar; P/LP: Pathogenic, Pathogenic/Likely pathogenic or Likely pathogenic in ClinVar.) Counts of variants were labeled below the violins. (C) Real patient data from eight well-curated cohorts demonstrated that SMuRF scores have the potential to predict disease severity. The additive SMuRF scores of the variant pairs associated with mild cases were significantly higher than those of intermediate and severe cases. Counts of cases were labeled below the violins. p-values were calculated using the Wilcoxon test. FS1, the SMuRF functional score of the variant on Allele1; FS2, the SMuRF functional score of the variant on Allele2. (D) The SMuRF scores are correlated with the disease onset age. Dashed trendlines represent linear regression. Spearman’s rank correlation rho: 0.72 (all data), 0.70 (male), 0.73 (female).

These analyses indicate the potential utility of the SMuRF scores for providing another line of evidence for clinical variant interpretation. According to the standards and guidelines developed jointly by the American College of Medical Genetics (ACMG) and the Association for Molecular Pathology (AMP), the SMuRF scores can potentially be applied as functional evidence under the PS3/BS3 criterion. This criterion pertains to "well-established" functional assays that demonstrate whether a variant has abnormal or normal gene/protein function, respectively^72,73^. Therefore, we employed the SMuRF scores to label the *FKRP* variants as SMuRF Benign, SMuRF Mild, SMuRF Intermediate and SMuRF severe, and *LARGE1* variants as SMuRF Benign and SMuRF Pathogenic (Tables 1, S2, S3 and STARl1lMethods). It is important to note that clinical variant interpretation involves the application of various criteria, and functional data such as SMuRF scores are just one piece of the contributing evidence, which is not intended to be used as a decisive classification in isolation. For *FKRP*, the SMuRF labeling was performed based on 3 boundary scores. The boundary score between SMuRF Benign and SMuRF Mild (-0.87) was determined according to the inflection point between the ClinVar P/LP and ClinVar B/LB densities. The boundary score between SMuRF Mild and SMuRF Intermediate (-1.62) was determined according to the peak value of the mild cases from the 8 well-curated cohorts. The boundary score between SMuRF Intermediate and SMuRF Severe (-2.24) was determined according to the peak value of the severe cases from the 8 well-curated cohorts. For *LARGE1*, the SMuRF labeling was performed based on 1 boundary score. The boundary score between SMuRF Benign and SMuRF Pathogenic (-1.32) was determined according to the inflection point between the ClinVar P/LP and ClinVar B/LB densities.

**Table 1.**
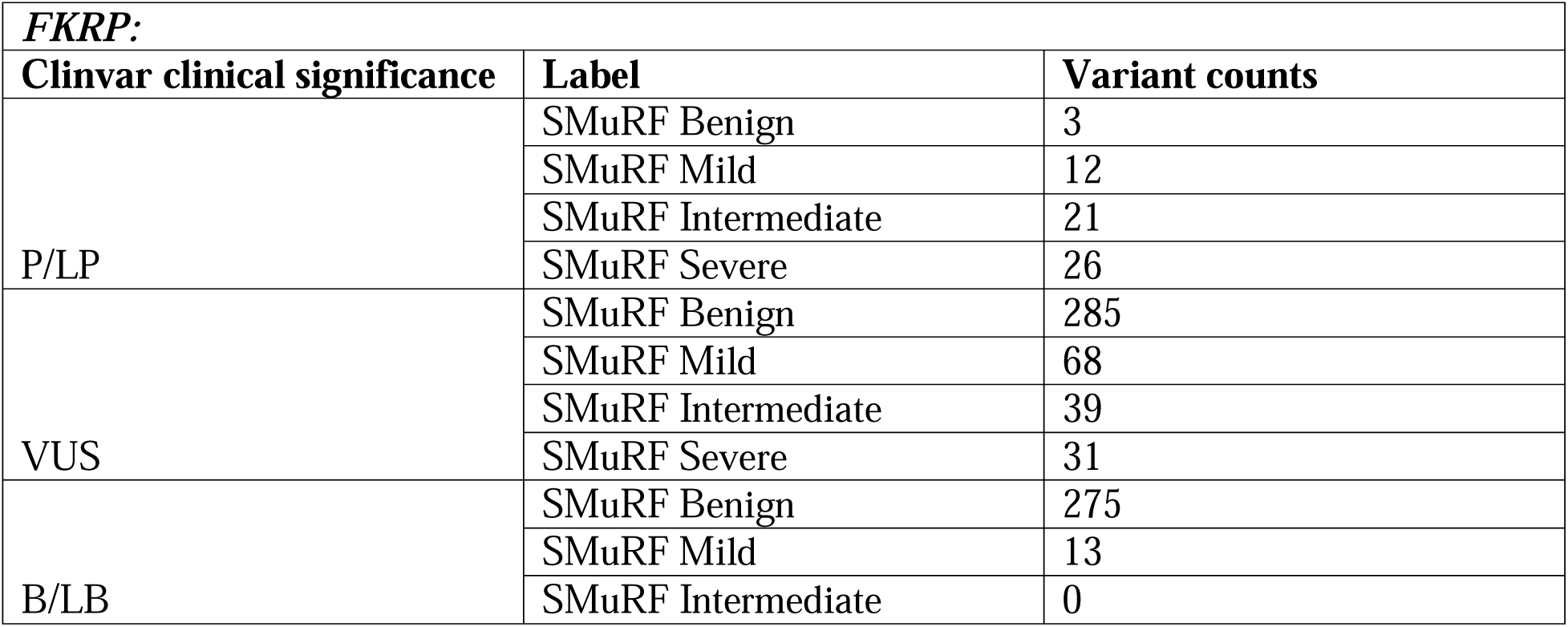
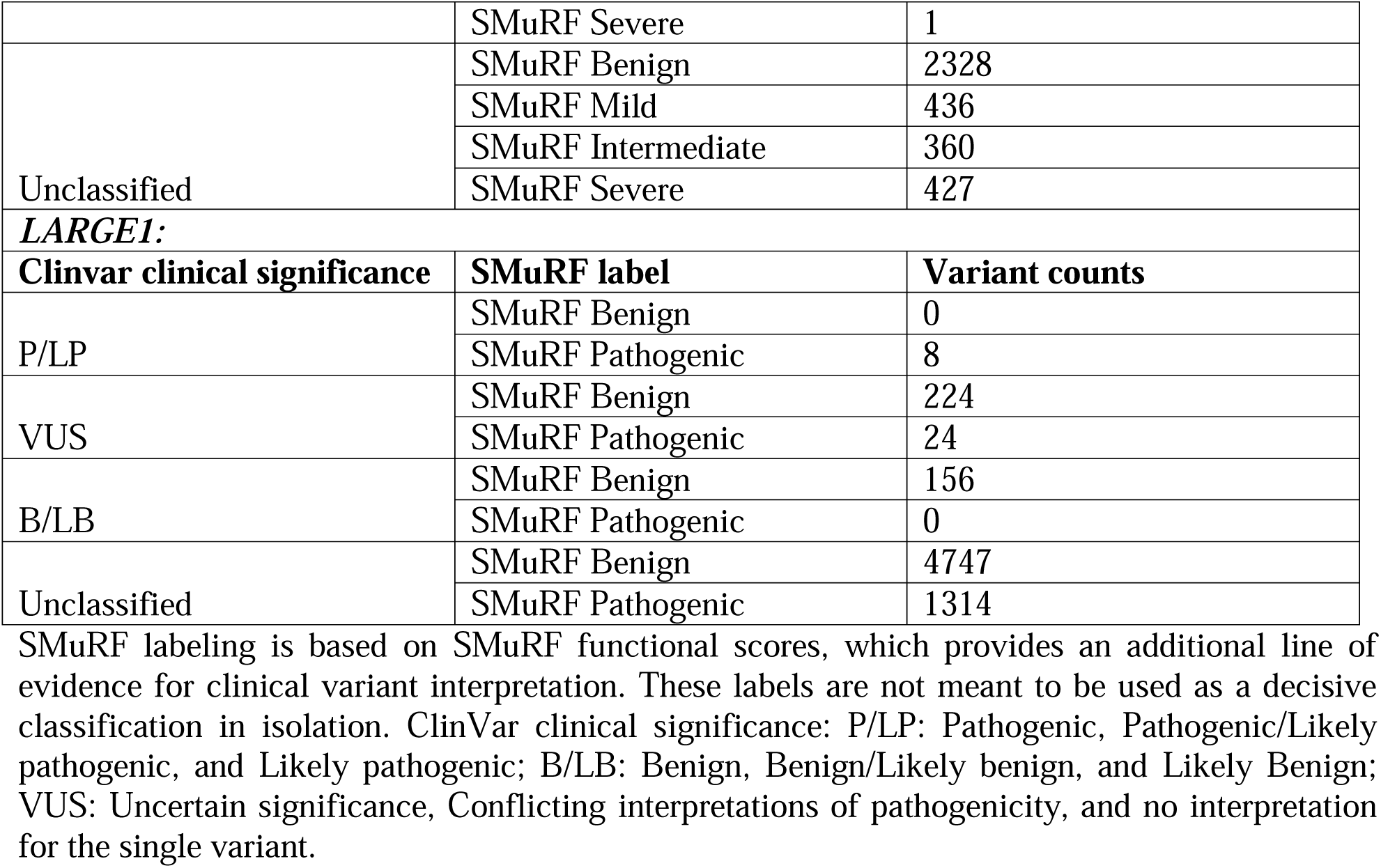
A potential SMuRF labeling.

### SMuRF scores can be employed to validate and improve computational predictors

In addition to directly assisting in variant classification, SMuRF scores can also be used to validate and improve computational predictors. Computational prediction is currently an active area of research with a wealth of methods recently developed, including CADD^74^, metaSVM^75^, REVEL^76^, MVP^77^, EVE^78^, MutScore^79^, PrimateAI-3D^80^, ESM1b^81^, MAVERICK^82^, and AlphaMissense^83,84^. To compare SMuRF with these computational predictors, we assessed the receiver operating characteristic (ROC) curves for all methods using the P, P/LP, and LP variants in ClinVar as the positive reference set. For the negative reference set, we used missense variants observed in the gnomAD v4 genome sequencing database. Variants present in both ClinVar P, P/LP, LP and gnomAD v4 were excluded from the negative reference set. A higher Area Under Curve (AUC) value indicates better discriminatory ability in classifying pathogenic variants (Figures 4A, 4B and Table S5). SMuRF outperforms all computational methods for *LARGE1* (AUC = 0.96), followed by REVEL (AUC = 0.88). For *FKRP*, two predictors, REVEL (AUC = 0.88) and EVE (AUC = 0.86) exhibit comparable performance to SMuRF (AUC = 0.86). It is noted that REVEL was trained with Human Gene Mutation Database (HGMD) pathogenic variants, which have an overlap with ClinVar variants, potentially contributing to its higher AUC when evaluated on the ClinVar dataset. We checked the correlation between the predictors and SMuRF. Most predictors assigned higher scores to pathogenic variants, and hence are negatively correlated with SMuRF scores (except ESM1b). Among all the predictors examined, AlphaMissense has the strongest correlation with SMuRF (rho = -0.70, *FKRP*; -0.54, *LARGE1*) (Figures 4C-E, S4A and Table S6). REVEL, among the predictors, demonstrates the best performance according to the ROC curves and also exhibits a relatively good correlation with SMuRF (rho = -0.65, *FKRP*; -0.47, *LARGE1*) (Figure S4B).

**Figure 4:**
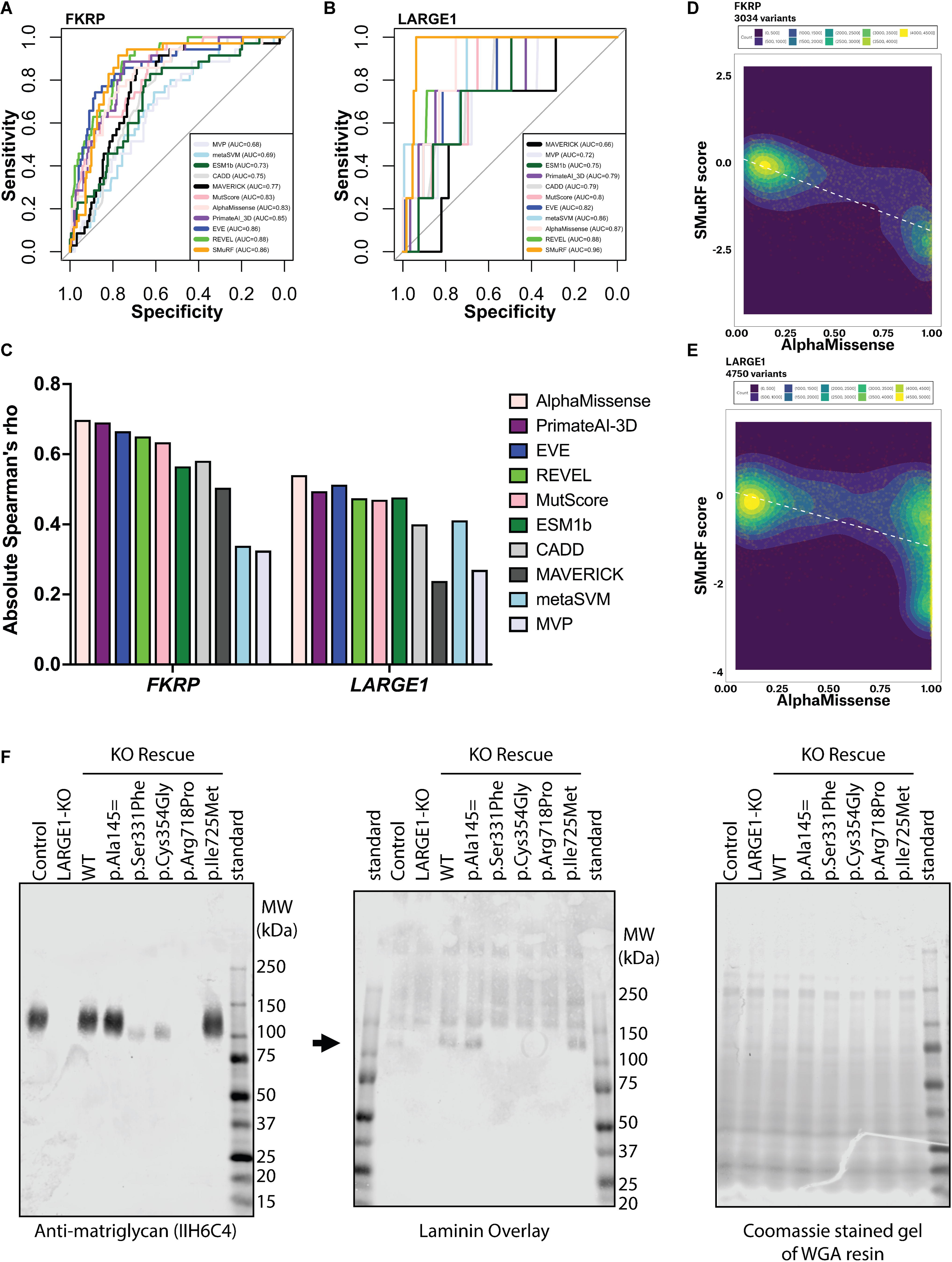
**SMuRF scores can be employed to validate and improve computational predictors.** (A and B) Receiver operating characteristic (ROC) curves of SMuRF and computational predictors (A, *FKRP*; B, *LARGE1*). AUC: Area Under Curve. Higher AUC indicates better performance in classifying pathogenic variants. (C) The correlation coefficient was calculated between SMuRF scores and scores generated by computational predictors. Figure depicts absolute correlation coefficient. (D and E) Among all the predictors examined, AlphaMissense has the strongest correlation with SMuRF (rho = -0.70, *FKRP*; -0.54, *LARGE1*). Density was calculated with contour_var = “count” in R. White dashed lines represent linear regression. (F) IIH6C4 blots indicate matriglycan synthesis activity of selected LARGE1 enzyme variants. The laminin overlay provides a different probe for matriglycan, with an arrow indicating the expected band size. The protein loading was controlled by Coomassie gel. Experiments were conducted with the myogenic cell line, MB135. Control: WT MB135. KO Rescue: endogenous *LARGE1* was knocked out and the cells were resecured with individual lentiviral transduction. P.Ala145= (SMuRF = -0.43) has the highest AF (0.41) in gnomAD v4, which was used as a high-function reference. P.Ser331Phe (SMuRF = -2.58) is Pathogenic in ClinVar, which was used as a low-function reference.

To explore the potential benefit of combining experimental and predicted scores for more accurate variant interpretation, we selected three *LARGE1* VUSs from ClinVar with concordant designations by both SMuRF and AlphaMissense: c.2175C>G p.Ile725Met (concordant likely benign; SMuRF = 0.43; AlphaMissense = 0.0733), c.1060T>G p.Cys354Gly (concordant likely pathogenic; -2.03; 0.9962), c.2153G>C p.Arg718Pro (concordant likely pathogenic; -2.56; 0.9995). Anti-matriglycan Western blot and laminin overlay results confirmed the expected glycosylation activity of these 3 variants, along with the variants with decisive clinical classification (Figure 4F). This approach showcases the strategy for enhancing variant interpretation by corroborating DMS scores with prediction scores.

### SMuRF highlighted the critical structural regions

The currently known disease-related mutations in *FKRP* and *LARGE1* are distributed throughout their entire sequences (Figure S5A), and only limited critical structural sites have been identified and associated with specific disease mechanisms. SMuRF can contribute to highlighting critical structural regions in the enzymes. The protein structures of both FKRP and LARGE1 have been previously studied. FKRP is known to have a stem domain (p.1-288) and a catalytic domain (p.289-495)^85^. SMuRF scores revealed that missense variants in the catalytic domain are generally more disruptive than those in the stem domain (p-values < 2.22e-16) (Figure 5A and 5B). Furthermore, it has been reported that a zinc finger loop within the catalytic domain (p.289-318) plays a crucial role in FKRP enzymatic function^85^. SMuRF analysis demonstrated that missense variants in the zinc finger loop (SMuRF median: -1.50; 95% CI: -2.00∼-1.10) exhibit greater disruption, compared to variants in the remaining region of the catalytic domain (SMuRF median: -1.19; 95% CI: -1.29∼-1.08), though not statistically significant (p-value = 0.17).

**Figure 5:**
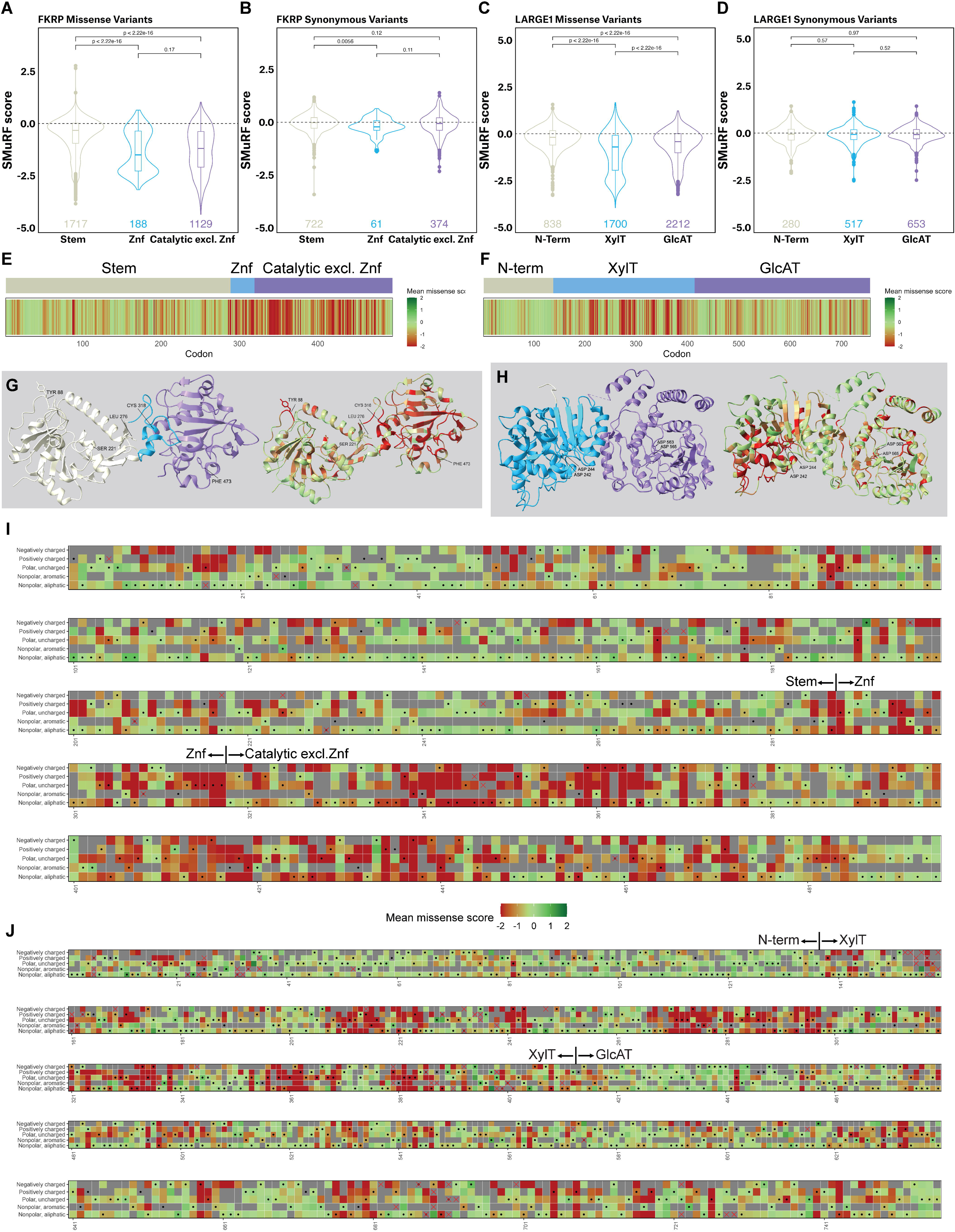
**SMuRF highlights the critical structural regions.** (A) SMuRF scores showed higher functional disruption by missense variants in the catalytic domain of *FKRP* compared to the stem domain. The zinc finger loop (Znf) within the catalytic domain exhibited greater disruption by missense variants. Box plots depict the 25th/75th percentiles (box boundaries), median (horizontal line), and an additional 1.5 times IQR (vertical line) above and below the box boundaries. p-values were calculated using the two-sided Wilcoxon test. Counts of variants were labeled below the violins. Dashed lines represent WT functional score. (B) The SMuRF scores of synonymous *FKRP* variants in different domains. (C) Missense variants in the catalytic domains of *LARGE1* showed higher disruption compared to the N-terminal domain. Missense variants in the XylT domain were more disruptive than those in the GlcAT domain. (D) The SMuRF scores of synonymous *LARGE1* variants in different domains. (E and F) Mean SMuRF scores were utilized to map SNV-generated single amino acid substitutions onto the 1D structures of the enzymes (E, *FKRP*; F, *LARGE1*). The mean SMuRF score per amino acid residue was calculated and visualized using a color scale, where red indicates positions sensitive to substitutions and green is tolerated. (G and H) Mean SMuRF scores were utilized to map SNV-generated single amino acid substitutions onto the 3D structures of the enzymes (G, *FKRP*; H, *LARGE1*). The crystal structure of human FKRP (PDB:6KAM, codon: 45-495) and the electron microscopy structure of LARGE1 (PDB:7UI7, codon: 34-756) were used. Same color scale is employed as E and F. (I and J) Heatmap representation of the mean SMuRF scores for each codon (I, *FKRP*; J, *LARGE1*). Amino acids were categorized into five groups: nonpolar, aliphatic (G, A, V, L, M, I); polar, uncharged (S, T, C, P, N, Q); positively charged (K, R, H); negatively charged (D, E); and nonpolar, aromatic (F, Y, W). Each cell in the heatmap corresponds to a codon position (x-axis) and an amino acid group (y-axis). The black dots indicate the wild-type amino acid group for each residue. Grey squares denote the scenario where the amino acid change is not possible with a single SNV within the codon, and a red cross marks positions where variants were filtered out due to low confidence.

LARGE1 has two catalytic domains: a xylose transferase (XylT) domain (p.138-413) and a glucuronate transferase (GlcAT) domain (p.414-756)^86^. They are each responsible for adding one unit of the polysaccharide matriglycan chain, which consists of alternating xylose and glucuronate units. SMuRF revealed that the missense variants in both catalytic domains tend to be significantly more disruptive than the variants in the N-terminal domain (p-values < 2.22e-16) (Figure 5C and 5D). Interestingly, SMuRF also showed that the missense variants in the XylT domain (SMuRF median: -0.70; 95% CI: -0.79∼-0.62) tend to be more disruptive than those in the GlcAT domain (SMuRF median: -0.42; 95% CI: -0.45∼-0.38) (p-value < 2.22e-16). A previous IIH6C4 western blot experiment revealed a similar observation, demonstrating that mutations deactivating the GlcAT domain, but not the XylT domain, can generate a faint band indicative of glycosylated matriglycan^86^. Moreover, AlphaMissense also presents the same effects, where missense variants in the XylT domain tend to be more disruptive (Figure S4C and Supplemental Note).

To explore the variants’ effects on different structural regions, we first mapped the mean SMuRF scores of SNV-generated single amino acid substitutions onto the 1D structures of the enzymes (Figures 5E and 5F). The heatmaps highlight the functional importance of the catalytical domains. We further mapped the same mean SMuRF scores onto the 3D structures of the enzymes (Figures 5G and 5H) (FKRP: PDB 6KAM; LARGE1: PDB 7UI7), thereby highlighting the critical regions that are susceptible to missense disruptions. SMuRF confirmed the functional importance of p.Cys318 in FKRP (mean SMuRF = -2.15), which is required for Zn2+ binding in the zinc finger loop^87^. The p.Cys318Tyr variant (SMuRF = -2.23) has been reported to be associated with WWS^70^. SMuRF also highlighted the functional importance of p.Phe473 in FKRP (mean SMuRF = -1.99), which is located in a small hydrophobic pocket essential for CDP-ribitol substrate binding within the catalytic domain^87^. Three important amino acids in the FKRP stem domain were labeled on the 3D structure as well: p.Tyr88 (mean SMuRF = -2.69) and p.Ser221 (mean SMuRF = -1.00), which are both situated at the subunit-subunit interface involved in FKRP tetramerization *in vivo* (Figure S5B), and p.Leu276 (mean SMuRF = -0.61), which interacts with the catalytic domain^85^. p.Tyr88Phe is likely associated with disease^88^, and has a low SMuRF score (-3.26). p.Ser221Arg was associated with CMD-MR (MR: mental retardation)^89^. All three p.Ser221Arg SNVs have low SMuRF scores (c.661A>C: -2.15; c.663C>A: -2.21; c.663C>G: -2.17). Moreover, c.663C>A was examined in the mini-library screen and presented low function (Figure S2E). p.Leu276Ile is a founder mutation in the European population^90^, which is commonly associated with milder symptoms^91^. Interestingly, it has a relatively higher SMuRF score (-0.44) and performed more similarly to the benign variants rather than other pathogenic variants in the mini-library screen (Figure S2E). In addition, SMuRF highlighted the importance of p.Asp242 (mean SMuRF = -2.61) and p.Asp244 (mean SMuRF = -2.34) in LARGE1, which are crucial for XylT activity, as well as p.Asp563 (mean SMuRF = -1.18) and p.Asp565 (mean SMuRF = -1.57), which are required for GlcAT activity^86^. Variants affecting different enzyme domains may require distinct treatment approaches^92^. SMuRF can assist in selecting appropriate treatments for different variants by highlighting critical regions in different domains.

Lastly, we separated the amino acids into five groups: “negatively charged”, “positively charged”, “polar, uncharged”, “non-polar, aromatic”, and “non-polar, aliphatic”. Subsequently, the mean SMuRF scores of the SNV-induced substitutions were plotted for each residue according to their corresponding group (Figure 5I and 5J). Together, these figures provide an intuitive grasp of the patterns of constraint throughout the primary sequence and potentially aid in helping interpret variants in critical regions such as the catalytical domains and the signal anchor regions (*e.g.*, p.1-44 of FKRP)^93,94^.

### Validations confirmed SMuRF findings in the myogenic context

One caveat of SMuRF is that the HAP1 platform cell line, although widely used in α-DG-related studies, may not fully reflect the clinical relevance of dystroglycanopathies, which primarily affect neuromuscular tissues^95^. To address this issue, we generated myogenic platform cell lines by engineering MB135, a human control myoblast cell line^96^. Endogenous *FKRP* or *LARGE1* were knocked out respectively in the MB135 cell line. Monoclonal homozygous KO lines were established for both genes (Figure S6A). Despite being incompatible with the flow cytometric assay (Figure S6B), the KO MB135 myoblasts were effectively utilized for individual variant validation using an immunofluorescence (IF) assay that we developed. The *FKRP*-KO and *LARGE1*-KO MB135 myoblasts were rescued by different individual variants using lentivirus and differentiated into myotubes for IIH6C4 IF staining (Figures 6A, 6B and S6C). The results were consistent with the SMuRF scores, the mini-library screen (Figure S2E) and the ClinVar reports.

**Figure 6:**
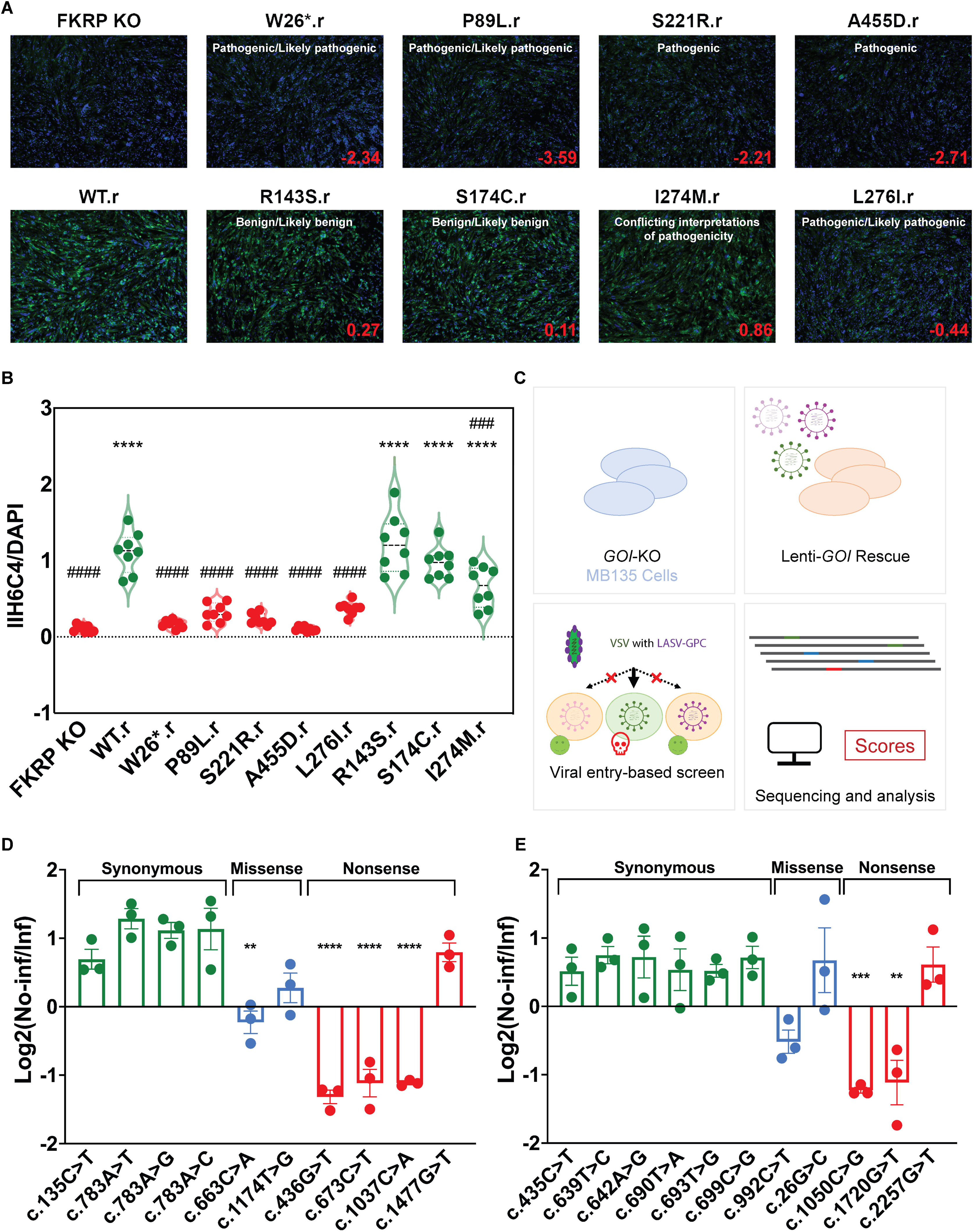
**Validations confirmed SMuRF findings in the myogenic context.** (A) Validation of individual *FKRP* variants using an IIH6C4 IF assay. The myoblasts underwent transduction and drug selection, followed by differentiation into myotubes, which were subsequently used for IF. “.r” denotes lentiviral transduction of an individual variant. Blue: DAPI. Green: IIH6C4, the glycosylation level of α-DG. Red: SMuRF scores; White: ClinVar clinical significance. The brightness and contrast of the photos were adjusted in Adobe Photoshop with the same settings. (B) Immunofluorescence intensity was quantified using integrated density (IntDen) of IIH6C4 relative to DAPI in differentiated myotubes by ImageJ. Analyses were conducted on 8 representative images. *p < 0.05, ****p < 0.0001 (compared with FKRP KO group). ##p < 0.01, ###p < 0.001, ####p < 0.0001 (compared with WT.r group). Multiple comparisons between groups were performed using analysis of variance (ANOVA) followed by Bonferroni post hoc test through GraphPad Prism 10.2.2. Experiments were conducted independently three times. (C) An orthogonal assay based on α-DG-dependent viral entry. Vesicular stomatitis virus (VSV) with Lassa fever virus glycoprotein complex (LASV-GPC) can infect cells in a glycosylated-α-DG-dependent manner. Variant enrichment before/after VSV infection can be used to quantify their performances regarding α-DG glycosylation. (D and E) The ppVSV assay can be employed to validate the findings from the flow cytometry assay (D, *FKRP*; E, *LARGE1*). 10 Lenti-*FKRP* variants were mixed with Lenti-WT-*FKRP* to rescue *FKRP*-KO MB135. 11 Lenti-*LARGE1* variants were mixed with Lenti-WT-*LARGE1* to rescue *LARGE1*-KO MB135. The functional score was quantified by the ratio of a variant’s enrichment in the non-infected group to its enrichment in the ppVSV-infected group. A higher functional score indicates better performance in α-DG glycosylation. *FKRP* c.135C>T (p.Ala45=) and *LARGE1* c.435C>T (p.Ala145=) have the highest AFs in gnomAD v4. *FKRP* c.663C>A (p.Ser221Arg) and *LARGE1* c.992C>T (p.Ser331Phe) are Pathogenic in ClinVar. Biological replicate N = 3: Lentiviral transduction and ppVSV infection were both performed independently. Figures display mean values with SEM. Additional discussions for the ppVSV results are included in Supplemental Note. Multiple comparisons were performed using ANOVA and Dunnett’s test.

To further validate the SMuRF scores, we developed an orthogonal assay that is independent of the IIH6C4 antibody to examine α-DG glycosylation level. Proper glycosylation of α-DG is crucial for the viral entry of Lassa fever virus (LASV) and its utility as an assay to study dystroglycanopathies has been demonstrated in previous studies^97^. LASV glycoprotein complex (LASV-GPC) has been employed to generate recombinant vesicular stomatitis virus (rVSV-LASV-GPC) as a safer agent for investigating LASV entry^98,99^. rVSV-LASV-GPC was utilized in a previous gene-trap screen in HAP1 cells to identify crucial genes involved in α-DG glycosylation, where cells with dysfunctional α-DG glycosylation genes exhibited increased resistance to rVSV-LASV-GPC infection, resulting in their enrichment in the population^52^. We increased the sensitivity of this VSV assay and employed it to validate interesting findings from the flow-cytometry assay (Figure 6C). Instead of using rVSV-LASV-GPC, whose genome contains the LASV-GPC coding sequence, allowing the virus to re-enter cells, we utilized pseudotyped ppVSV-LASV-GPC. The ppVSV cannot re-enter cells as it is pseudotyped using a LASV-GPC helper plasmid and lacks a viral glycoprotein coding sequence in its genome (STARl1lMethods). The *GOI*-KO MB135 myoblasts were first transduced with Lenti-*GOI* variants and selected with blasticidin. Subsequently, half of the cells were infected by the ppVSV-LASV-GPC, while the other half was kept as the non-infected control.

We mixed 10 Lenti-*FKRP* variants with Lenti-WT-*FKRP* to rescue *FKRP*-KO MB135 (Figure 6D), and mixed 11 Lenti-*LARGE1* variants with Lenti-WT-*LARGE1* to rescue *LARGE1*-KO MB135 (Figure 6E). *FKRP* c.135C>T (p.Ala45=; SMuRF = 0.17; AF = 0.14) and *LARGE1* c.435C>T (p.Ala145=; SMuRF = -0.43; AF = 0.41) have the highest AFs in gnomAD v4, which were used as high-function references. *FKRP* c.663C>A (p.Ser221Arg; SMuRF = -2.21) and *LARGE1* c.992C>T (p.Ser331Phe; SMuRF = -2.58) are Pathogenic in ClinVar, which were used as low-function references. Nonsense variants with high SMuRF scores, as well as synonymous outliers with either low or high scores, were included to be validated (inclusion criteria: Supplemental Method). The ppVSV assay supports the notion that the 3’-most nonsense variants, *FKRP* c.1477G>T (p.Gly493Ter; SMuRF = -0.44) and *LARGE1* c.2257G>T (p.Glu753Ter; SMuRF = 0.15), can retain the enzymatic functions, while other nonsense variants (including those with 2^nd^ highest SMuRF scores for both genes) did not exhibit high functionality in the ppVSV assay. With the successful application of the ppVSV assay in small-scale variant interpretation experiments, we have laid the foundation for optimizing the conditions to make the assay applicable for large-scale, all-possible variant interpretation experiments (Supplemental Note).

## Discussion

In this study, we introduced SMuRF as a framework for DMS studies and demonstrated its utility in improving variant interpretation in genetic diseases. We developed an accessible saturation mutagenesis method and functional assays with the sensitivity and accuracy required by DMS studies. We generated high-confidence functional scores for over 97% and 95% of all possible coding SNVs of *FKRP* and *LARGE1*, respectively. The SMuRF scores enable severity prediction, which was validated with well-curated patient reports. SMuRF also resolved critical protein structural regions susceptible to missense disruptions, aiding in variant interpretation.

The SMuRF framework builds upon prior DMS approaches but distinguishes itself by prioritizing accessibility and simplicity. The PALS-C method of saturation mutagenesis can be performed in a pooled fashion using common reagents and an unmodified plasmid template. In contrast, other similar methods either require a large number of individual reactions^20^, or require a specialized template (*e.g.*, uracil-containing single-stranded DNA)^19,56,100,101^. Our method has avoided these needs by optimizing the cloning steps. In addition, we employed a block-by-block approach of introducing variants and then performing sorting on each block. This approach avoids assigning barcodes for each variant and the expenses associated with it. Lastly, we have made detailed protocols and code available for designing and cloning of libraries and downstream data processing, which can be non-trivial barriers of entry for researchers.

Our study highlighted the need to control expression levels when studying enzymes. In developing our assay, we observed that over-expression of a *FKRP* mutant can compensate for its enzyme deficiency. We used the weaker UbC promoter, which is better correlated with endogenous expression and avoids this compensation. Overcoming these expression challenges in *FKRP* and *LARGE1*, SMuRF can be readily adapted for studying other enzymes involved in α-DG Core M3 glycosylation (Figure S1B). Additionally, variants in POMGNT1, an enzyme that mainly participates in the glycosylation of Core M1 and M2, can perturb the IIH6C4 signal^37^ (Figure S1B), potentially by serving as an “enzymatic chaperone” for FKTN^62,102–104^. However, variants in GnT-Vb/IX, an enzyme that participates in Core M2 glycosylation, are unlikely to alter IIH6C4 signal^105^. In summary, SMuRF is readily applicable to at least 12 other dystroglycanopathy enzymes, and the PALS-C saturation mutagenesis method can be easily adapted for other genes.

Notably, the MaveDB database aggregates data from published DMS studies, but so far, only about 50 human genes have been recorded^106^. Charting the "Atlas of Variant Effects" across the entire human genome should be a crowdsourced effort, thus requiring the associated technologies and resources used in DMS to be accessible to as many laboratories as possible. We developed the SMuRF workflow to reduce DMS cost and complexity to encourage crowd-sourcing efforts required for coverage across thousands of disease genes.

### Limitations of the study

In this study, we decided to explore all possible SNVs, therefore not all amino acid substitutions were represented. However, population sequencing data has demonstrated that multiple SNVs in the same codon (i.e. multi-nucleotide variants) are extremely rare^7^, and the majority of VUS are SNVs^55^, thus justifying that representing all possible SNVs is more practical and cost-effective when data is to be used to interpret variants observed in sequenced individuals. Additionally, it is worth noting that the SMuRF workflow can be extended to analyze small-sized variants beyond SNVs, including single amino acid substitutions, small insertions, or deletions. Future research covering these variants can provide further functional insights.

It is important to consider certain factors when utilizing scores from SMuRF or other DMS studies. The majority of dystroglycanopathies are recessive and patients may harbor compound heterozygous variants, which raises the need for further investigation on how to apply DMS functional scores in interpreting such cases. In this study, we explored a naive additive model where the biallelic functional scores were calculated by the simple addition of the SMuRF scores of the variants on both alleles. This model demonstrated a promising correlation between SMuRF scores and disease severity (Figures 3C and 3D). The higher-allelic score model was also tested, which agreed with the additive model (Figure S3I). However, future experiments that can better model biallelic effects, such as preinstallation of one allele, are required to improve these analyses. Additionally, our results emphasized the significance of well-curated reports in predicting disease severity (Figure S3J). Our findings revealed a correlation between the disease onset age and SMuRF scores (Figures 3D and S3K). However, creatine kinase (CK) values did not show a significant correlation with SMuRF scores (Figure S3L). This observation is consistent with the knowledge in the field that CK levels can fluctuate with activity and decrease when muscle mass is lost over time^107^. Additionally, it is important to acknowledge the need for a larger patient dataset to enhance the predictive accuracy of SMuRF scores for patient phenotypes, including age of onset and disease severity. In our study, we evaluated 54 cases, with 6 cases classified as "Severe". The limited sample size restricts the robustness and reliability of our conclusions. Future cohort research expanding the dataset, will help improve the accuracy.

The use of SMuRF scores for the interpretation of genetic variants also poses challenges in the following aspects. Firstly, the IIH6C4 assay may only capture one of the multiple functions associated with FKRP and LARGE1. For example, FKRP is also known to participate in the glycosylation of fibronectin^108^. Second, the presence of paralogs may impact the clinical relevance of the SMuRF scores. For instance, *LARGE2*, a paralog of *LARGE1* resulting from a duplication event first observed in Chondrichthyes^62^, may act as an effective modifier in *LARGE1*-related diseases^109–111^. Third, SMuRF analysis for nonsense variants has inherent limitations. In the context of the CDS constructs used in SMuRF, a nonsense mutation does not trigger exon-junction complex (EJC)-enhanced NMD. However, in the correct multi-exon genomic context, when a nonsense mutation is located upstream of the last EJC, it undergoes EJC-dependent NMD, further reducing any residual functions that the truncated proteins may possess^112^.

Despite the limitations of this study, the SMuRF scores hold potential for being considered as functional evidence for the PS3/BS3 criterion following ACMG/AMP guidelines. Lastly, a careful curation of evidence from all available sources, including but not limited to clinical reports, DMS, computational predictors, will aid in the generation of a comprehensive and accurate variant interpretation guidebook.

## Key resources table

| REAGENT or RESOURCE | SOURCE | IDENTIFIER |
| --- | --- | --- |
| Antibodies |  |  |
| IIH6C4 antibody | Sigma-Aldrich<br>Dr. Kevin Campbell | 05-593 (Discontinued) |
| Rabbit anti-Mouse IgM FITC Secondary Antibody | Invitrogen | 31557 |
| Primary Anti-FKRP antibody | Abcam | ab220059 |
| Primary Anti-Vinculin antibody | Sigma-Aldrich | V9131 |
| Anti-laminin antibody | Sigma-Aldrich | L9393 |
| Anti-rabbit HRP conjugated secondary antibody | Cell Signaling Technology | 7074S |
| Anti-mouse HRP conjugated secondary antibody | Cell Signaling Technology | 7076S |
| IRDye 800 CW Goat anti-mouse IgM ($\mu$ chain specific) Secondary Antibody | LI-COR Biosciences | 926-32280 |
| anti-rabbit IgG (H + L); donkey polyclonal | LI-COR Biosciences | 926-32213 |
| Bacterial and virus strains |  |  |
| Endura Electrocompetent Cells | Lucigen | 60242-1 |
| rVSV-LASV-GPC viral particles | Dr. Melinda A. Brindley |  |
| ppVSVΔG-VSV-G viral particles | Dr. Melinda A. Brindley | |
| ppVSV-LASV-GPC-Generation 1 | This paper |  |
| ppVSV-LASV-GPC-Generation 2 | This paper |  |
| ppVSV-LASV-GPC-Generation 3 | This paper |  |
| Chemicals, peptides, and recombinant proteins |  |  |
| SpCas9 2NLS Nuclease | Synthego |  |
| SE Cell Line Nucleofector Solution | Lonza | PBC1- 00675 |
| NEBuilder HiFi DNA Assembly Master Mix | NEB | E2621 |
| T4 DNA Ligase | NEB | M0202 |
| SsoAdvanced Universal SYBR Green Supermix | Bio-Rad | 1725271 |
| TransIT-LT1 Transfection Reagent | Mirus | MIR 2300 |
| Lenti-X Concentrator | Takara | 631232 |
| Lenti-X GoStix Plus | Takara | 631280 |
| Blasticidin S HCl | Gibco | A1113903 |
| Puromycin Dihydrochloride | Gibco | A1113803 |
| Versene | Gibco | 15040066 |
| Viability 405/452 Fixable Dye | Miltenyi Biotec | 130-130-420 |
| MACS BSA Stock Solution | Miltenyi Biotec | 130-091-376 |
| autoMACS Rinsing Solution | Miltenyi Biotec | 130-091-222 |
| Human BD Fc Block | BD Pharmingen | 564220 |
| Gelatin | Sigma-Aldrich | G9391 |
| PFA | Sigma-Aldrich | 158127 |
| Bovine Serum Albumin | Sigma Aldrich | A9647 |
| Antifade Mounting Medium with DAPI | Vector Laboratories | H1500 |
| NH<sub>4</sub>Cl | Sigma-Aldrich | A9434 |
| Q5 High-Fidelity DNA Polymerase | NEB | M0491SVIAL |
| Type2M enzyme DpnI | NEB | R0176S |
| Q5 Reaction Buffer | NEB | B9027SVIAL |
| Q5 High GC Enhancer | NEB | B9028AVIAL |
| dNTPs | NEB | N0447 |
| BsmBI-v2 | NEB | R0739S |
| BsaI-HFv2 | NEB | R3733S |
| MspJI | NEB | R0661S |
| XbaI | NEB | R0145S |
| BamHI-HF | NEB | R3136S |
| Protease Inhibitor Cocktail | Sigma-Aldrich | P8340-1ML |
| RIPA lysis and extraction buffer | Pierce | PI89900 |
| 4x Laemlli | BioRad | 1610747 |
| PageRuler Plus Prestained Protein Ladder | ThermoFisher Scientific | 26619 |
| Clarity Max ECL substrate | BioRad | 1705062 |
| Clarity ECL substrate | BioRad | 1705060 |
| UDP-xylose | Sussex Research Laboratories | Custom order |
| UDP-glucuronate | Sigma-Aldrich | U6751 |
| 4-methylumbelliferone-glucuronate-xylose | Sussex Research Laboratories | Custom order |
| Laminin (Natural, mouse) Lam-111 | Gibco | 23017-015 |
| Critical commercial assays |  |  |
| Synthetic Single Guide RNA Kit | Synthego |  |
| PrimeScript RT Reagent Kit | Takara | RR037 |
| Phusion High-Fidelity DNA Polymerase | NEB | M0530 |
| NEBuilder | NEB | E2621 |
| NucleoSpin Gel and PCR Clean-Up Kit | Takara | 740609 |
| PureLink Midiprep Kit | Invitrogen | K210014 |
| PureLink Genomic DNA Mini Kit | Invitrogen | K182002 |
| SE Cell Line 4D-Nucleofector X Kit S | Lonza | V4XC-1032 |
| BioRad DC Assay II Kit | BioRad | 5000112 |
| Experimental models: Cell lines |  |  |
| HAP1 | Horizon Discovery | C631 |
| DAG1-KO HAP1 | Horizon Discovery | HZGHC000120c016 |
| HEK293T | ATCC | CRL-3216 |
| MB135 | Dr. Stephen J. Tapscott |  |
| FKRP-KO HAP1 (c.181Adup) | This paper |  |
| FKRP-KO MB135 (c.181Adup) | This paper |  |
| LARGE1-KO HAP1 (c.121_214del) | This paper |  |
| LARGE1-KO MB135 (c.121_214del) | This paper |  |
| Oligonucleotides |  |  |
| *FKRP*-KO and *LARGE1*-KO sgRNAs, genotyping primers | This paper (See Method S1) |  |
| *DAG1*, *FKRP*, *LARGE1* CDS sequences, primers for CDS plasmid construction | This paper (See Method S2) |  |
| RT-PCR, RT-qPCR, real-time PCR primers | This paper (See Method S3) |  |
| Primers for PALS-C | This paper (See Method S4) |  |
| R1 oligo lists | This paper | https://github.com/leklab/Balthazar/tree/main/PALS-C_oligos |
| Primers for plasmid pool QC | This paper (See Method S6) |  |
| Primers for lentiviral sequence isolation, *FKRP* Sanger sequencing, *FKRP* Amplicon-EZ sequencing, *LARGE1* Sanger sequencing | This paper (See Method S7) |  |
| Primers for PCR1, PCR2 | This paper (See Method S8) |  |
| Primers for NGS library construction for VSV related experiments | This paper (See Method S10) |  |
| Recombinant DNA |  |  |
| pLV-hTERT-IRES-hygro | Hayer et al.<sup>104</sup> | Addgene Plasmid #85140 |
| lentiCRISPR v2 | Sanjana et al.<sup>113</sup> | Addgene Plasmid #52961 |
| lentiCas9-Blast | Sanjana et al.<sup>113</sup> | Addgene Plasmid #52962 |
| pAAV-UbC-eGFP-F | Dr. Pantelis Tsoulfas | Addgene Plasmid #71545 |
| lentiGuide-Puro | Sanjana et al.<sup>113</sup> | Addgene Plasmid #52963 |
| Lenti-DAG1 | This paper | Addgene Plasmid #205149 |
| Lenti-UbC-*FKRP*-EF1a-BSD | This paper | Addgene Plasmid #205150 |
| Lenti-UbC-*LARGE1*-EF1a-BSD | This paper | Addgene Plasmid #205151 |
| psPAX2 | Dr. Didier Trono | Addgene Plasmid #12260 |
| pMD2.G | Dr. Didier Trono | Addgene Plasmid #12259 |
| LASV-GPC | Dr. Melinda Brindley |  |
| Software and algorithms |  |  |
| R | R Core Team | https://www.r-project.org/contributors.html |
| Scripts for Oligo design, pre-SMuRF experiments | This paper | https://github.com/leklab/Balthazar |
| Raw NGS data analytical pipeline | This paper | https://github.com/leklab/Gargamel |
| Scripts for generating SMuRF scores and downstream analyses | This paper | https://github.com/leklab/Azrael |
| ChatGPT 3.5 | OpenAI | https://chat.openai.com/g/g-F00faAwkE-open-a-i-gpt-3-5 |
| Gemini | Google | https://gemini.google.com/ |
| Other |  |  |
| 0.45 μm PES filter | Thermo Scientific | 165-0045 |
| 40 μm Cell Strainer | Falcon | 352340 |
| Thermanox Coverslips | Thermo Scientific | 174969 |
| SMuRF scores | This paper | https://www.mavedb.org/experiments/urn:mavedb:00001197-a |
| FFC and FACS datasets | This paper | FlowRepository (FR-FCM-Z6LL, FR-FCM-Z6LM, FR-FCM-Z6LN, FR-FCM-Z6LP, FR-FCM-Z6LQ, FR-FCM-Z6LR, FR-FCM-Z6LX, FR-FCM-Z77U, FR-FCM-Z794) |
| NGS raw data | This paper | Sequence Read Archive of the National Center for Biotechnology Information (accession: PRJNA993285). |
| Online protocol | This paper | https://www.protocols.io/view/using-saturation-mutagenesis-reinforced-functional-cwvvxe66/materials |
| 4-15% mini-protean TGX stain free protein gel | BioRad | 4568084 |
| Nitrocellulose Membrane | BioRad | 1704158 |
| WGA resin | Vector Laboratories | AL-1023 |

## RESOURCE AVAILABILITY

### Lead Contact

Further information and requests for resources and reagents should be directed to and will be fulfilled by the lead contact, Monkol Lek.

### Materials Availability

- Plasmids generated in this study have been deposited to Addgene: Lenti-*DAG1* (205149), Lenti-UbC-*FKRP*-EF1α-BSD (205150), and Lenti-UbC-*LARGE1*-EF1α-BSD (205151)

### Data and Code Availability

- NGS raw data have been deposited at the Sequence Read Archive (SRA) of the National Center for Biotechnology Information (NCBI) and are publicly available as of the date of publication. Accession numbers are listed in the Key Resources Table.
- SMuRF scores have been deposited on MaveDB. The link is listed in the Key Resources Table.
- FFC and FACS datasets have been deposited on FlowRepository. IDs are listed in the Key Resources Table.
- All original code has been deposited on GitHub (https://github.com/leklab) and is publicly available as of the date of publication. DOIs are listed in the key resources table.
- Online protocol has been deposited at https://www.protocols.io/view/using-saturation-mutagenesis-reinforced-functional-cwvvxe66/materials.
- Any additional information required to reanalyze the data reported in this paper is available from the lead contact upon request.

## EXPERIMENTAL MODEL AND STUDY PARTICIPANT DETAILS

### Cell Lines

- Wildtype HAP1 (C631) and *DAG1*-KO HAP1 (HZGHC000120c016) cells (male lacking Y chromosome) were ordered from Horizon Discovery. All HAP1 cells were cultured at 37°C in Iscove’s Modified Dulbecco’s Medium (IMDM) (Gibco, 12440053) with 10% Fetal Bovine Serum (FBS, R&D Systems, S11150) and 1x Antibiotic-Antimycotic (Anti-anti, Gibco, 15240062). The medium was replaced every 2 days, unless otherwise stated. HAP1 cells tend to grow into multi-layers; hence, to keep the cells in optimal status, TrypLE Express Enzyme (Gibco, 12605010) was used to passage the cells to maintain the cells in healthy confluency (30-90%). HAP1 cells used in SMuRF were immortalized using lentivirus packaged with pLV-hTERT-IRES-hygro (Addgene, 85140), a gift from Dr. Tobias Meyer.
- HEK293T cells (female) were cultured at 37°C in DMEM (Gibco, 11995065) with 10% FBS and 1x Anti-anti. The medium was replaced every 2 days, unless otherwise stated.
- MB135 cells (female) were cultured at 37°C in Ham’s F-10 Nutrient Mix (Gibco, 11550043) with 20% FBS, 1x Anti-anti, 51 ng/ml dexamethasone (Sigma-Aldrich, D2915) and 10 ng/mL basic fibroblast growth factor (EMD/Millipore, GF003AF-MG). The medium was replaced every 2 days, unless otherwise stated. MB135 cells were differentiated in Skeletal Muscle Differentiation Medium (PromoCell, C-23061) with 1x Anti-anti. The differentiation medium was replaced every 4 days, unless otherwise stated.

## METHOD DETAILS

### CRISPR RNP nucleofection

Synthetic Single Guide RNA (sgRNA) Kits and SpCas9 2NLS Nuclease were ordered from Synthego. RNP complexes were prepared in SE Cell Line Nucleofector Solution (Lonza, PBC1-00675) and delivered into cells with a Lonza 4D-Nucleofector. The program used for HAP1 was EN-138; the program used for MB135 was CA-137. Single clones were isolated from pooled nucleofected cells and genotyped by targeted Sanger sequencing. sgRNA sequences, RNP complex preparation conditions, and genotyping primers are provided in Method S1. *FKRP*-KO HAP1 carries a 1-bp insertion (c.181Adup); *FKRP*-KO MB135 is homozygous for the same mutation. *LARGE1*-KO HAP1 carries a 94-bp deletion (c.121_214del); *LARGE1*-KO MB135 is homozygous for the same mutation.

### Plasmid construction

Lenti-*DAG1* plasmid used the backbone of lentiCRISPR v2, which was a gift from Dr. Feng Zhang (Addgene, 52961). *DAG1* coding exons were cloned from human genome DNA by PCR. Lenti-*FKRP* plasmids and Lenti-*LARGE1* plasmids used the backbone of lentiCas9-Blast, which was a gift from Dr. Feng Zhang (Addgene, 52962). *FKRP* coding exon was cloned from HAP1 genome DNA. *LARGE1* coding sequence was cloned from HEK293T cDNA. HEK293T carries a *LARGE1* mutation (c.1848G>A) on one allele, which was removed from the Lenti-*LARGE1* plasmids to make the pooled variant library. The removal of this mutation used the same strategy as the introduction of individual variants to the lentiviral plasmids for the mini-libraries: briefly, a short localized region was cut with restriction enzymes from the wildtype plasmid and 2 variant-carrying inserts, each covering 1 of 2 sides of this region were inserted. The UbC promoter was cloned from pAAV-UbC-eGFP-F, which was a gift from Dr. Pantelis Tsoulfas (Addgene, 71545). The EF-1α promoter was taken from lentiGuide-Puro, which was a gift from Dr. Feng Zhang (Addgene, 52963). BSD-WPRE was from lentiCas9-Blast. The lentiviral plasmids used for the pooled library contain a UbC-driven gene-of-interest CDS and an EF-1α-driven *BSD*. *BSD* encodes Blasticidin S deaminase (BSD), which confers blasticidin resistance in transduced cells. Plasmid assemblies were achieved either with NEBuilder HiFi DNA Assembly Master Mix (NEB, E2621) or T4 DNA Ligase (M0202). Cloning details of plasmid construction and the list of plasmids deposited to Addgene are provided in Method S2.

### RT-PCR and RT-qPCR

RT-PCR and RT-qPCR were performed following manufacturers’ manuals. PrimeScript RT Reagent Kit (Takara, RR037) was used for cDNA synthesis. Phusion High-Fidelity DNA Polymerase (NEB, M0530) was used for PCR reactions. SsoAdvanced Universal SYBR Green Supermix (Bio-Rad, 1725271), Hard-Shell 96-Well PCR Plates (Bio-Rad, HSP9601), Plate Sealing Film (Bio-Rad, MSB1001) and Bio-Rad C1000 Touch Thermal Cycler were used for qPCR experiments. Primers are provided in Method S3.

### Lentivirus packaging and transduction

Lentivirus was packaged by HEK293T cells. For a 10-cm dish (90% confluency), 1.5 mL Opti-MEM (Gibco, 31985062), 10 µg psPAX2 (Addgene, 12260), 2 µg pMD2.G (Addgene, 12259), 9 µg lentiviral plasmid, and 50 µL TransIT-LT1 Transfection Reagent (Mirus, MIR 2300) were mixed at room temperature for 15 mins and then added to the cells. 3.5 mL DMEM was added to the cells. 72 hrs later, the supernatant in the dish was filtered with 0.45 μm PES filter (Thermo Scientific, 165-0045), mixed with 5 mL Lenti-X Concentrator (Takara, 631232) and rocked at 4 °C overnight. The viral particles were then spun down (1800 ×g, 4 °C, 1hr) and resuspended in 200 µL DMEM. Lentivirus was titrated with Lenti-X GoStix Plus (Takara, 631280). For lentiviral transduction, the cells to be transduced were plated in wells of plates. One day after seeding, the medium was replaced and supplemented with polybrene (final conc. 8 µg/mL). Lentivirus was then added to the wells for a spinfection (800 ×g, 30 °C, 1hr). One day post-transduction, the medium was replaced, and drug selection was started if applicable. For constructs with BSD, Blasticidin S HCl (Gibco, A1113903, final conc. 5 µg/mL) was used for drug selection. For constructs with PuroR, Puromycin Dihydrochloride (Gibco, A1113803, final conc. 1 µg/mL) was used. Drug selection was performed for 10-14 days.

### PALS-C cloning for saturation mutagenesis

Each variant of all possible CDS SNVs (Figure S2A) was included in a 64-bp ssDNA oligo. The oligos were synthesized (one pool per GOI) by Twist Bioscience. PALS-C is an 8-step cloning strategy to clone lentiviral plasmid pools from the oligos. An elaborate protocol can be found in Method S4. Briefly, the oligos were used as PCR reverse primers, which were annealed to the plasmid template and extended towards the 5’ end of the gene of interest. The resulting products of each block were isolated using block-specific primers. Then the variant strands were extended towards the 3’ end to get the full-length sequences, which were subsequently inserted into the plasmid backbone using NEBuilder (NEB, E2621). The purifications for PALS-C steps were done with NucleoSpin Gel and PCR Clean-Up kit (Takara, 740609). Final assembled products were delivered to Endura Electrocompetent Cells (Lucigen, 60242-1) via electrotransformation (Bio-Rad Gene Pulser II). Transformed bacteria were grown overnight and plasmid pools were extracted using the PureLink Midiprep Kit (Invitrogen, K210014). To check library complexity, colony forming units (CFUs) were calculated and a minimum 18 × coverage was achieved for the plasmid pool of each block of *FKRP* and *LARGE1*. Variants that created new type2S enzyme recognition sites tended to be underrepresented in the pool. These variants are reported in Method S5.

### Plasmid pools QC and saturation mutagenesis

Quality control (QC) was performed for the plasmid pools using the Amplicon-EZ service provided by GENEWIZ (Figure S2B and Method S6). 99.6% of the SNVs of both genes were represented in the plasmid pools (Figure S2C). Lentivirus of each block was packaged by HEK293T cells in one 10-cm dish. Small-scale pre-experiments were performed to determine the viral dosage for optimal separation. GoStix Value (GV) quantified by the Lenti-X GoStix App (Takara) was used to scale the titer of each block to be the same. GV is subject to viral-packaging batch effects; hence, lentiviral pools of all blocks were packaged at the same time using the reagents and helper plasmids of the same batch. Depending on the specific batch and packaging system, 1e3-1e5 GV×µL of lentivirus was used for each block. Considerations for controlling the Multiplicity of Infection (MOI) of the lentiviral pool were discussed in Note S1. For each block, 600k HAP1 cells or 200k MB135 cells were plated in a well of a 6-well plate for transduction. The cell number was counted with an Automated Cell Counter (Bio-Rad, TC20). The cell number for each block was expanded to more than 30M for FACS.

### Mini-libraries for the FACS assay

Mini-libraries of variants were employed to examine and optimize the separation of the FACS assay. Lentiviral constructs were cloned and packaged individually for 8 *FKRP* variants and 3 *LARGE1* variants in addition to the wild-type constructs. These lentiviral particles were mixed to make a mix-9 *FKRP* mini-library and a mix-4 *LARGE1* mini-library. Conditions of transduction, staining, sorting and gDNA extraction were optimized using the mini-libraries. Relative enrichment of variants was defined as the ratio of the variants’ representation in the high-glycosylation sample to their representation in the low-glycosylation group, which was quantified with either Sanger sequencing or Amplicon-EZ NGS (Method S7).

### Fluorescence flow cytometry and FACS

Reagent volumes were determined based on sample size. Below, the staining for samples of one gene block is described as an example. The cells were washed twice with DPBS (Gibco, 14190144), digested with Versene (Gibco, 15040066), and counted. 30M cells were used for staining, which was performed in a 15 mL tube. The cells were spun down (700 ×g, 4 °C, 15 mins) and resuspended in 3mL DPBS supplemented with 30 µL Viobility 405/452 Fixable Dye (Miltenyi Biotec, 130-130-420). All the following steps were done in the dark. The sample was gently rocked at room temperature for 30 mins, and then 7 mL PEB buffer (1 volume of MACS BSA Stock Solution, Miltenyi Biotec, 130-091-376 ;19 volumes of autoMACS Rinsing Solution, Miltenyi Biotec, 130-091-222) was added to the tube. The cells were spun down (700 ×g, 4 °C, 15 mins) and resuspended in 3mL DPBS supplemented with 30 µL Human BD Fc Block (BD Pharmingen, 564220). The sample was gently rocked at room temperature for 30 mins, and then 7 mL DPBS was added. The cells were spun down (700 ×g, 4 °C, 10 mins) and resuspended in 3mL MAGIC buffer (5% FBS; 0.1% NaAz w/v; 10% 10× DPBS, Gibco, 14200166; water, Invitrogen, 10977015) supplemented with 15 µL IIH6C4 antibody (Sigma-Aldrich, 05-593, discontinued; or antibody made in Dr. Kevin Campbell’s lab). The sample was gently rocked at 4 °C for 20 hrs. 7 mL MAGIC buffer was added before the cells were spun down (700 ×g, 4 °C, 10 mins) and resuspended in 3 mL MAGIC buffer supplemented with 60 µL Rabbit anti-Mouse IgM FITC Secondary Antibody (Invitrogen, 31557). The sample was gently rocked at 4 °C for 20 hrs. 7 mL DPBS was added to the sample before the cells were spun down (700 ×g, 4 °C, 10 mins), resuspended with 4 mL DPBS and filtered with a 40 µm Cell Strainer (Falcon, 352340). Note: IIH6C4 antibody from Santa Cruz (sc-73586) is not effective for the experiments listed here.

### Gating parameters

Fluorescence flow cytometry (FFC) experiments were performed with a BD LSR II Flow Cytometer; FACS experiments were performed with a BD FACSAria Flow Cytometer. Forward scatter (FSC) and side scatter (SSC) were used to exclude cell debris and multiplets. Singlets were isolated for downstream analysis. Pacific Blue (450/50 BP) or an equivalent channel was used to detect the Viobility 405/452 Fixable Dye and isolate the live cells for analysis. FITC (530/30 BP), GFP (510/20 BP) or an equivalent channel was used to detect the FITC secondary antibody signal. 20k events were recorded for each block to decide the gating parameters. For FACS, the top ∼20% of the cells were isolated as the high-glycosylation group and the bottom ∼40% of the cells were isolated as the low-glycosylation group. The .fcs files, the FlowJo .wsp files and the software interface reports of the sorter were made available on FlowRepository. A minimum ∼1000 × coverage (*e.g.*, 750k cells harvested for a block with 750 variants) was achieved for both groups of each block.

### NGS library construction

The cells were spun down (800 ×g, 4 °C, 10 mins), and gDNA was harvested from each sample with PureLink Genomic DNA Mini Kit (Invitrogen, K182002). A 3-step PCR library construction was performed to build the sequencing library. Step 1: lentiviral sequence isolation. A pair of primers specific to the lentiviral backbone was used to amplify the lentiviral CDS sequences of each sample. Step 2: block isolation. Each primer contained a 20-bp flanking sequence of the specific block and a partial Illumina adaptor sequence. Forward primers contained the barcodes to distinguish the high-glycosylation group and the low-glycosylation group. Step 3: adaptor addition. Step 2 products were multiplexed and the rest of the Illumina adaptor was added to the amplicons. An elaborate protocol is provided in Method S8. The NGS libraries were sequenced using Psomagen’s HiSeq X service. ∼400M reads were acquired per library.

### Variant counting and scoring

Raw sequencing data was processed with the “Gargamel” pipeline. Variant counting was performed after the data cleaning process that essentially removed the reads with more than one variant. Enrichment of a variant (E_var) in a FACS group is calculated as a ratio of the count of the variant (c_var) to the total count (c_total) at the variant site:

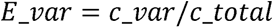

Enrichment of the WT (E_WT) is calculated separately for each block. E_WT is calculated as a ratio of the number of the reads without variant (r_WT) to the number of the reads with one or no variant (r_clean).

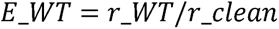

Relative enrichment (rE) is a ratio of the enrichment in the high-glycosylation group to the enrichment in the low-glycosylation group:

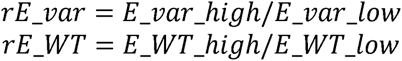

The functional score of a variant (in one biological replicate) is calculated as the ratio of its relative enrichment to that of the WT sequence in the corresponding block:

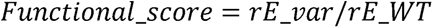

Count of variants and reads were generated from raw sequencing data using the analytical pipeline deposited in the GitHub repository Gargamel. SMuRF score is generated by combing 3 biological replicates (see below).

### Fitness score combination using DiMSum

The DiMSum pipeline was employed to process the variant counts to generate a fitness score across all biological replicates for *FKRP* and *LARGE1*. For data preprocessing, the input files were prepared by block to account for block-specific WT variant counts. Variants with a count of 0 were replaced by the WT value. The variant counts for each replicate within each gene block were processed through DiMSum (https://github.com/lehner-lab/DiMSum/) from the STEAM stage. Default settings were used for the pipeline, aside from *--sequenceType noncoding* to calculate the scores on a variant-level instead of AA-level. The pipeline aggregated these counts to generate a fitness score for each variant. DiMSum incorporates both the frequency of each variant and the observed effects on fitness across all replicates and blocks, producing a merged fitness score for *FKRP* and *LARGE1*. To ensure accurate and reliable fitness estimates, DiMSum employs an advanced error modeling approach. The model addresses multiple sources of error to produce robust fitness scores. Firstly, Poisson-based errors are used to account for the high variance in fitness estimates that arise from low sequencing depth. Since count data often exhibit over-dispersion relative to a Poisson distribution, DiMSum integrates both multiplicative and additive error terms to model this variance effectively. Multiplicative errors, which scale with the variant sequencing counts, address variability arising from workflow-related inconsistencies. In contrast, additive errors are independent of sequencing counts and affect all variants uniformly, accounting for differential handling across replicates. These error components are summarized into a variable, sigma (σ), which is used to adjust fitness estimates on a variant-level. The primary output from the DiMSum pipeline is the 1) “DiMSum score” – fitness score from the direct output of the DiMSum pipeline, converted to the log2 scale, and 2) sigma – a numerical value representing the reliability of the variant count based on error modeling.

### Fitness score normalization

To normalize the fitness scores, we used all synonymous variants as the reference, assuming these variants should have no impact and thus a fitness score of 1. We first transformed the fitness scores of the synonymous variants from their natural logarithm form to a linear scale using the exponential function. For each block, we calculated the mean fitness score of the synonymous variants, treating this mean as the scaling factor. Next, we normalized the fitness scores of all variants by transforming their scores back to the linear scale, dividing by the block’s scaling factor, and then applying a log2 transformation to the normalized values to get the “SMuRF score”. This approach ensures that the scores are adjusted relative to the neutral impact of synonymous variants, allowing for consistent comparisons across different blocks.

### Confidence score generation and classification

Following the DiMSum pipeline for variant score combination and error modeling, each identified variant was assigned a sigma (σ) value representing the estimated error. A higher sigma indicates a higher degree of error in the variant assessment. To classify variants into high and low confidence groups, we applied a threshold based on the 1.5 * interquartile range (IQR) rule, a commonly used outlier detection method. First, we calculated the first quartile (Q1) and third quartile (Q3) of the sigma distribution across all variants. The interquartile range (IQR) was then determined as IQR = Q3 - Q1. The upper threshold for high confidence variants was set at Q3 + 1.5*IQR. Any variant with a sigma value exceeding this threshold was classified as low confidence, while variants below the threshold were designated as high confidence. For all downstream analyses, low confidence variants were excluded to ensure the robustness of results. This approach provided an objective method to filter out potentially unreliable sites prior to further examination and interpretation of the data.

### ClinVar classification

ClinVar classification of the variants was downloaded from the ClinVar webpage through its “Download” option as tabular files. The data were restructured using customized scripts to facilitate the relevant analyses. The version we downloaded was from April 20^th^, 2023. The ClinVar classification of the variants was included in Tables S2 and S3.

### Comparison of SMuRF score to *in silico* predictors

To compare the performance of SMuRF with various computational predictors, we conducted a receiver operating characteristic (ROC) curve analysis. This analysis assesses the ability of each method to identify pathogenic variants, using the ClinVar annotations as the reference standard. For *FKRP* and *LARGE1*, we used a dataset containing annotated missense variants with functional scores from SMuRF and computational predictors including CADD, metaSVM, REVEL, MVP, MutScore, PrimateAI-3D, EVE, ESM1b, and MAVERICK. First, we subsetted the data to include only missense variants. We then further restricted the dataset to include only those variants scored by all computational predictors to ensure a fair comparison. The positive reference set comprised missense variants classified as pathogenic (P), likely pathogenic (LP), or pathogenic/likely pathogenic (P/LP) in ClinVar. The negative reference set consisted of missense variants that are observed in gnomAD v4 genomes and not classified as P, LP, or P/LP in ClinVar. To perform the ROC curve analysis, we employed the pROC package in R. First, we created a binary response variable indicating whether each variant is part of the positive reference set or the negative reference set. Using the roc function from the pROC package, we computed ROC curves for each tool. The ROC curve plots the true positive rate (sensitivity) against the false positive rate (1-specificity) at various threshold settings. The area under the curve (AUC) value was computed for each tool, providing a quantitative measure of their discriminatory ability.

### Correlation of SMuRF scores with disease onset

To investigate the correlation between SMuRF scores and disease onset, a Cox proportional hazards model was employed. The dataset included SMuRF scores for biallelic variants and the corresponding ages of disease onset. Entries with missing values for these variables were excluded. The combined functional score for each patient was calculated as logarithm base 2 of the sum of the exponentiated individual SMuRF scores for each of the biallelic variants.

The age of onset was used as a quantitative variable in months. An event indicator variable was defined to distinguish between disease onset at birth (0 months) and later onset (>0 months). This binary event indicator was coded as 1 for non-zero onset months and 0 for onset at birth.

The Cox proportional hazards model was applied using the *coxph* function from the *survival* package in R. The model evaluated the association between the combined functional score and the age of disease onset. In this analysis, the age of onset was treated as the time variable, the occurrence of the event (disease onset) was treated as a binary outcome variable, and the combined functional score (SMuRF score) was the predictor variable.

After fitting the model, the summary statistics were extracted and interpreted to assess the significance and strength of the association between the functional scores and disease onset. This included evaluating the hazard ratio and its confidence intervals, as well as the p-value for the predictor variable. These results provided insight into how the combined functional score influenced the likelihood of earlier disease onset, offering a quantitative measure of its predictive capacity.

### Immunoblotting for LARGE1 enzymatic activity

Cell pellets expressing LARGE1 variants were resuspended in 50 mM tris pH 7.6, 150 mM NaCl, 1% Triton X-100 containing protease inhibitors (PMSF, benzamide, aprotinin, leupeptin and pepstatin A) and rotated at 4°C for 1 hr. The cleared lysate was incubated with WGA resin (Vector laboratories) for 1 hr at 4°C with rotation. The resin was washed two times with the same buffer with reduced triton X-100 (0.1%). The third wash was completed in matriglycan reaction buffer (20 mM PIPES pH 6.6, 150 mM NaCl, 2 mM CaCl_2_, 2 mM MgCl_2_ and 2 mM MnCl_2_). An equivalent volume as resin of a reaction master mix to a final concentration of 1 mM UDP-xylose (Sussex Research Laboratories), 1 mM UDP-glucuronate (Sigma-Aldrich), and 0.4 mM 4-methylumbelliferone-glucuronate-xylose (Sussex Research Laboratories) in reaction buffer containing protease inhibitors was added. The mixture was incubated at 37°C for 72 hrs. The reaction was terminated with 50 mM EDTA. The cleared supernatant was resolved on a strong anion exchange column using a gradient of increasing anionic strength up to 500 mM NaCl over a background of 50 mM ammonium phosphate pH 6.0. The resin was combined with SDS loading buffer and resolved on a homemade 3-15% gradient SDS-PAGE at 60 V for 16 hrs at room temperature. Proteins were stained with Bio-Safe Coomassie G-250 Stain according to the manufacturer’s instructions (Bio-Rad) and imaged on the Li-COR. The proteins were transferred to PVDF membrane (800 mA for 5 hrs at 4°C), blocked with 2% milk in 50 mM tris pH 7.6 and 75 mM NaCl (low salt-TBS) and incubated with anti-matriglycan primary antibody (IIH6C4) overnight at 4°C with rocking. The membrane was washed three times with low salt-TBS prior to incubation with anti-mouse IgM secondary antibody for 30 minutes at room temperature with rocking. The membrane was washed three times and imaged on a Li-COR. Another PVDF membrane was incubated with 5% milk dissolved in LBB buffer (10 mM triethanolamine, 140 mM NaCl, 1 mM CaCl_2_ and 1 mM MgCl_2_). The membrane was washed three times with 3% BSA in LBB buffer and then incubated with 7.5 nM mouse laminin (Lam-111) dissolved in LBB overnight at 4°C. The membrane was washed three times with LBB and then incubated with anti-laminin antibody dissolved in 5% milk-LBB overnight at 4°C. The membrane was washed three times with LBB and incubated with anti-rabbit IgG for 30 minutes at room temperature. The membrane was washed three times and imaged on a Li-COR.

### Visualization of data on protein 3D structures

Protein 3D structures from Protein Data Bank (PDB) were visualized using UCSF ChimeraX v1.3^114^. The crystal structure of human FKRP (PDB:6KAM)^85^ and the electron microscopy structure of LARGE1 (PDB:7UI7)^86^ were used. Figures displaying domain locations or the log2 mean missense score per residue on 3D structures were generated using custom ChimeraX command files. Domain coordinates displayed are per Ortiz-Cordero *et al.* for FKRP^87^, and Joseph *et al.* for LARGE1^86^. The log2 of the mean missense score per residue was calculated using a custom Python script.

### Immunofluorescence

15 mm round Thermanox Coverslips (Thermo Scientific, 174969) were placed in the wells of 24-well plates. To coat the coverslips, 0.1% gelatin (Sigma-Aldrich, G9391) was added to the wells and immediately removed. After the coverslips were air-dried, 250k MB135 cells were resuspended in 0.5 mL growth medium and seeded into each well. One day after plating the cells, the medium was changed to the differentiation medium, and cells were differentiated for 3-7 days until mature myotubes were formed. The cells were washed with DPBS and fixed with 4% PFA (Sigma-Aldrich, 158127) for 10 mins at room temperature. The cells were blocked with 2% Bovine Serum Albumin (BSA, Sigma-Aldrich, A9647) at room temperature for 1 hr before undergoing incubation with the IIH6C4 antibody (1:200 in 2% BSA, Sigma-Aldrich, 05-593, discontinued) at 4 °C for 20 hrs. The cells were then washed with DPBS before undergoing incubation with the secondary antibody (1:100 in 2% BSA, Invitrogen, 31557) at room temperature for 2 hrs in the dark. Antifade Mounting Medium with DAPI (Vector Laboratories, H1500) was dropped onto Microscope Slides (Fisher Scientific, 22-037-246). The coverslips with cells were washed again with DPBS, put facedown onto the drops on the slides, and kept at room temperature for 30 mins in the dark. Pictures were taken with a Revolve ECHO microscope. Note: IIH6C4 antibody from Santa Cruz (sc-73586) is not effective for the experiments listed here.

### Packaging and infection of rVSV / ppVSV

rVSV-LASV-GPC viral particles, ppVSVΔG-VSV-G viral particles, and the LASV-GPC plasmid were obtained from Dr. Melinda Brindley. To package ppVSV-LASV-GPC viral particles, HEK293T cells were transfected with the LASV-GPC plasmid and then transduced with ppVSVΔG-VSV-G viral particles. The resulting particles were referred to as ppVSV-LASV-GPC-Generation1. A new batch of LASV-GPC transfected HEK293T cells were subsequently transduced with ppVSV-LASV-GPC-Generation1 to produce ppVSV-LASV-GPC-Generation2, reducing residual VSV-G in the pseudotyped particles. Similarly, later generations can be packaged. The experiments in this study utilized ppVSV-LASV-GPC-Generation2 and 3. The 50% tissue culture infectious dose (TCID50) of the VSV was determined using the Spearman-Karber method^115,116^. Lentiviral transduction and blasticidin drug selection were performed in the same manner as those in the FACS assay. Afterward, cells were divided into two groups (∼1M cells each): a no-infection group and an infection group. rVSV infection was conducted at an approximate MOI of 0.5. NH4Cl (Sigma-Aldrich, A9434, final conc. 5mM) was added during the infection and subsequent recovery. After 60 hours of infection, the medium was replaced, and the cells were allowed to recover for 12 hours before harvesting. ppVSV infection was performed at an approximate MOI of 1∼3, and the infected cells were recovered to ∼1M prior to harvesting. Detailed ppVSV packaging protocol and considerations for application are provided in Method S10.

### Validation using the ppVSV assay

10 Lenti-*FKRP* variants were mixed with Lenti-WT-*FKRP* to rescue *FKRP*-KO MB135. 11 Lenti-*LARGE1* variants were mixed with Lenti-WT-*LARGE1* to rescue *LARGE1*-KO MB135. The lentiviral plasmids were individually constructed and validated using the whole-plasmid sequencing service provided by Plasmidsaurus. The individual plasmids were mixed to package the *FKRP* mix11 and *LARGE* mix12 lentiviral pools. Lentiviral transduction, drug selection, and ppVSV infection were performed similarly as stated above. After harvesting the cells, the lentiviral GOI sequence was amplified with the PCR1 protocol in Method S8 and sequenced with nanopore sequencing provided by Plasmidsaurus. The functional score was quantified by the ratio of a variant’s enrichment in the non-infected group to its enrichment in the ppVSV-infected group. The list of the variants included in the mini-libraries and other details are provided in Method S11.

## QUANTIFICATION AND STATISTICAL ANALYSIS

### Statistical Analysis

Two-sided Wilcoxon tests were performed with the “ggsignif” R package. Spearman’s rank correlation coefficients were calculated with the “cor.test” function in R. Multiple comparisons between groups were performed using analysis of variance (ANOVA) followed by Bonferroni post hoc test through GraphPad Prism 10.2.2.

## ADDITIONAL RESOURCES

Detailed experimental protocols: https://www.protocols.io/view/using-saturation-mutagenesis-reinforced-functional-cwvvxe66/materials.

## Acknowledgments

We thank Sander Pajusalu and Heather Best for their efforts which helped lead to this project. We thank Nicholas Johnson and the GRASP LGMD consortium for granting us access to their deidentified patient data, which was funded by ML Bio Solutions. We thank Stephen Tapscott for the gift of MB135 cells. This work is supported by a grant to M.L. from the Muscular Dystrophy Association (629095) for “improved clinical interpretation of rare variants in muscle diseases”. K.P.C. is an investigator of the Howard Hughes Medical Institute. This work was supported in part by a Paul D. Wellstone Muscular Dystrophy Specialized Research Center grant (1U54NS053672 to K.P.C.). M.A.B. was supported by the National Institute of Allergy And Infectious Diseases of the National Institutes of Health under Award Number R01AI139238 (MAB). S.H. is supported by a Muscular Dystrophy Association Development Grant (MDA 963708). J.C. is supported by The Chris Carrino Foundation for FSHD and NIH/5F32AR079892-02. We thank Yale Flow Cytometry for their assistance with FFC and FACS services. The Core is supported in part by an NCI Cancer Center Support Grant # NIH P30 CA016359.

## Author contributions

K.M. and M.L. conceived and designed this study. M.L. supervised the experiments and analyses of this study. K.M. designed and performed the experiments for the establishment, application, and validation of SMuRF, with the help of S.H. and other authors. K.K.N., K.M., and M.L. developed the analytical pipeline. S.H. and K.M. performed the IF experiments. S.J. conducted the matriglycan-related validation experiments. K.M. and A.L. created the lentiviral plasmid pools. K.G.W. and K.E.K. participated in the early development of this study. V.H. and C.L.O’C. assisted in performing the experiments. S.J. and K.P.C. participated in the analysis of IIH6C4-related experiments. M.A.B. participated in designing and interpreting the VSV experiments. K.M., K.K.N., N.J.L. and M.L. performed computational analyses, with the help of J.X. and L.G.. K.M., K.K.N., N.J.L., J.X., J.C., and M.L. wrote this manuscript with the help of other authors.

## Declaration of generative AI and AI-assisted technologies

We used ChatGPT 3.5 and Gemini to improve the readability and language in this manuscript. The manuscript was first drafted by us and polished with the tools sentence-by-sentence where we deemed necessary. We then reviewed and finalized the text. We take full responsibility for the contents of this manuscript.

## Supplemental information

Document S1. Figures S1–S7, Table S6 and Note S1

Table S1. Datasets collected for the generation of SMuRF scores.

Table S2. SMuRF scores and other scores used in this manuscript for *FKRP*.

Table S3. SMuRF scores and other scores used in this manuscript for *LARGE1*.

Table S4. Restructured data from 8 well-curated FKRP cohorts, related to Figure 3C and 3D.

Table S5. Variant inclusion details for the generation of the ROC curves, related to Figure 4A and 4B.

## Note S1: Considerations for implementation and future development of SMuRF

The utilization of lentiviral delivery primarily stems from technical and economic considerations. Delivery of variants through this exogenous method is artificial, as it removes the genes’ cis-regulatory elements and intron/exon structures. On-site saturation genome editing is preferred for maintaining genomic context. However, lentiviral systems offer benefits by increasing the useful event for flow cytometric assays. Currently, saturation genome editing is limited by its efficiency, which is heavily influenced by nature of specific genomic regions. Achieving comprehensive unbiased high coverage for saturation mutagenesis in certain genes is challenging. Particularly in flow cytometric assays, low efficiency results in longer sorting times and poses significant economic challenges. Lentiviral delivery presents a trade-off where genomic context is sacrificed for more comprehensive coverage.

The impact of the Multiplicity of Infection (MOI) of the lentiviral pool on the outcome of the functional characterization is profound (Figure S7A). Several factors might need to be considered here: (1) Higher MOI may render higher expression of the gene of interest (GOI), thus compensating for the pathogenic effects of some pathogenic variants. (2) Different levels of MOI result in different co-existing status of different variants in the same single cell. Whether co-existing variants can generate modifying effects through protein dimerization, and whether different co-existing combinations generate different modifying effects (positive or negative), are unclear and require further research. In the SMuRF workflow, for the purpose of better separating variants with different levels of function, the MOI was controlled to ensure that the lentiviral GOI RNA level is comparable to the endogenous level in WT cells. The titer was determined in pre-experiments. Future improvement to the workflow entails identifying and employing the endogenous promoter core element as a replacement for the UbC promoter^1^.

We utilized rVSV-LASV-GPC or ppVSV-LASV-GPC to infect *GOI*-KO MB135 myoblasts that had been rescued by all-possible Lenti-*GOI* variant pools. The rVSV genome incorporates the LASV-GPC coding sequence, allowing it to re-enter cells. In contrast, the ppVSV lacks this capability as it is pseudotyped using a LASV-GPC plasmid (STARl7JMethods). Likely due to the re-entering feature of rVSV, the rVSV screen did not yield meaningful results (Figure S7B). The ppVSV screen for both FKRP and LARGE1 showed a tendency where start-loss variants were the most enriched in the infected group, suggesting a higher disruptive effect on α-DG glycosylation. Nonsense variants were the next most enriched, followed by missense variants, while synonymous variants were the least enriched (Figures S7C and S7D). This tendency aligns with what we observed in the SMuRF FACS assay. However, the ppVSV assay, in general, lacked the sensitivity to distinguish differences among the variants in the whole-CDS scale. Therefore, we employed the ppVSV assay for the mini-library rescued myoblasts (Figures 6D and 6E). In the *FKRP* mini-library, we also included *FKRP* c.1174T>G (p.Phe392Val; SMuRF=-1.13; SMuRF_mild, Confidence=HIGH), which is a previously unknown variant and has been observed in a patient with compound mutations (Leu276Ile on the other allele). ppVSV assay supports the notion that this variant is mildly pathogenic.

Synonymous variants are typically considered to have limited effects on gene function^2^. In our study, we adopted the assumption that synonymous variants have no effects when normalizing our scores. However, it is possible that some synonymous variants can affect RNA motifs, structure, or splicing, leading to potential changes in gene function^3–5^. While the effects of synonymous variants on splicing cannot be evaluated within the SMuRF framework, their impact on certain RNA motifs can be captured by SMuRF. We selected two synonymous variants in *LARGE1*, c.639T>C (SMuRF=0.45) and c.642A>G (SMuRF score=0.64), for evaluation, which might have gain-of-function (GOF) effects by removing a poly(A) motif from the coding sequence (Figure S7E). The poly(A) signal is associated with transcription termination, and its removal may increase the level of functional transcripts^6^. However, validation using the orthogonal ppVSV assay did not support the hypothesis of these 2 synonymous variants having GOF effects (Figure 6E). Nevertheless, it’s essential to note that even if this GOF effect is genuine, it may not fully apply to the endogenous *LARGE1* in its correct genomic context, despite the fact that this poly(A) motif is not separated by exon junctions *in vivo*.

synVep (synonymous Variant effect predictor) is a machine learning-based predictor for evaluating the effects of synonymous SNVs^5^. Higher synVep scores (range: 0-1) indicate stronger potential effects. We compared the synVep scores with the SMuRF scores for *FKRP* and *LARGE1* (Figure S7F). For *FKRP*, c.783A>T has an ultra-low SMuRF score and is also predicted to have strong effects by synVep (SMuRF=-3.42, synVep=0.99). For *LARGE1*, c.477C>A has an ultra-low SMuRF score and is also predicted to have strong effects by synVep (SMuRF=-2.47, synVep=0.99). *FKRP* c.783A>T, together with c.783A>C (SMuRF=-2.62, confidence=LOW, synVep=0.92) and c.783A>G (SMuRF=0.02, synVep=0.83), was included in our ppVSV test, as well as *LARGE1* c.690T>A (SMuRF=-1.28, synVep=0.95), c.693T>G (SMuRF=-1.33, synVep=0.98) and c.699C>G (SMuRF=-1.67, synVep=0.99). However, the results did not support any of them having LOF effects (Figures 6D and 6E). Further research, such as endogenous gene editing, is required to fully evaluate the effects of these synonymous variants.

LoGoFunc is an ensemble machine-learning predictor for pathogenic GOF, pathogenic LOF, and neutral genetic variants^7^. While it did not predict the existence of any GOF variants in *FKRP*, it did predict the existence of several GOF variants in *LARGE1*, including c.26G>C (p.Arg9Pro) (Figure S7G). We examined c.26G>C (SMuRF=0.18, LoGoFunc_GOF=0.36) with the ppVSV assay, but the result did not support the notion that this variant possesses GOF effects (Figure 6E).

Consistent with what was shown in Figure 5C, nonsense variants also generally exhibit higher SMuRF scores in the GlcAT region of *LARGE1* than those in the XylT region (p-value=0.00034, Figure S7H). Since the SMuRF system employs the CDS of the gene, the nonsense mutations are expected to produce truncated protein without significantly affected by EJC-enhanced NMD. It may be assumed that the SMuRF score difference between the GlcAT region and the XylT region is partly technical. That is, in the absence of a functional GlcAT domain, a functional XylT domain can still add one single xylose unit to α-DG, which may be sufficient to be detected by IIH6C4, albeit generating a very weak signal. In this scenario, the IIH6C4 signal may not truthfully represent the physiological function, as it is uncertain how much function a single xylose possesses in comparison to the complete matriglycan chain. Nevertheless, AlphaMissense also presents a similar result, where missense variants generally exhibit lower AlphaMissense scores (higher functionality) in the GlcAT region of *LARGE1* than those in the XylT region (p-value<2.22e-16, Figure S4C). This suggests that the difference in these two domains is to some extent biologically relevant, that is the XylT domain may possess a more important functional role than the GlcAT domain.

It has been previously reported that FKTN, FKRP, and RXYLT1 (TMEM5) may form a complex^8^. The SMuRF scores of SNV-generated single amino acid substitutions were projected to the FKTN-FKRP-RXYLT1 complex predicted with Alphafold2^9,10^ (Figure S5D). One interesting residue identified by SMuRF is p.Trp255 (mean SMuRF=-0.79), which is predicted to interact with FKTN p.Leu263. *FKTN* c.787C>G (p.Leu263Val) and *FKTN* c.788T>G (p.Leu263Arg) are variants of uncertain significance in ClinVar (Oct. 11^th^, 2023).

## Method S1

*FKRP*-KO sgRNA:

GTTCGAGGCATTTGACAACG

*LARGE1*-KO sgRNAs (used as a mixture): GATGGGATGGGGCTCGGCCC GGTGCGCATGCGCGAGGTGG TCCAGCGGTGACAGAGACAC

RNP preparation:

18 µL supplemented SE Solution + 6 µL 30 µM sgRNA + 1 µL 20 µM Cas9 Room temperature 10 mins

Add to 150 k cells (spun down 100 ×g, 10 mins; resuspended in 5 µL supplemented SE Solution) SE Cell Line 4D-Nucleofector X Kit S was used (Lonza, V4XC-1032)

Pick single clones:

Allow the nucleofected cells to recover in the growth medium; plate the nucleofected cells sparsely and allow them to form monoclonal clusters; pick monoclonal cells under microscope.

Genotyping primers:

FKRP-GT-F: CATCACCCTCAACCTTCTGGTC FKRP-GT-R: CATCAGGTACTAGGGCCACAAACTC LARGE1-GT-F: GGCAATCGGGACTTTGGACA LARGE1-GT-R: GCCTCGCCATGTAGTAAGGG

## Method S2

Q5 High-Fidelity DNA Polymerase (NEB, M0491SVIAL) was used for the PCR reactions according to the manufacturers’ manuals.

The *DAG1* CDS sequence in this project is:

ENST00000308775.7 but carrying a common missense variant c.41C>G (p.Ser14Trp)

The Allele Frequency of this variant is 0.9762 in gnomAD v3.1.2 and 0.9955 in gnomAD v4.0.0

The *FKRP* CDS sequence in this project is as in: ENST00000318584.10

The *LARGE1* CDS sequence in this project is as in: ENST00000354992.7

Primers for CDS plasmid construction:

DAG1-Exon2-F: GGTTCTAGAGCGCTGCCACCATGAGGATGTCTGTGGGCCTC DAG1-Exon2-R: CCGCTGATACCTTGATGATATCTCCACTGGAGGCAAT

DAG1-Exon3-F: TATCATCAAGGTATCAGCGGCAGGGAAGGAG

DAG1-Exon3-R: AAGTTTGTTGCGCCGGATCCAGGTGGGACATAGGGAGGAGG

FKRP-clone-F (matching part): ATGCGGCTCACCCGCT FKRP-clone-R (matching part): GCCGCTTCCCGTCAGA

LARGE1-clone-F (matching part): ATGCTGGGAATCTGCAGGG LARGE1-clone-R (matching part): CTAGCTGTTGTTCTCGGCTGTGAG UbC-clone-F (matching part): GGCCTCCGCGCCGG

UbC-clone-R (matching part): ACCAAGTGACGATCACAGCGATCC EF1α-clone-F (matching part): GGCTCCGGTGCCCGTCA

EF1α-clone-R (matching part): GGTGGCCGTACGTCACGACA

BSD-WPRE-clone-F (matching part): ATGGCCAAGCCTTTGTCTCAAGAAG BSD-WPRE-clone-R (matching part): GCGGGGAGGCGGCC

*The overhangs were designed according to different assembly schemes.

Two cloning strategies to introduce individual variant to the WT plasmid:

1. Cut and Swap: A short, localized region was cut with restriction enzymes from the wild-type plasmid to serve as the backbone. Two variant-carrying inserts were amplified using PCR. The variant of interest was introduced in the R primer for the 5’ insert and the F primer for the 3’ insert. The backbone and those two inserts were assembled with NEBuilder HiFi DNA Assembly Master Mix (NEB, E2621)
2. Site-directed mutagenesis: The whole plasmid was amplified using a pair of primers carrying the variant of interest. The WT plasmid was removed from the reaction using Type2M enzyme DpnI (NEB, R0176S). F primers were designed as: (5’->3’) 7-bp flanking sequence A + 1-bp variant + 7-bp flanking sequence B + 15-bp matching sequence; R primers were designed as (5’->3’) 7-bp flanking sequence B + 1-bp variant + 7-bp flanking sequence A + 15-bp matching sequence.

Oxford Nanopore long-read sequencing services were provided by Plasmidsaurus to validate the plasmid sequences.

Plasmids deposited to Addgene:

Lenti-*DAG1*: 205149

Lenti-UbC-*FKRP*-EF1α-*BSD*: 205150

Lenti-UbC-*LARGE1*-EF1α-*BSD*: 205151

## Method S3

RT-PCR, RT-qPCR and real-time PCR primers:

DAG1-cDNA-F: GATCTGCTACCGCAAGAAGC

DAG1-cDNA-R: ATGGTGTCCTGGTTCAGAGG

FKRP-rtPCR-F: TGGGCATCTACTTGGAGGAC

FKRP-rtPCR-R: TTGCTTTCGCTGTACTGCAC

LARGE1-cDNA-F: GAGCAGTGCTACAGAGACGTGT

LARGE1-cDNA-R: TGCCGTCATACTCCAGGAAGGT

LARGE1-gDNA-F: ATCAGCCTGGCCCTCTACC

LARGE1-gDNA-R: TGTAGGGAGTGCTGATGTGC

## Method S4

Primers used for PALS-C:

PALS-C-universal-F1: GAACAGGCGAGGAAAAGTAG

PALS-C-universal-F2: GATCGTCACTTGGTACCGGTTCTAGA

PALS-C-universal-R2-FKRP: TGGCACTTTTCGGGGGATCCTC

PALS-C-universal-R2-LARGE1: TGGCACTTTTCGGGGGATCCCT

All possible SNVs were generated and visualized using the following scripts:

All possible SNVs package

R1 oligos were generated using the following Python scripts:

PALS_C_oligos_FKRP.py

PALS_C_oligos_LARGE1.py

R1 oligo structure (5’->3’):

8-nt block-specific adaptor

Type2S enzyme recognition site

5-nt block-specific insulator (the 3’ 4-nt sequence does not appear in this block)

19-nt *GOI* downstream sequence

Variant

25-nt *GOI* upstream sequence

R1 oligo lists: FKRP_oligo_to_order_out.fa

LARGE1_oligo_to_order_out.fa

Plasmid templates:

Lenti-UbC-FKRP-EF1α-BSD

Lenti-UbC-LARGE1-EF1α-BSD

Reagents:

Q5 Reaction Buffer (NEB, B9027SVIAL)

Q5 High GC Enhancer (NEB, B9028AVIAL)

10 mM dNTPs (NEB, N0447)

Q5 High-Fidelity DNA Polymerase (NEB, M0491SVIAL)

BsmBI-v2 (NEB, R0739S)

BsaI-HFv2 (NEB, R3733S)

DpnI (NEB, R0176S)

MspJI (NEB, R0661S)

XbaI (NEB, R0145S)

BamHI-HF (NEB, R3136S)

NEBuilder HiFi DNA Assembly Master Mix (NEB, E2621)

Endura Electrocompetent Cells (Lucigen, 60242)

PALS-C Step1:

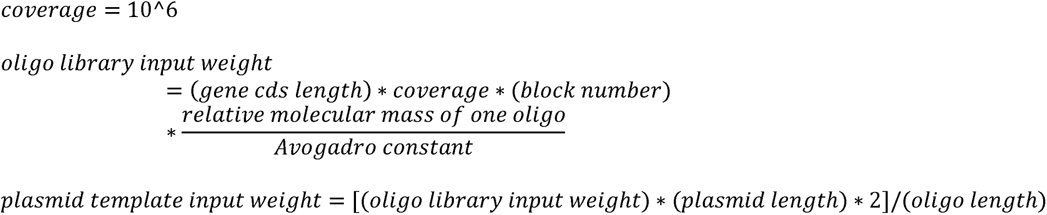

|  |  |
| --- | --- |
| Q5 Rxn buffer | 10 μL |
| Q5 Enh buffer | 10 μL |
| 10 mM dNTPs | 1 μL |
| Oligo library |  |
| Plasmid template |  |
| Q5 polymerase* | 2 μL |
| Water | To 50 μL* |

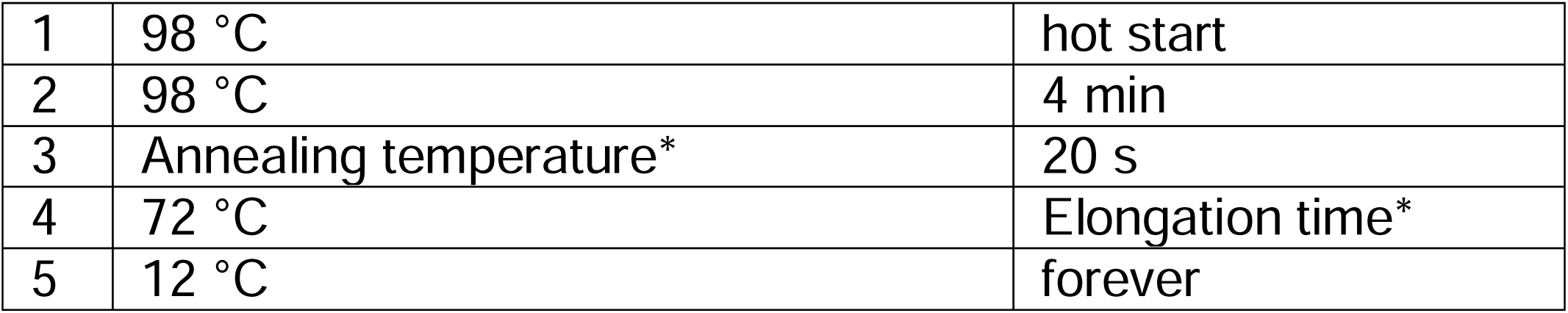

|  |  |  |
| --- | --- | --- |
| 1 | 98 °C | hot start |
| 2 | 98 °C | 4 min |
| 3 | Annealing temperature* | 20 s |
| 4 | 72 °C | Elongation time* |
| 5 | 12 °C | forever |

\*Multiple 50-μL reactions are needed if product yield is insufficient for subsequent steps.

*Q5 polymerase is likely to be the limiting factor of Step1, volume of which requires optimization.

*Multiple 50-μL reactions are needed if product yield is insufficient for subsequent steps.

*Annealing temperature and elongation time are decided based on the product that is the most difficult to amplify. One universal condition was feasible for FKRP (70 °C; 40 s) and LARGE1 (68 °C ; 90 s); however, if a GOI requires drastically different conditions for different blocks, reactions with different conditions can be set accordingly.

PCR purification with NucleoSpin Gel and PCR Clean-Up Kit (Takara, 740609).

PALS-C Step2:

Starting from Step2, the reactions are done independently for different blocks.

|  |  |
| --- | --- |
| Q5 Rxn buffer | 10 μL |
| Q5 Enh buffer | 10 μL |
| 10 mM dNTPs | 1 μL |
| Purified Step1 product | * |
| 10 μM Universal F1 | 2.5 μL |
| 10 μM Block specific R1 | 2.5 μL |
| Q5 polymerase | 0.5 μL |
| Water | To 50 μL |

|  |  |  |
| --- | --- | --- |
| 1 | 98 °C | hot start |
| 2 | 98 °C | 3 min |
| 3 | 98 °C | 8 s |
| 4 | Annealing temperature | 20 s |
| 5 | 72 °C | Elongation time |
| 6 | Repeat 3-5 for 34 more cycles |  |
| 7 | 72 °C | 5 min |
| 8 | 12 °C | Forever |

*The input should be decided based on the position of the block. The more distant a block is from the 5’ side, the more input is required. Evenly distributed input for all 6 blocks of FKRP generated enough yield for subsequent steps, while the 3’ side blocks of LARGE1 required extra input. Step1-2 should be repeated if product yield is insufficient for subsequent steps.

PCR purification with NucleoSpin Gel and PCR Clean-Up Kit (Takara, 740609).

PALS-C Step3:

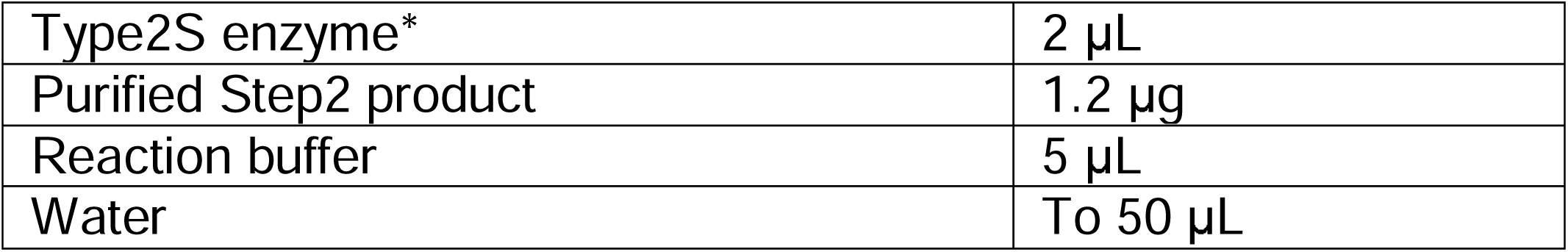

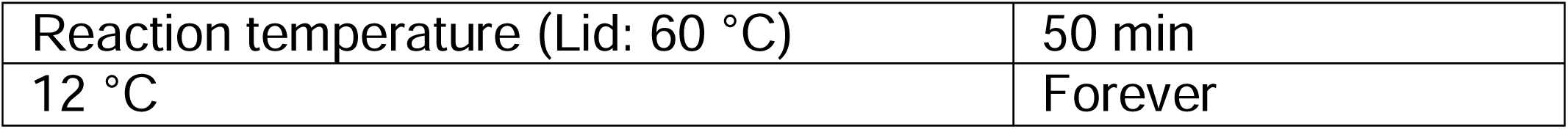

*BsmBI was used for FKRP and BsaI was used for LARGE1. The type2S enzymes were picked to avoid the presence of their recognition sites within the CDS.

Electrophoresis in 1% agarose gel was performed and the bands within the correct size range were cut from the gel; gel purification with NucleoSpin Gel and PCR Clean-Up Kit (Takara, 740609).

PALS-C Step4:

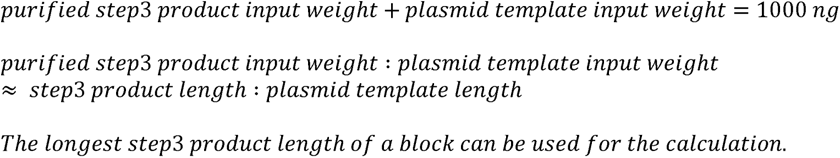

|  |  |
| --- | --- |
| Q5 Rxn buffer | 10 μL |
| Q5 Enh buffer | 10 μL |
| 10 mM dNTPs | 1 μL |
| Purified Step3 product |  |
| Plasmid template |  |
| Q5 polymerase* | 2 μL |
| Water | To 50 μL |

|  |  |  |
| --- | --- | --- |
| 1 | 98 °C | hot start |
| 2 | 98 °C | 5 min |
| 3* | 72 °C | 5 s |
| 4* | 66 °C | 20 s |
| 5 | 72 °C | Elongation time* |
| 6 | 12 °C | forever |

\*Optional steps: the purpose is to enhance the annealing of all strands.

*Q5 polymerase is likely to be the limiting factor of Step4, volume of which requires optimization.

*Optional steps: the purpose is to enhance the annealing of all strands.

*The elongation time should be sufficient for the shortest strand to be elongated to the R2 primer site. PCR purification with NucleoSpin Gel and PCR Clean-Up Kit (Takara, 740609).

PALS-C Step5:

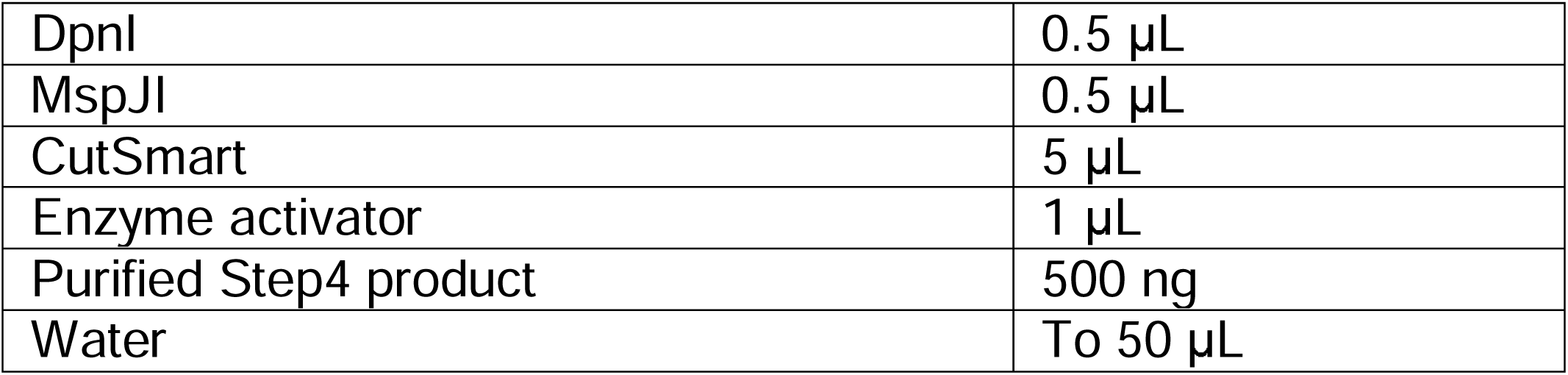

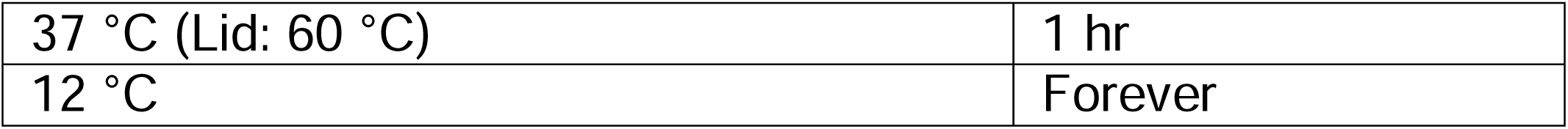

Column purification with NucleoSpin Gel and PCR Clean-Up Kit (Takara, 740609). (Follow the PCR purification protocol.)

PALS-C Step6:

|  |  |
| --- | --- |
| Q5 Rxn buffer | 10 μL |
| Q5 Enh buffer | 10 μL |
| 10 mM dNTPs | 1.5 μL |
| Purified Step5 product | 100 ng |
| 10 μM F2 | 2.5 μL |
| 10 μM R2 | 2.5 μL |
| Q5 polymerase | 1 μL |
| Water | To 50 μL |

|  |  |  |
| --- | --- | --- |
| 1 | 98 °C | hot start |
| 2 | 98 °C | 5 min |
| 3 | 98 °C | 6 s |
| 4 | Annealing temperature | 20 s |
| 5 | 72 °C | Elongation time |
| 6 | Repeat 3-5 for 34 more cycles |  |
| 7 | 72 °C | 5 min |
| 8 | 12 °C | Forever |

Electrophoresis in 1% agarose gel was performed and the bands with the correct size were cut from the gel; gel purification with NucleoSpin Gel and PCR Clean-Up Kit (Takara, 740609).

Important: Use 20 μL water or less to dissolve after gel purification; or use a vacuum concentrator to evaporate excess water.

PALS-C Step7:

Prepare the backbone:

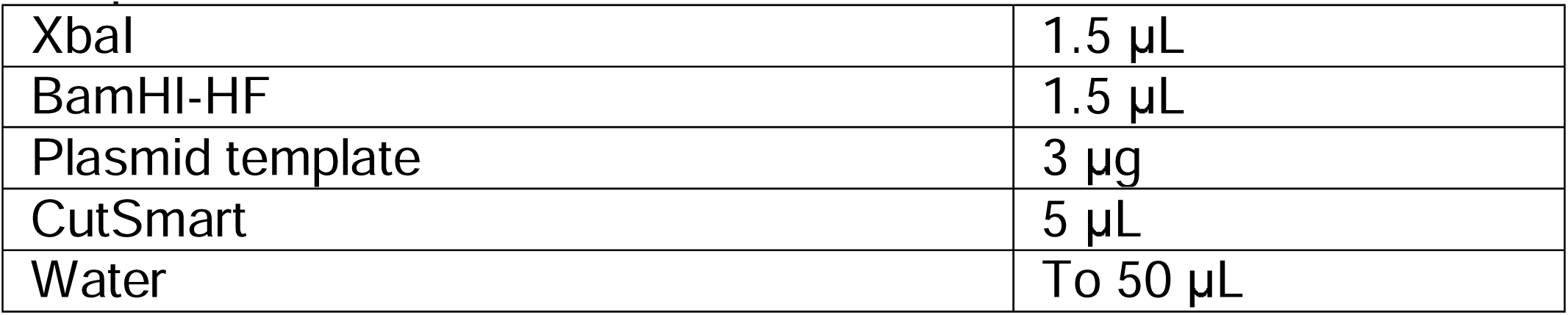

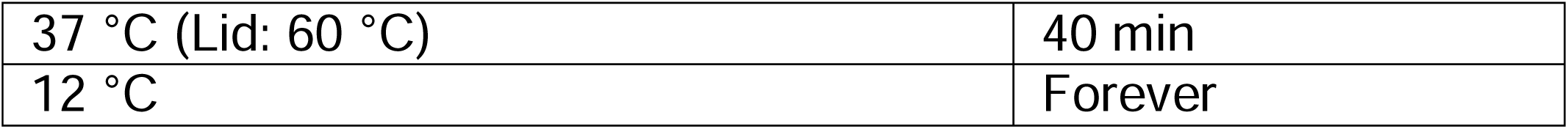

Electrophoresis in 1% agarose gel was performed and the band with the correct size was cut from the gel; gel purification with NucleoSpin Gel and PCR Clean-Up Kit (Takara, 740609).

For Gibson assembly:

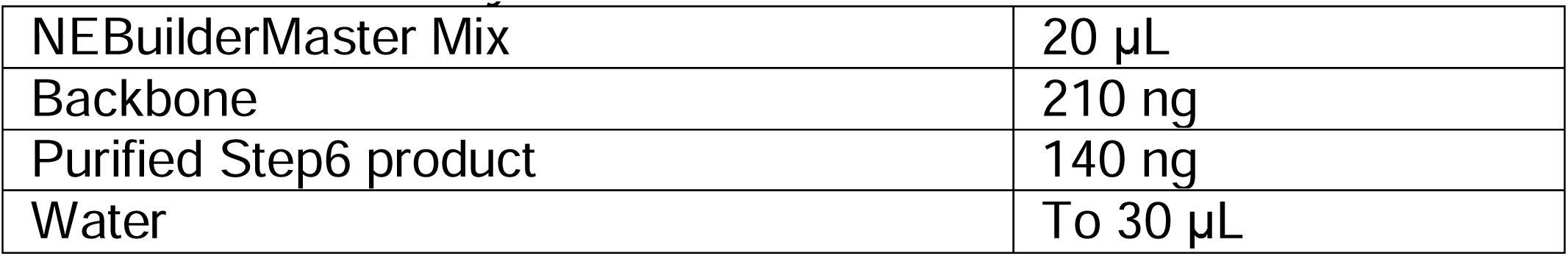

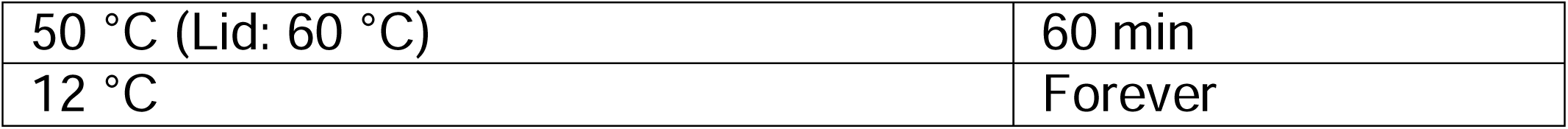

PALS-C Step8:

For each block:

|  |  |
| --- | --- |
| Electrocompetent cells | 40 µL |
| Assembly reaction | 4 µL |
| Water | 160 µL |

Split the mixed sample into two pre-chilled 0.1 cm cuvettes (Bio-Rad, 1652089).

Use Bio-Rad Gene Pulser II: (ATTENTION: avoid bubbles) 25 μF; 200 Ohms; 1800 volts. Add the sample to 900 μL recovery media per cuvette: 250 rpm, 1hr, 37°C.

Combine transformed bacteria from 2 cuvettes.

Add 1/500 volume of the bacteria to 200 μL LB broth and plate it on an ampicillin LB agar plate for quick estimation of complexity. (We achieved 19.1 × to 115.3 x coverage for all blocks)

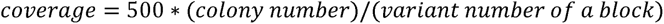

Seed the bacteria in 150 mL LB broth supplemented with ampicillin (100 µg/ml) for overnight culture.

## Method S5

Scripts used to list underrepresented variants due to the limitation of PALS-C: Cloning alert package

Group1 variants introduce a cut before the variants which will remove the variant from the top strand; Group2 variants introduce a cut after the variants which might affect the elongation of the top strand.

FKRP:

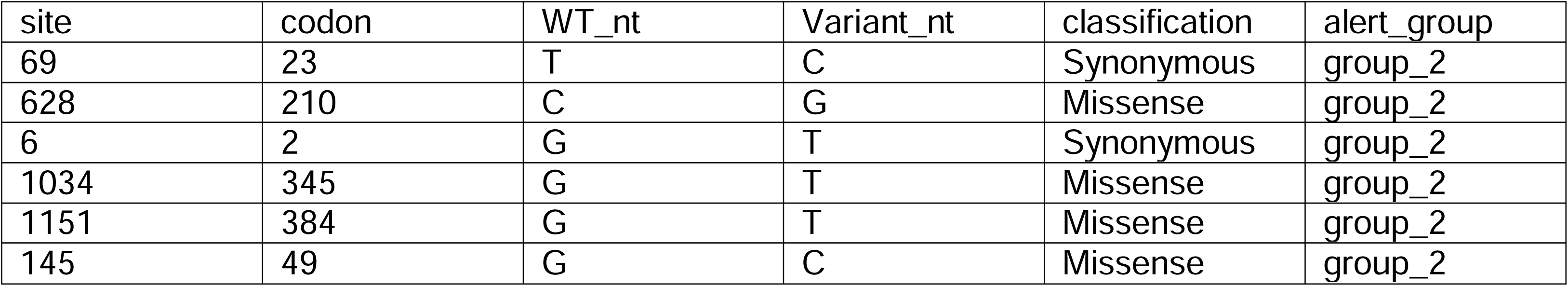

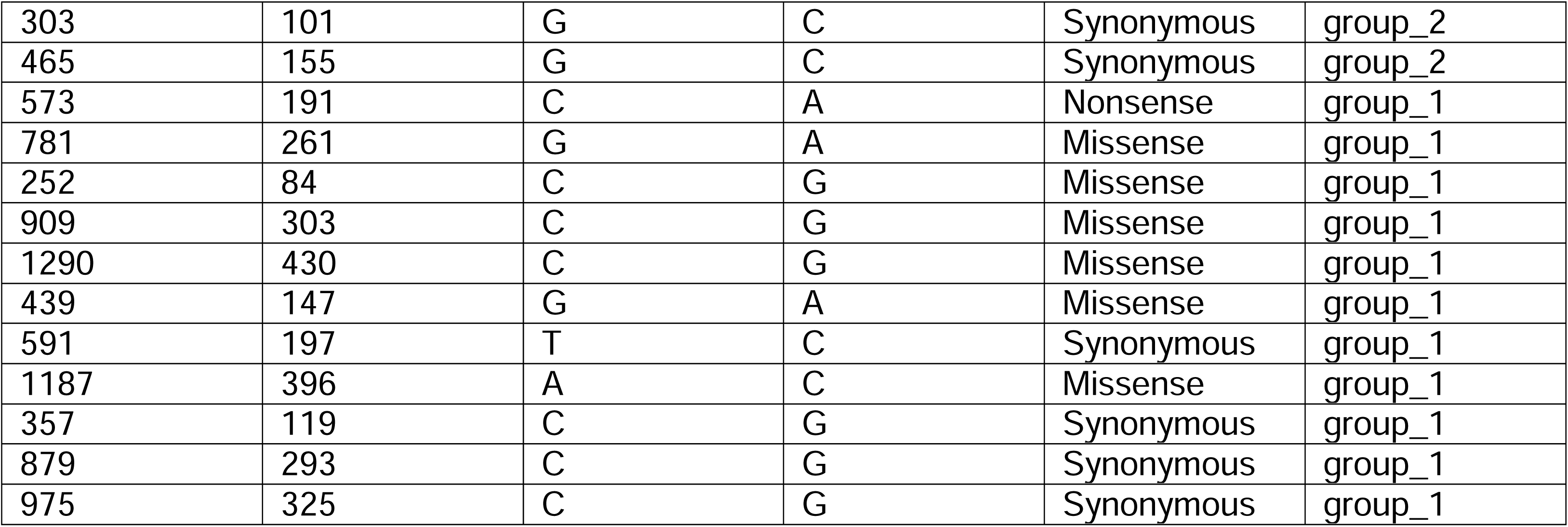

LARGE1:

| site | codon | WT_nt | Variant_nt | classification | alert_group |
| --- | --- | --- | --- | --- | --- |
| 119 | 40 | T | G | Missense | group_2 |
| 2176 | 726 | T | G | Missense | group_2 |
| 982 | 328 | C | G | Missense | group_2 |
| 167 | 56 | C | T | Missense | group_2 |
| 1556 | 519 | G | T | Missense | group_2 |
| 1713 | 571 | G | T | Synonymous | group_2 |
| 1432 | 478 | G | T | Missense | group_2 |
| 666 | 222 | G | C | Synonymous | group_2 |
| 298 | 100 | A | G | Missense | group_1 |
| 559 | 187 | C | G | Missense | group_1 |
| 1082 | 361 | C | G | Nonsense | group_1 |
| 1859 | 620 | G | A | Missense | group_1 |
| 838 | 280 | A | G | Missense | group_1 |
| 139 | 47 | T | A | Missense | group_1 |
| 269 | 90 | C | A | Missense | group_1 |
| 1468 | 490 | G | A | Missense | group_1 |
| 1528 | 510 | G | A | Missense | group_1 |
| 1963 | 655 | G | A | Missense | group_1 |
| 191 | 64 | G | C | Missense | group_1 |
| 227 | 76 | A | C | Missense | group_1 |
| 338 | 113 | A | C | Missense | group_1 |
| 815 | 272 | A | C | Missense | group_1 |
| 1347 | 449 | G | C | Missense | group_1 |
| 2261 | 754 | A | C | Missense | group_1 |
| 26 | 9 | G | C | Missense | group_1 |
| 331 | 111 | A | C | Missense | group_1 |
| 411 | 137 | A | C | Synonymous | group_1 |
| 1117 | 373 | G | C | Missense | group_1 |
| 1797 | 599 | A | C | Synonymous | group_1 |
| 1994 | 665 | G | C | Missense | group_1 |

## Method S6

Primers used for plasmid pool QC:

FKRP-QC-block1-F: GCCAGAACACAGGACCGGTTCTAGA

FKRP-QC-block1-R: GCCAGGGGCGGGTAGGGGAGCGTGT

FKRP-QC-block2-F: GCCCAGCCCGTGGTGGTGGCAGCCG

FKRP-QC-block2-R: CAGGCTGACGTTCAGGGCCAGGCAC

FKRP-QC-block3-F: CGGTTGCCACGGCCAACCCTGCCAG

FKRP-QC-block3-R: GCTCAGCCTTCCAGCGCGCGTGGGC

FKRP-QC-block4-F: GGCGCGCCAGCCCCCGCTGGCCACG

FKRP-QC-block4-R: AGCCAGTAGCGCACGCCCGCAGCCT

FKRP-QC-block5-F: GCCCGCTATGTGGTGGGCGTGCTGG

FKRP-QC-block5-R: GCCATTGCGGGGGTAGAAGGGCCAC

FKRP-QC-block6-F: AAAGCAACCACTTGCACGTGGACCT

FKRP-QC-block6-R: GAGAGAAGTTTGTTGCGCCGGATCC

LARGE1-NGS-blk1-QC-F: CCCTACACGACGCTCTTCCGATCTCCTACGTCACTTGGTACCGGTTCTAGA

LARGE1-NGS-blk2-QC-F: CCCTACACGACGCTCTTCCGATCTCCTACGTCGCGAGGTGGAGGAGGAGAA

LARGE1-NGS-blk3-QC-F: CCCTACACGACGCTCTTCCGATCTCCTACGTCGGATACAATGCCAGCCGGG

LARGE1-NGS-blk4-QC-F: CCCTACACGACGCTCTTCCGATCTCCTACGTGTCTGATGAAGCTTGTCCTG

LARGE1-NGS-blk5-QC-F: CCCTACACGACGCTCTTCCGATCTCCTACGTACAGGGGTGATCCTGTTACT

LARGE1-NGS-blk6-QC-F: CCCTACACGACGCTCTTCCGATCTCCTACGTCGTGTCTGATCTAAAGGTCA

LARGE1-NGS-blk7-QC-F: CCCTACACGACGCTCTTCCGATCTCCTACGTGAGAGCGCTTCACTGTCCAC

LARGE1-NGS-blk8-QC-F: CCCTACACGACGCTCTTCCGATCTCCTACGTATGAGCCGCCACAACGTGGG

LARGE1-NGS-blk9-QC-F: CCCTACACGACGCTCTTCCGATCTCCTACGTACTGCGCTACCGGCTGTCCT

LARGE1-NGS-blk10-QC-F: CCCTACACGACGCTCTTCCGATCTCCTACGTTAGGCTTTGGCTGGAACAAA

LARGE1-NGS-blk1-R: GACTGGAGTTCAGACGTGTGCTCTTCCGATCTGCTGCCTGCGGAGGGCGCGG

LARGE1-NGS-blk2-R: GACTGGAGTTCAGACGTGTGCTCTTCCGATCTTTTGACCAGGGTGACGACAT

LARGE1-NGS-blk3-R: GACTGGAGTTCAGACGTGTGCTCTTCCGATCTTTGGCAGGAAGAGTCTTGGT

LARGE1-NGS-blk4-R: GACTGGAGTTCAGACGTGTGCTCTTCCGATCTTCTTCCGCAGCTTATCCAGA

LARGE1-NGS-blk5-R: GACTGGAGTTCAGACGTGTGCTCTTCCGATCTCTTGGGGGAGTTCCAGTGAA

LARGE1-NGS-blk6-R: GACTGGAGTTCAGACGTGTGCTCTTCCGATCTAGGAAGTACAGGTGGGTGCG

LARGE1-NGS-blk7-R: GACTGGAGTTCAGACGTGTGCTCTTCCGATCTCCTTGTACACGATGTGGTAG

LARGE1-NGS-blk8-R: GACTGGAGTTCAGACGTGTGCTCTTCCGATCTCTCCGCTTTTGACTTGGGGA

LARGE1-NGS-blk9-R: GACTGGAGTTCAGACGTGTGCTCTTCCGATCTAGCTCCATGATATGAGCCAC

LARGE1-NGS-blk10-R: GACTGGAGTTCAGACGTGTGCTCTTCCGATCTCACTTTTCGGGGGATCCCTA

NGS-PCR3-F: AATGATACGGCGACCACCGAGATCTACACTCTTTCCCTACACGACGCTCTTCCGATCT

NGS-PCR3-R: CAAGCAGAAGACGGCATACGAGATCGCGCGGTGTGACTGGAGTTCAGACGTGTGCTCTT

For *FKRP* plasmid pool QC:

For the plasmid pool of each block:

|  |  |
| --- | --- |
| Q5 Rxn buffer | 10 µL |
| Q5 Enh buffer | 10 µL |
| 10 mM dNTPs | 1 µL |
| Plasmid | ~300 ng |
| 10 µM F primer | 2.5 µL |
| 10 µM R primer | 2.5 µL |
| Q5 polymerase | 0.5 µL |
| Water | To 50 µL |

|  |  |  |
| --- | --- | --- |
| 1 | 98 °C | hot start |
| 2 | 98 °C | 3 min |
| 3 | 98 °C | 6 s |
| 4 | 70 °C | 15 s |
| 5 | 72 °C | 5 s |
| 6 | Repeat 3-5 for 32 more cycles |  |
| 7 | 72 °C | 5 min |
| 8 | 12 °C | Forever |

Electrophoresis and gel purification (Takara, 740609) were performed.

The purified products were sent for Amplicon-EZ sequencing.

For *LARGE1* plasmid pool QC:

For the plasmid pool of each block:

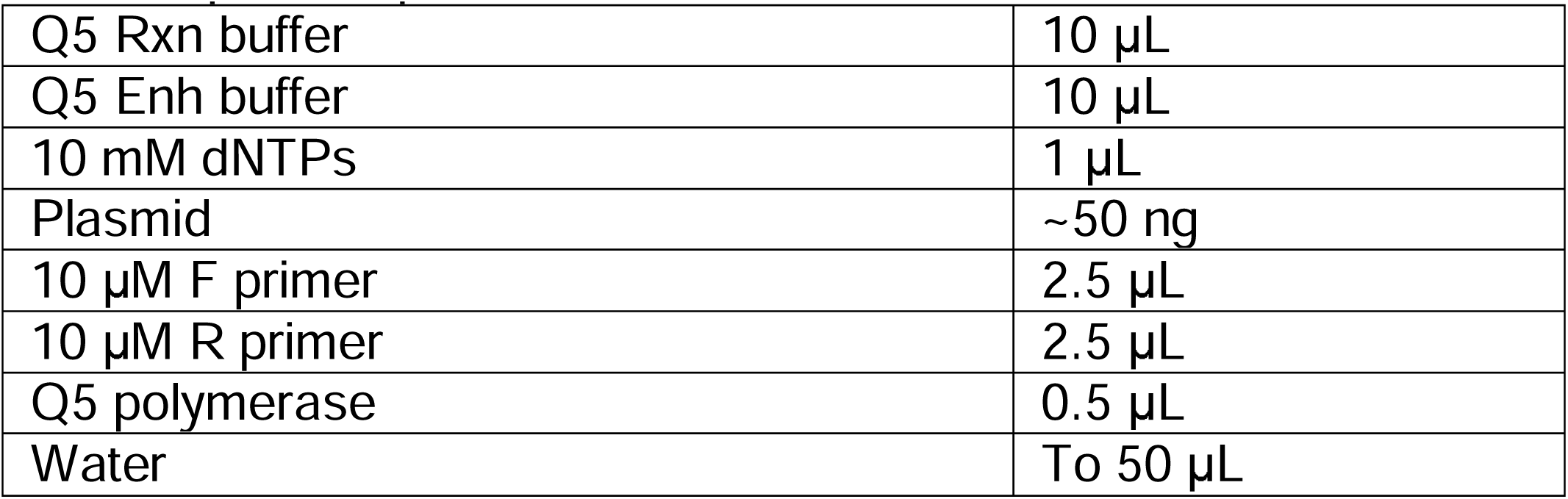

|  |  |
| --- | --- |
| Q5 Rxn buffer | 10 µL |
| Q5 Enh buffer | 10 µL |
| 10 mM dNTPs | 1 µL |
| Plasmid | ~50 ng |
| 10 µM F primer | 2.5 µL |
| 10 µM R primer | 2.5 µL |
| Q5 polymerase | 0.5 µL |
| Water | To 50 µL |

|  |  |  |
| --- | --- | --- |
| 1 | 98 °C | hot start |
| 2 | 98 °C | 3 min |
| 3 | 98 °C | 6 s |
| 4 | Annealing temperature | 15 s |
| 5 | 72 °C | 7 s |
| 6 | Repeat 3-5 for 33 more cycles |  |
| 7 | 72 °C | 5 min |
| 8 | 12 °C | Forever |

|  |  |
| --- | --- |
| Annealing temperature | Blocks |
| 61 °C | 1, 4, 6, 7 |
| 64 °C | 5, 9, 10 |
| 66 °C | 2, 3, 8 |

Electrophoresis and gel purification (Takara, 740609) were performed. The purified products were mixed for the following reaction:

|  |  |
| --- | --- |
| Q5 Rxn buffer | 10 µL |
| Q5 Enh buffer | 10 µL |
| 10 mM dNTPs | 1 µL |
| Mixed purified products | 100 ng |
| 10 µM NGS-PCR3-F | 2.5 µL |
| 10 µM NGS-PCR3-R | 2.5 µL |
| Q5 polymerase | 1 µL |
| Water | To 50 µL |

|  |  |  |
| --- | --- | --- |
| 1 | 98 °C | hot start |
| 2 | 98 °C | 3 min |
| 3 | 98 °C | 6 s |
| 4 | 72 °C | 15 s |
| 5 | 72 °C | 8 s |
| 6 | Repeat 3-5 for 19 more cycles |  |
| 7 | 72 °C | 5 min |
| 8 | 12 °C | Forever |

PCR purification (Takara, 740609) was performed and the sample was sent for Amplicon-EZ sequencing.

Scripts for the plasmid pool QC analytical pipelines:

FKRP plasmid QC

package LARGE1 plasmid QC package

## Method S7

Variants in the mini-libraries/used for individual validation:

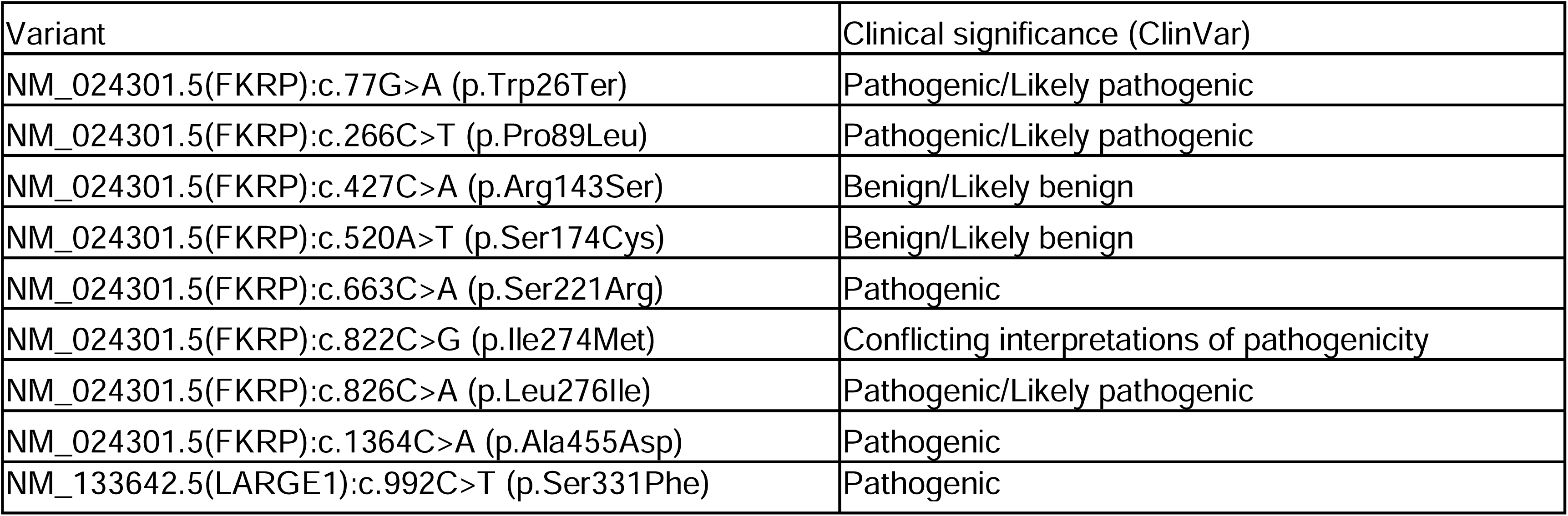

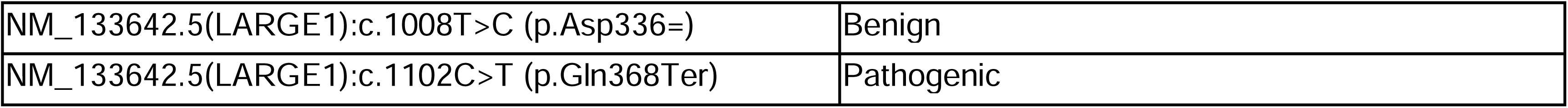

Primers used for lentiviral sequence isolation:

PALS-C-universal-F2: GATCGTCACTTGGTACCGGTTCTAGA

PALS-C-universal-R2-FKRP: TGGCACTTTTCGGGGGATCCTC

PALS-C-universal-R2-LARGE1: TGGCACTTTTCGGGGGATCCCT

Primers used for *FKRP* Sanger sequencing:

FKRP-QC-block2-F: GCCCAGCCCGTGGTGGTGGCAGCCG

FKRP-QC-block3-F: CGGTTGCCACGGCCAACCCTGCCAG

FKRP-QC-block5-F: GCCCGCTATGTGGTGGGCGTGCTGG

FKRP-GT-R: CATCAGGTACTAGGGCCACAAACTC

Primers used for *FKRP* Amplicon-EZ sequencing:

Amplicon1:

PALS-C-universal-F2: GATCGTCACTTGGTACCGGTTCTAGA FKRP-GT-R: CATCAGGTACTAGGGCCACAAACTC

Amplicon2:

FKRP-QC-block2-F: GCCCAGCCCGTGGTGGTGGCAGCCG FKRP-QC-block3-R: GCTCAGCCTTCCAGCGCGCGTGGGC

Amplicon3:

FKRP-QC-block3-F: CGGTTGCCACGGCCAACCCTGCCAG FKRP-QC-block3-R: GCTCAGCCTTCCAGCGCGCGTGGGC

Amplicon4:

FKRP-QC-block4-F: GGCGCGCCAGCCCCCGCTGGCCACG FKRP-QC-block4-R: AGCCAGTAGCGCACGCCCGCAGCCT

Amplicon5:

FKRP-QC-block6-F: AAAGCAACCACTTGCACGTGGACCT

FKRP-CDS-backbone-junction-R: GTGGCACTTTTCGGGGGATCCTCAGCCGCTTCCCGTCAGACTC

Primers used for *LARGE1* Sanger sequencing:

LARGE1-block4-seq-F: TTATGGTCTGATGAAGCTTGTCCTG

## Method S8

PCR1:

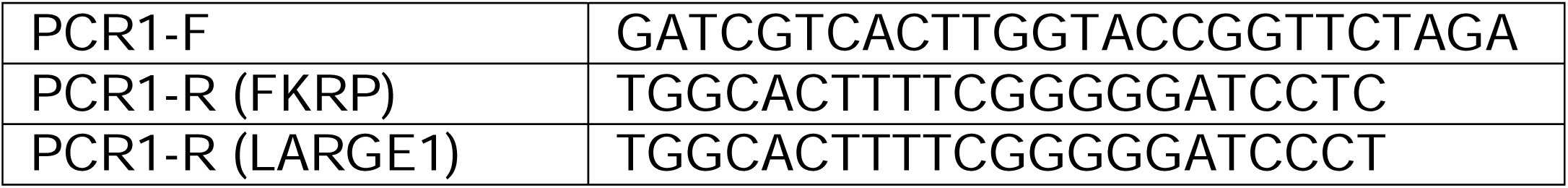

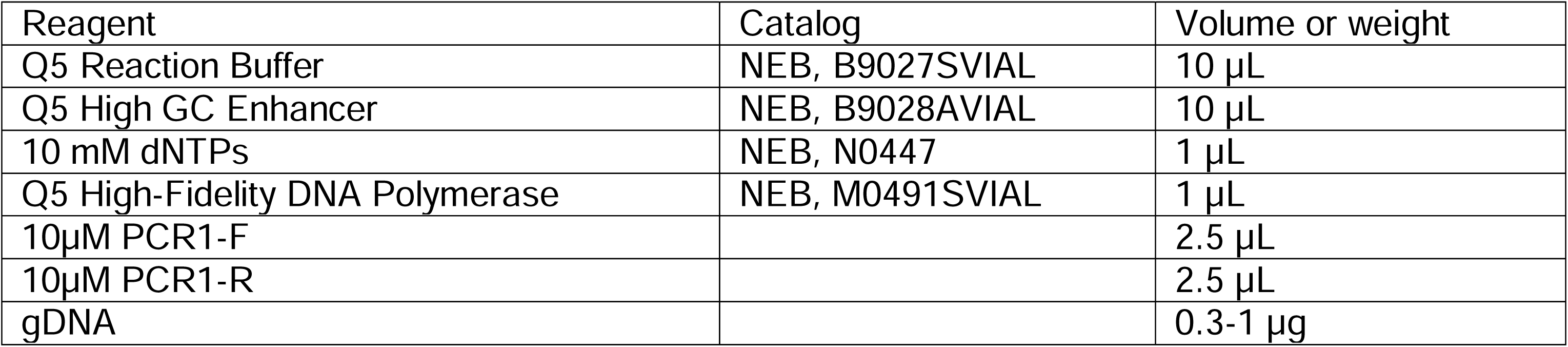

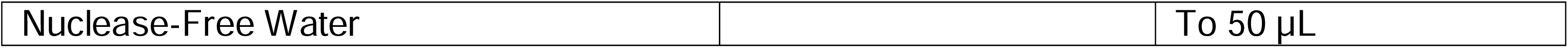

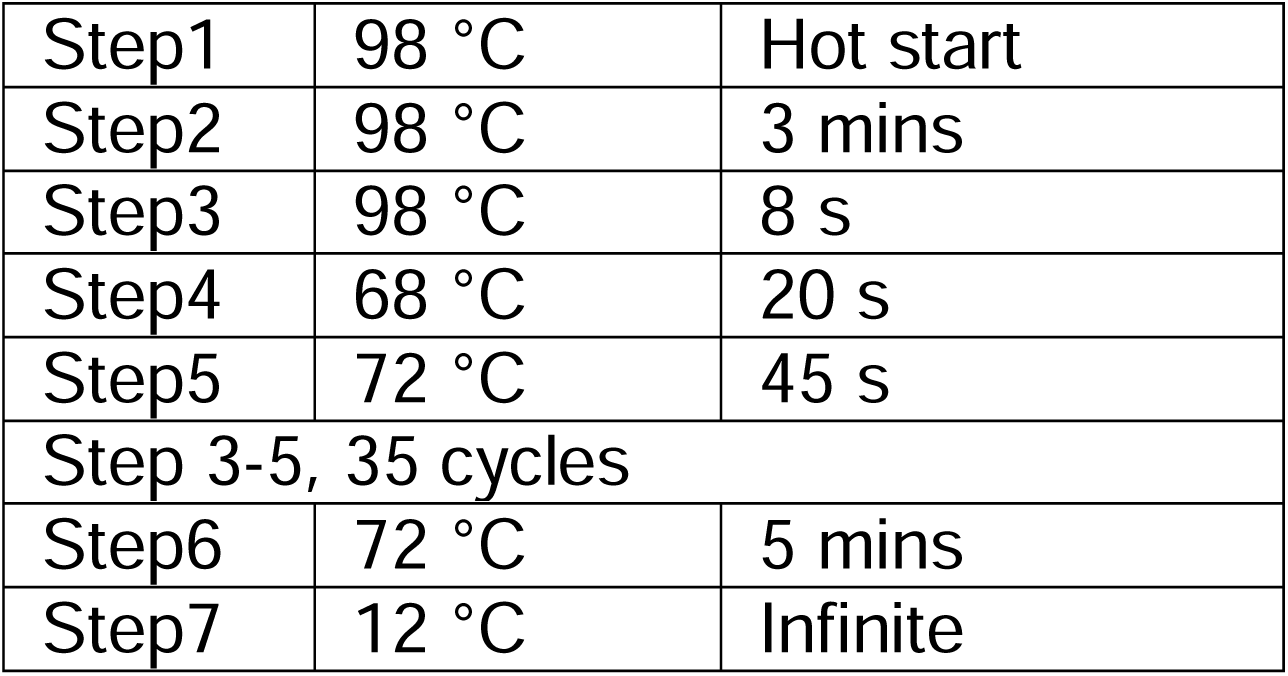

Electrophoresis in 1% agarose gel. Product band size is 1534 bp (FKRP) or 2317 bp (LARGE1). Gel purification with NucleoSpin Gel and PCR Clean-Up Kit (Takara, 740609). Elute with 25 µL Nuclease-Free Water.

PCR2:

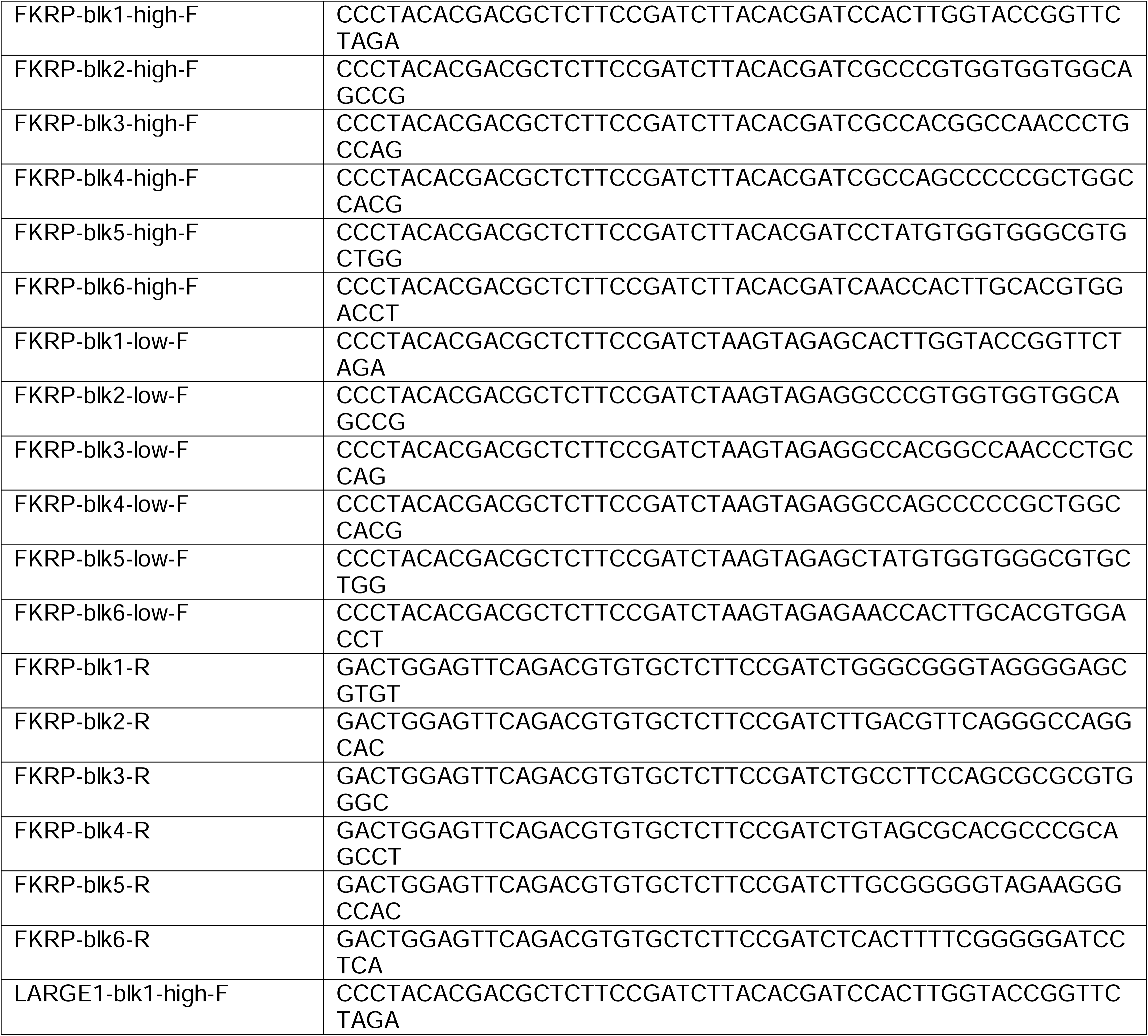

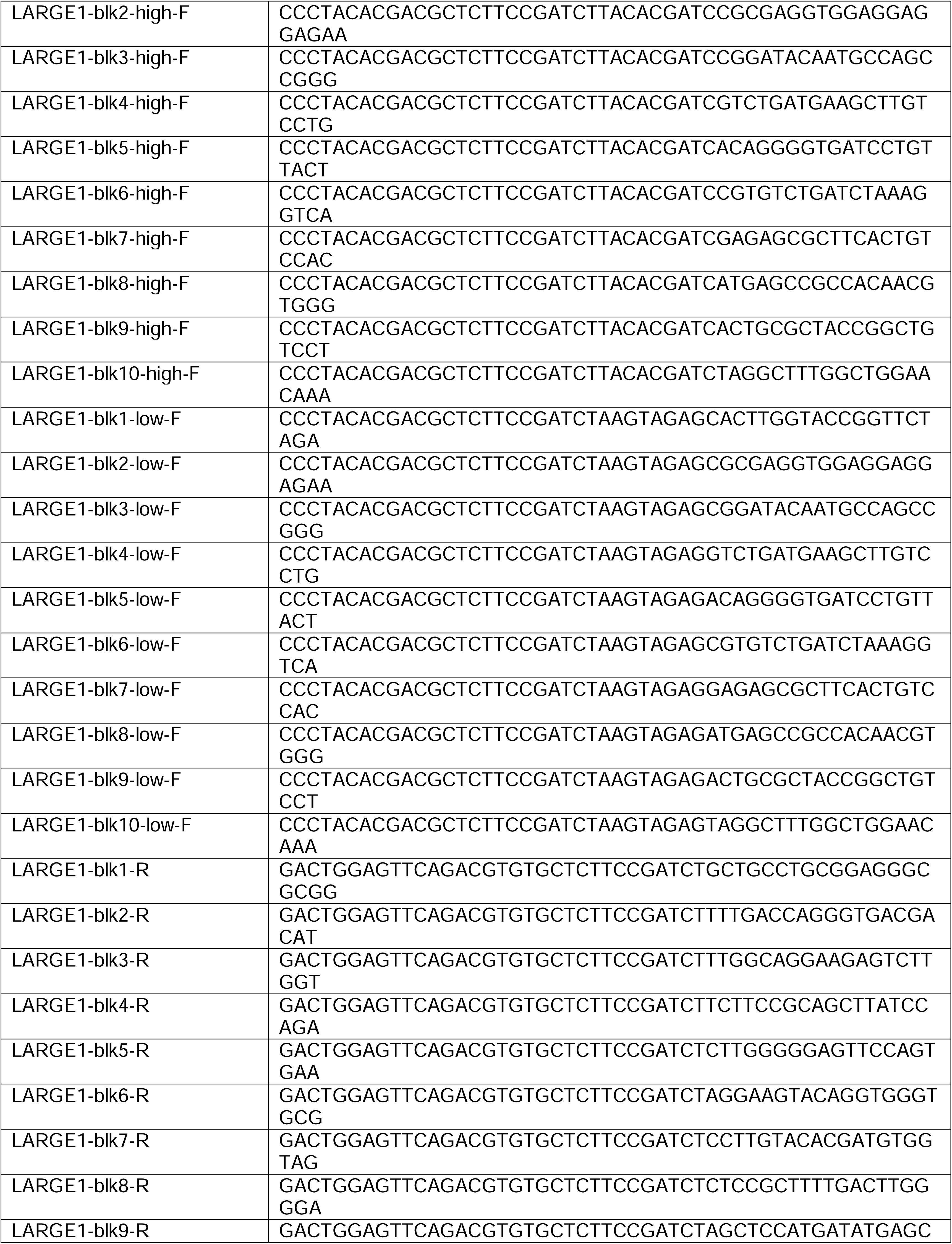

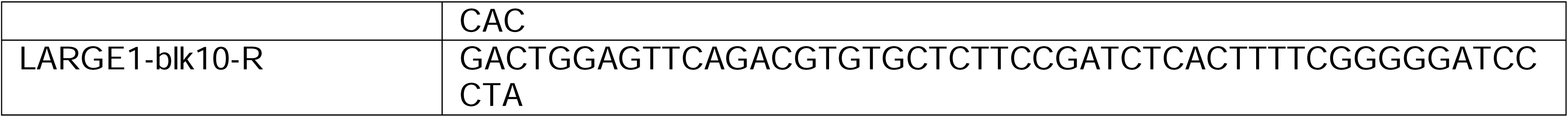

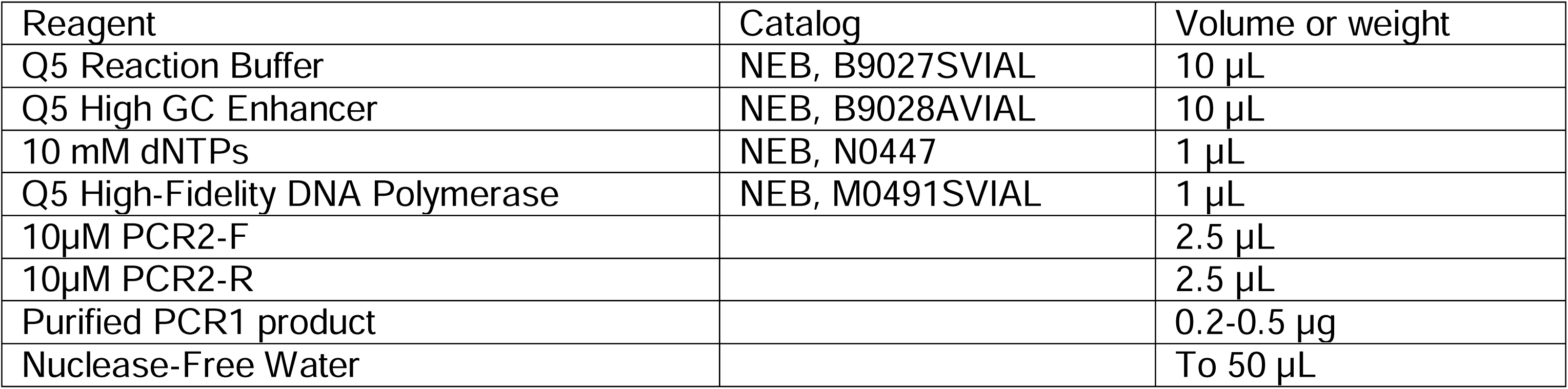

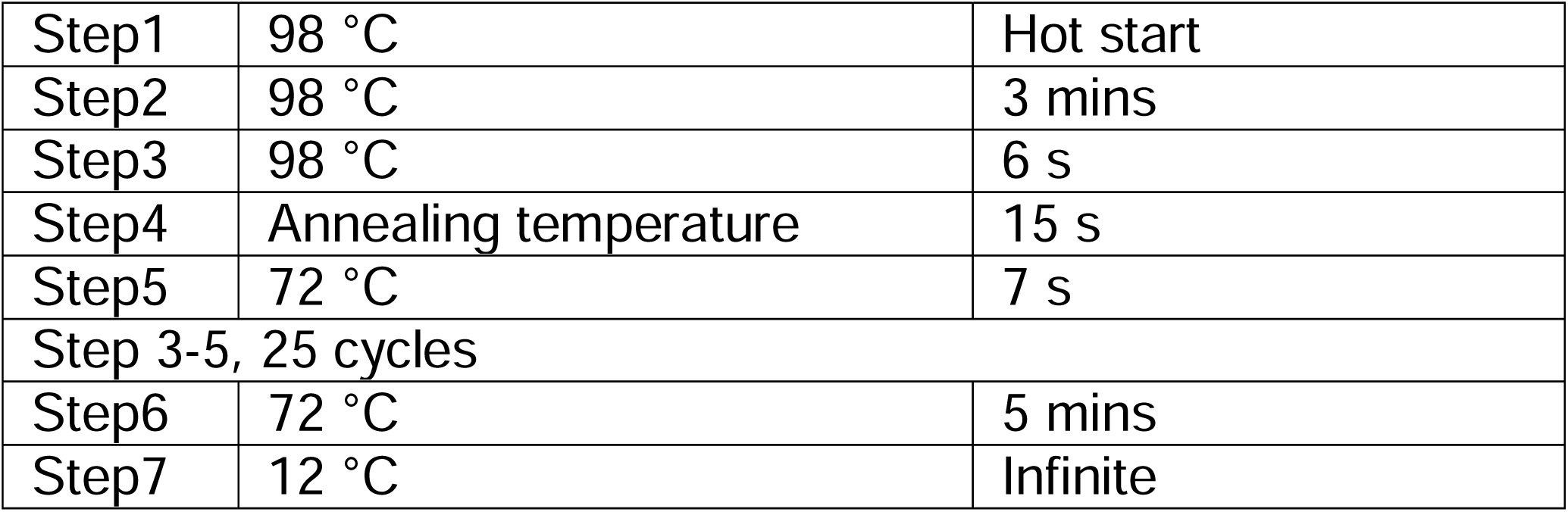

Annealing temperature:

|  |  |
| --- | --- |
| 61 °C | LARGE1-blk1, LARGE1-blk4, LARGE1-blk6, LARGE1-blk7 |
| 64 °C | FKRP-blk1, LARGE1-blk5, LARGE1-blk9, LARGE1-blk10 |
| 66 °C | LARGE1-blk2, LARGE1-blk3, LARGE1-blk8 |
| 68 °C | FKRP-blk6 |
| 72 °C | FKRP-blk2, FKRP-blk3, FKRP-blk4, FKRP-blk5 |

PCR purification with NucleoSpin Gel and PCR Clean-Up Kit (Takara, 740609). Elute with 40 µL Nuclease-Free Water.

PCR3:

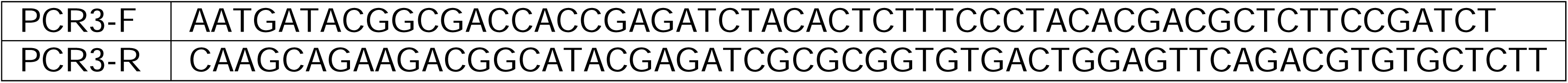

Mix purified PCR2 products (200 ng each). Dilute the mixed sample to 11 ng/µL.

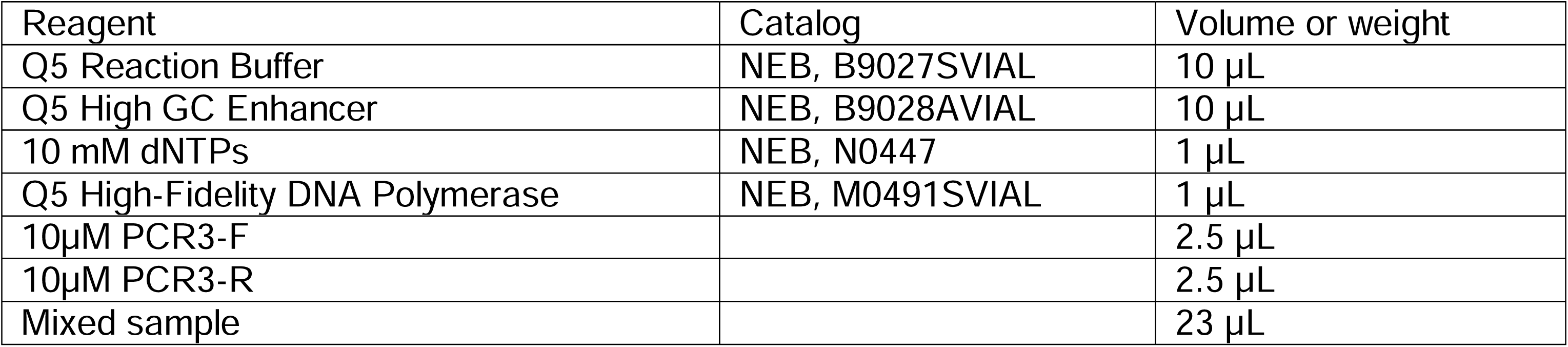

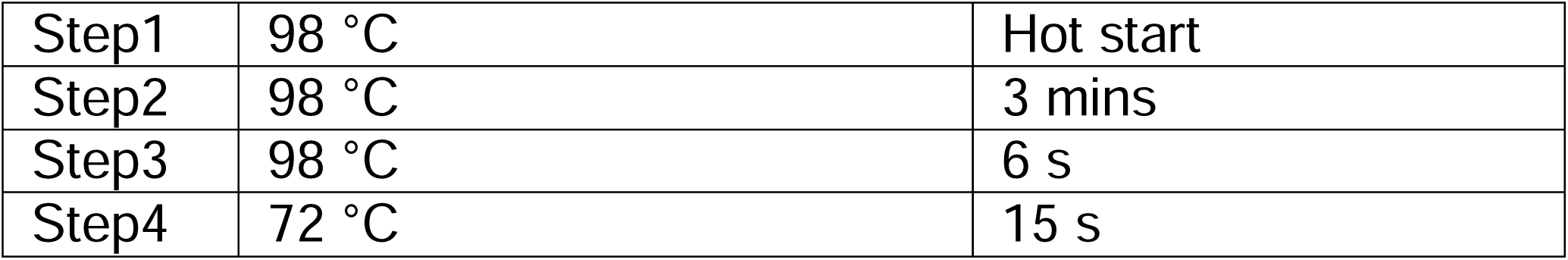

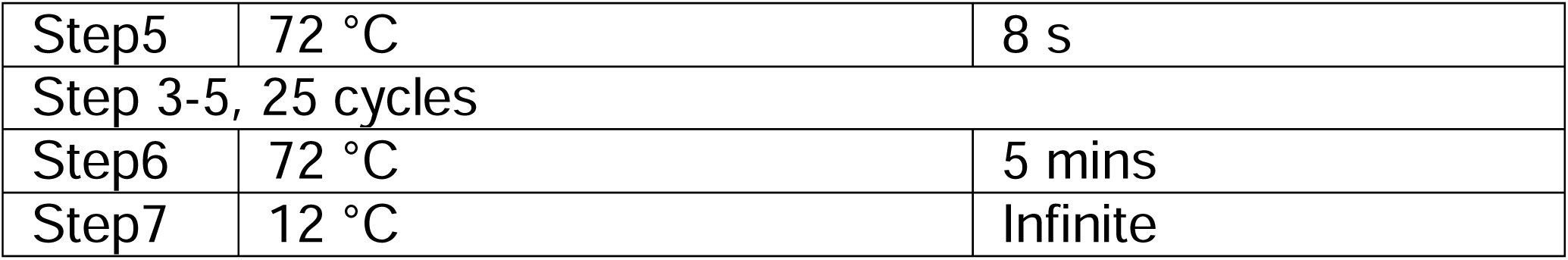

Set 3*50 µL reactions.

PCR purification with NucleoSpin Gel and PCR Clean-Up Kit (Takara, 740609). Elute with 50 µL Nuclease-Free Water for each column. The purified PCR3 product was used for the next-generation sequencing. Amplicon-EZ service (GENEWIZ) was used to check the sample quality and coverage. The sample was then sequenced using the HiSeq X service (Psomagen; size selection also performed there).

## Method S9

Protein extraction and quantitation:

Ice-cold protein extraction lysis buffer made by adding 10 μL of protease inhibitor cocktail (Sigma Aldrich, P8340-1ML) to every 1 mL of RIPA lysis and extraction buffer (Pierce, PI89900) was used to treat the cells. After 15 minutes of rocking incubation on ice, the samples were centrifuged for 30 minutes at 16,000 × g at 4 °C in a refrigerated centrifuge (Eppendorf, 5404). The supernatant was transferred to a clean 1.5 mL microcentrifuge tube for the following steps.

Protein was quantitated using the BioRad DC Assay II kit (BioRad, 5000112). Using the BSA reagent from the kit, seven standards are prepared from 0–1.5 mg/mL in the protein extraction lysis buffer from above to create a standard curve. Measurements were done according to the instruction manual, and absorbances at 750nm were read on a plate reader (BioRad, #1681135). Concentration is calculated relative to the standard curve.

SDS-PAGE and western blotting:

Appropriate amount of protein extraction lysis buffer from above was added into each protein sample (∼10 µg) to balance the samples to achieve an equal volume. 4x Laemlli (BioRad, 1610747) was supplemented with 10% b-mercaptoethanol. 1 µL of this supplemented buffer was then added to every 3 µL of the balanced sample. Samples were mixed by a brief vortex and then heated at ∼95°C on a heat block (BioRad, 1660571) for 5 minutes. Samples were cooled down on ice and then centrifuged briefly at max speed to ensure equal loading (Eppendorf, 5404).

3.5 μL of PageRuler Plus prestained protein ladder (ThermoFisher Scientific, 26619) and all the samples were then loaded onto a 4-15% mini-protean TGX stain free protein gel (*e.g.*, BioRad, 4568084) that had been placed into a Mini-PROTEAN tetra cell electrophoresis system (BioRad, 1658000). SDS-PAGE was performed at 120V for ∼60 minutes using a PowerPac HC power supply (BioRad, 1645052). Upon completion of the SDS-PAGE, the gel was removed and transferred onto a nitrocellulose membrane (BioRad, 1704158) by the TransBlot Turbo System (BioRad, 1704150EDU) using the 7-minute transfer program (2.5 A constant, up to 25 V variable).

Vinculin was used as a loading control. Hence, the membrane was cut at the correct position for separate FKRP and vinculin WBs. Membrane blocking buffer was made of 5% Omniblock non-fat dry milk (American Bio, AB10109-00100) in 1× TBS-Tween 20 (TBS-T). The membrane pieces were then blocked for 1 hour in membrane-blocking buffer separately.

Primary Anti-FKRP antibody (Abcam, ab220059) was made up 1:1,000 in membrane-blocking buffer. Primary Anti-Vinculin antibody (Sigma Aldrich, V9131-100 µl) was made up 1:80,000 in membrane-blocking buffer. The membrane pieces were incubated in the corresponding primary antibodies overnight at 4°C whilst rocking to equally distribute antibody (BenchRocker 2D Genesee Scientific, 31-201).

The following day, the membrane pieces were washed 5 times for 5 minutes in TBS-T and incubated in corresponding secondary antibodies for 2 hours at room temperature. Anti-rabbit HRP conjugated secondary antibody (Cell Signaling Technology, 7074S) was used for FKRP at 1:2,000 in membrane-blocking buffer. Anti-mouse HRP conjugated secondary antibody (Cell Signaling Technology, 7076S) was used for vinculin at 1:2,000 in membrane-blocking buffer.

Membrane pieces were washed again 5 times for 5 mins at room temperature in TBS-T prior to developing with Clarity Max ECL substrate (BioRad, 1705062) or Clarity ECL substrate (BioRad, 1705060). Using the Chemidoc Imaging system (BioRad, 12005606), both chemiluminescent and colorimetric images were taken to observe protein expression and size of bands.

## Method S10

Packaging ppVSV-LASV-GPC viral particles:

General information:

* The LASV-GPC plasmid and the ppVSVDG-VSV-G viral particles were obtained from Dr. Melinda Brindley’s group.

* The ppVSVDG-VSV-G viral particles carry a GFP-coding sequence in their genomes, which can be used to determine the TCID50 (tissue culture infectious dose that will infect 50% of the cell monolayers).

* TCID50 of the viral particles was experimentally determined using a protocol provided by Dr. Melinda A. Brindley. The protocol was based on the Spearman-Karber method.

* To calculate the multiplicity of infection (MOI) based on TCID50, a working estimate was adopted:

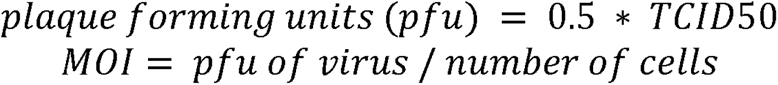

* Exercise standard caution when handling viral particles!

* Biosafety Level 2 (BSL-2) for activities with materials and cultures known or reasonably expected to contain VSV.

Making ppVSV-LASV-GPC-Generation1:

|  |  |
| --- | --- |
| Day 0 | Seed HEK293T cells in a well of a 6-well plate. |
| Day 1 | Perform Lipofectamine 3000 (Invitrogen, L3000001) transfection per manufacturer's instruction. |
|  | *HEK293T should reach 70%-90% confluency. |
|  | *4 µg LASV-GPC plasmid was used to transfect the cells. |
| Day 2 | 24 hrs after transfection, ppVSVDG-VSV-G (MOI=0.5) was added to the well. |
|  | The cell number is estimated to be ~2 M. Use this number to decide the viral dose. |
|  | 1 hr after adding the ppVSVDG-VSV-G, the medium was removed, the well was washed with DPBS (Gibco, 14190144), and fresh medium was added to the well. |
| Day 3 | Collect the newly generated viral particles, *i.e.*, ppVSV-LASV-GPC-Generation1, and titrate its titer. |

Making ppVSV-LASV-GPC-Generation2:

|  |  |
| --- | --- |
| Day 0 | Seed 6M HEK293T cells in a 10-cm dish. |
| Day 1 | Perform Lipofectamine 3000 (Invitrogen, L3000001) transfection per manufacturer's instruction. |
|  | *30 µg LASV-GPC plasmid was used to transfect the cells. |
| Day 2 | 24 hrs after transfection, ppVSV-LASV-GPC-Generation1 (MOI=0.1) was added to the well. |
|  | The cell number is estimated to be ~12 M. Use this number to decide the viral dose. |
|  | 1 hr after adding the ppVSV-LASV-GPC-Generation1, the medium was removed, the well was washed with DPBS (Gibco, 14190144), and fresh medium was added to the well. |
| Day 3 | Collect the newly generated viral particles, *i.e.*, ppVSV-LASV-GPC-Generation2, and titrate its titer. |

Using ppVSV-LASV-GPC-Generation2, ppVSV-LASV-GPC-Generation3 can be packaged, similar to ppVSV-LASV-GPC-Generation2.

NGS library construction for VSV-related experiments was performed using a procedure similar to that outlined in Method S8. The "high" primers were utilized for the infected groups, while the primers provided below were used for the non-infected groups.

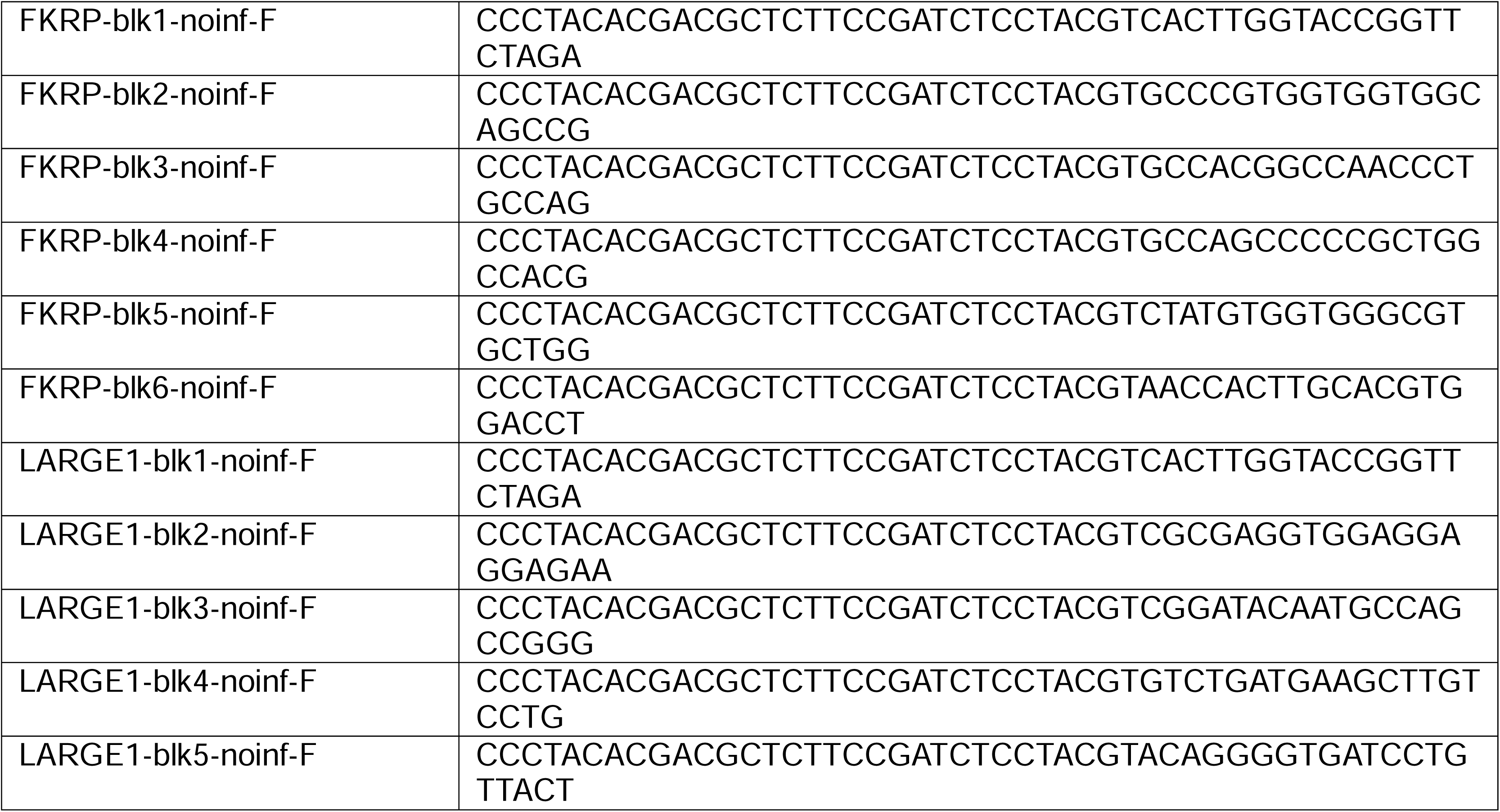

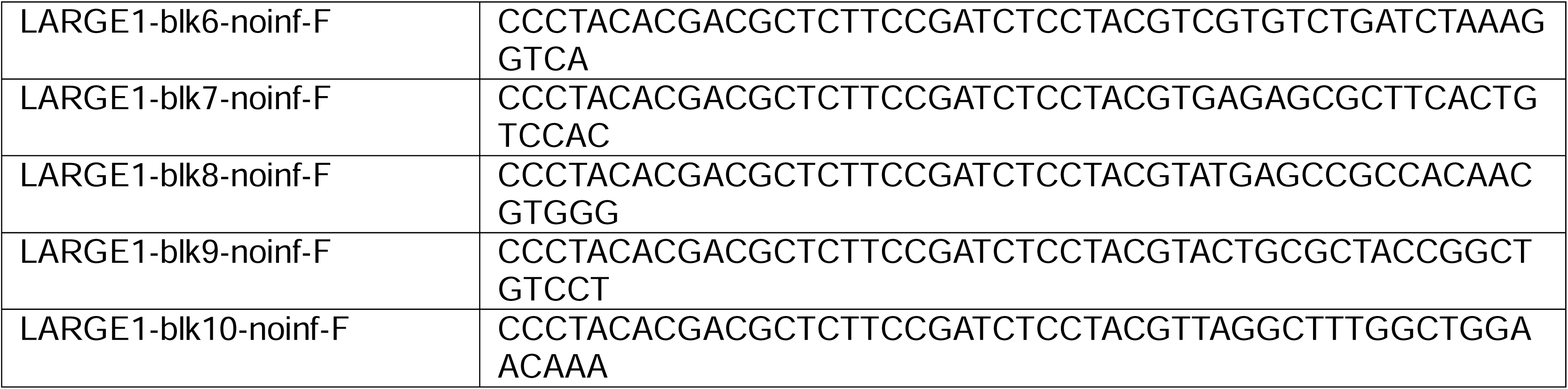

Considerations for ppVSV application:

In our experiment, we noticed that the Lenti-*LARGE1* rescued *LARGE1*-KO MB135 appeared to be more sensitive to the ppVSV infection when compared to the Lenti-*FKRP* rescued *FKRP*-KO MB135. For *FKRP* application, the ppVSV MOI should be controlled to be approximated 2.5, while for *LARGE1* application, the MOI should be around 1. For other genes, we recommend testing the MOI in a range of 1-3.

Lentiviral MOI, ppVSV MOI, and cell confluency can all affect the sensitivity of the ppVSV assay. We recommend starting the optimization with a low lentiviral MOI (<1) and a 20-40% confluency.

## Method S11

Variants in the mini-libraries:

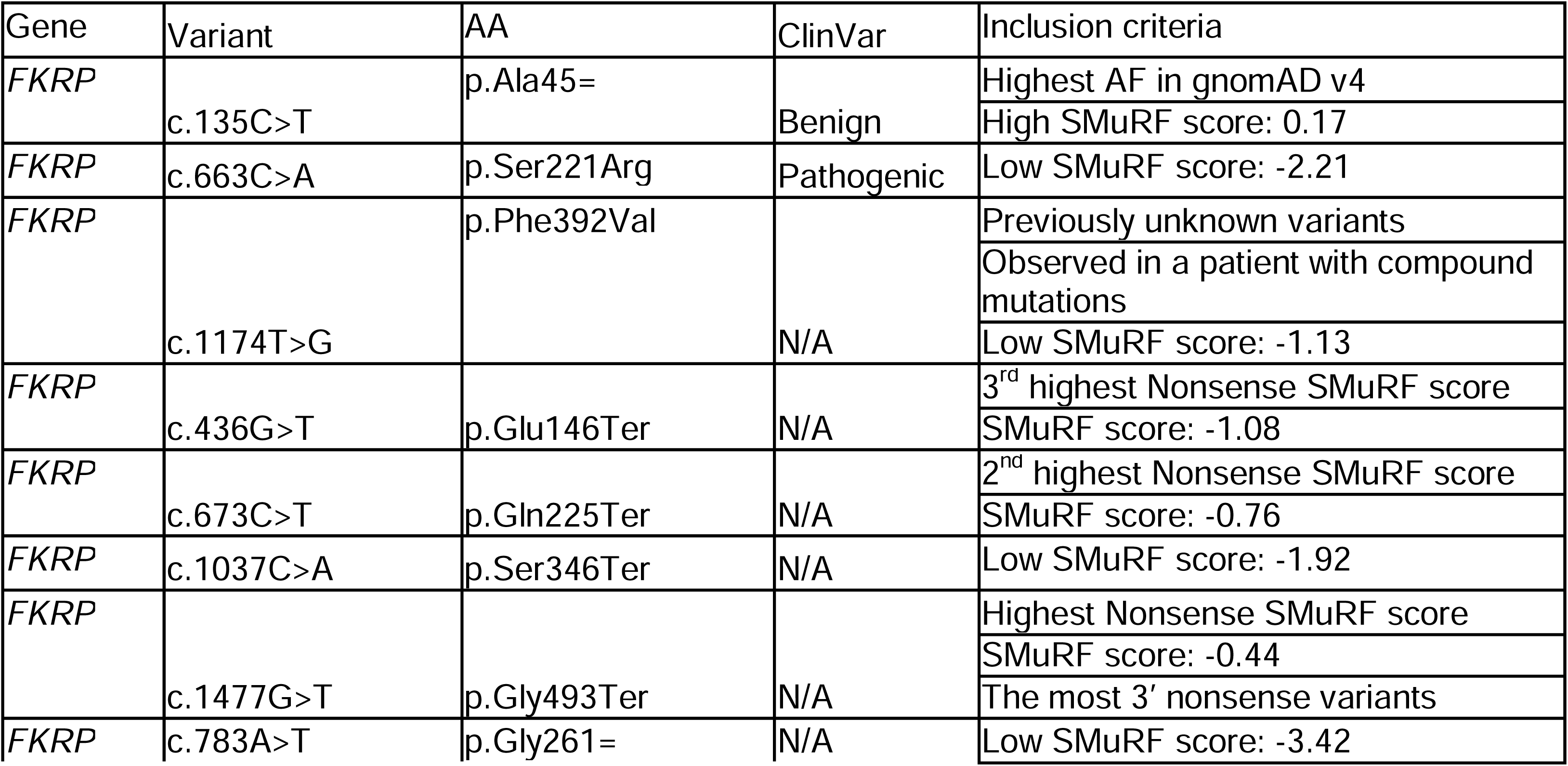

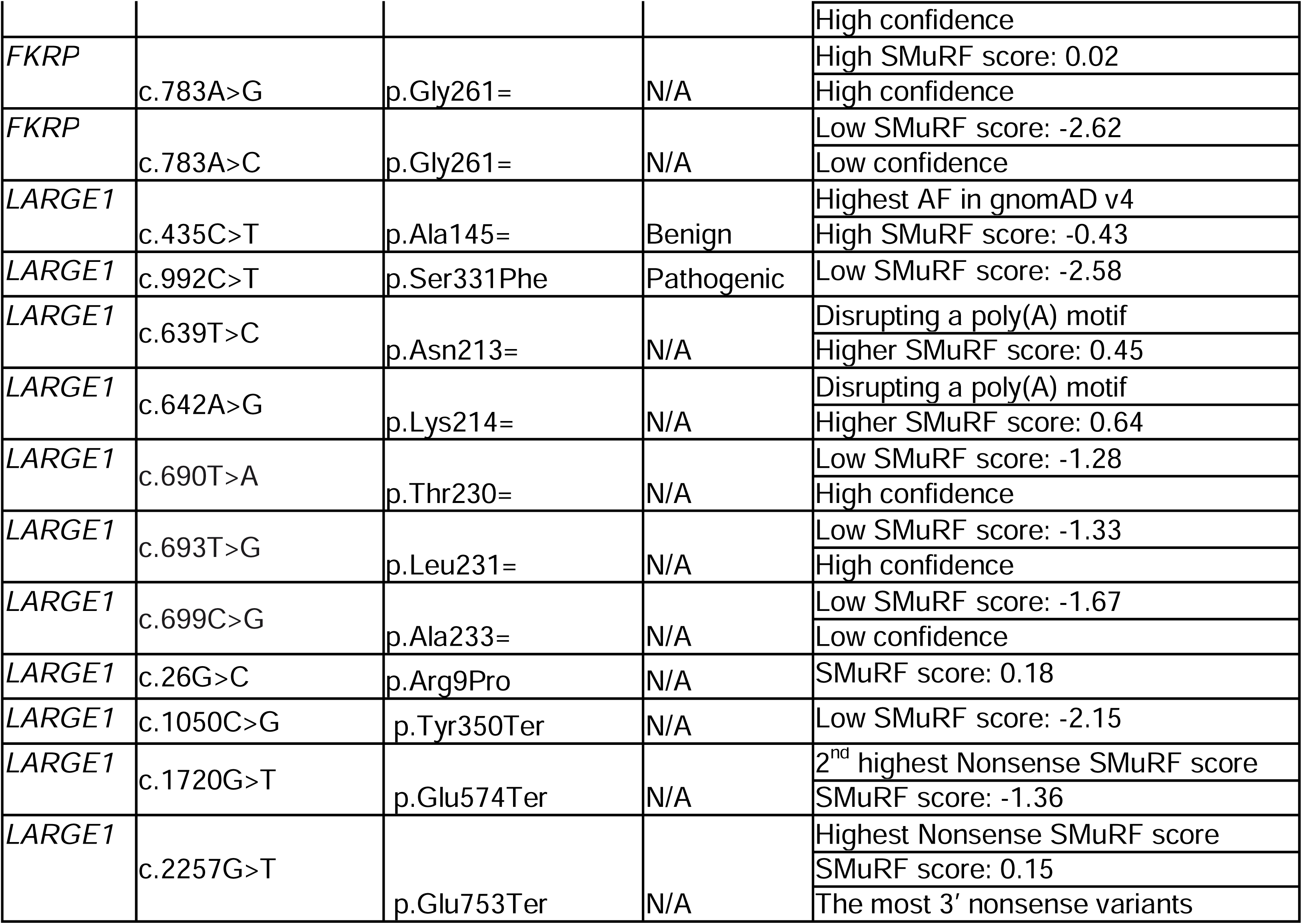

Analytical script example: plasmidsaurus_enrichment_calculation_v2.py

Considerations for nanopore sequencing:

Reproducible high noise may arise at certain sites due to the nature of nanopore sequencing. It is recommended to ensure that such noise does not impact the enrichment calculations. If it does, consider conducting short-read sequencing instead.

---

**Figure S1:**
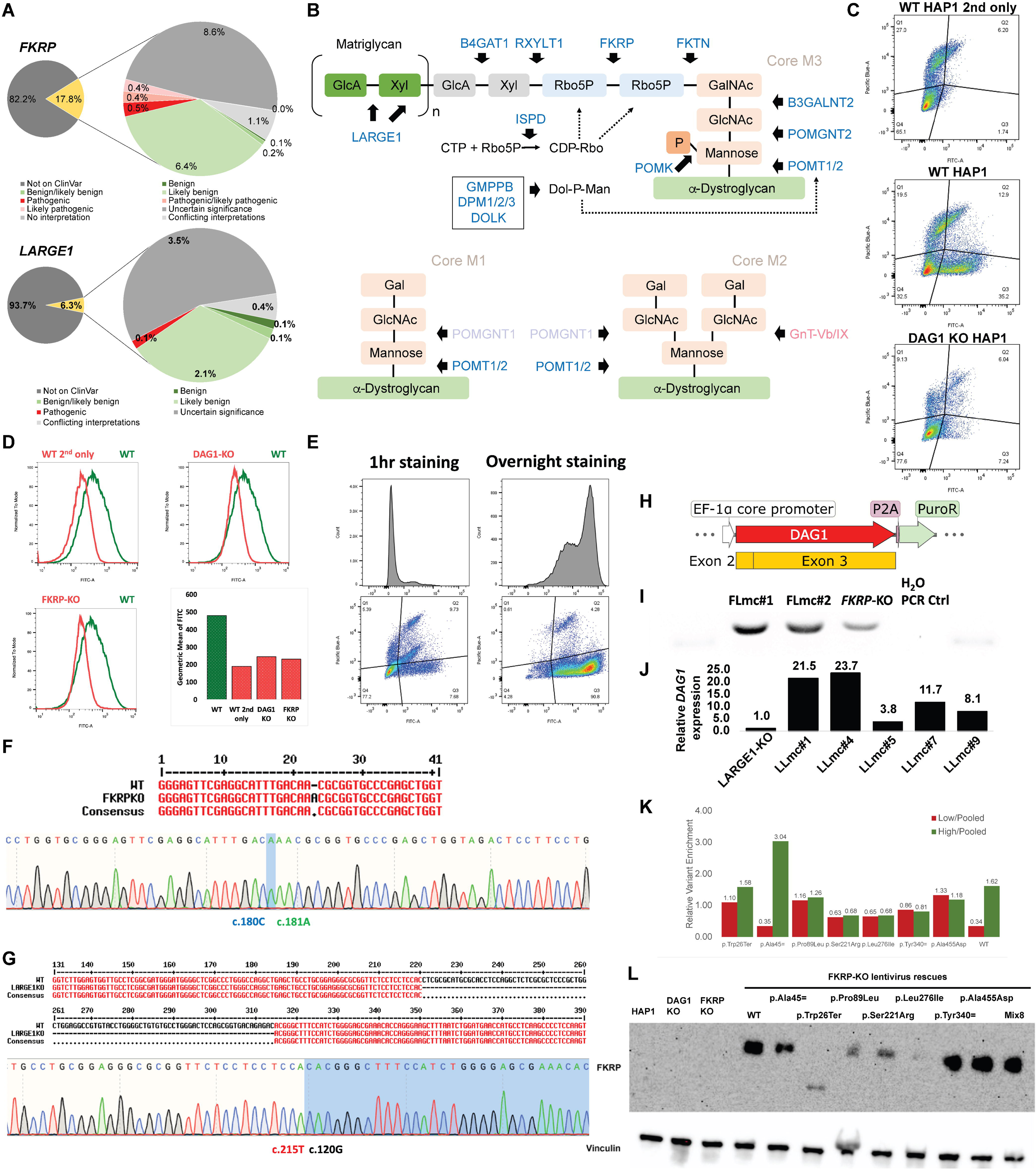
**The background and preparation of the implementation of SMuRF** (A) The majority of possible coding SNVs of α-DG glycosylation enzyme-coding genes lack clinical interpretation (*e.g.*, no ClinVar record, April 20, 2023). Around half of the reported SNVs of these genes in ClinVar were classified as variants of uncertain significance. (B) Functions of most α-DG glycosylation enzymes can be evaluated by the IIH6C4 antibody (blue texts). Bold arrows link enzymes to the modifications or glycan additions they catalyze; for instance, POMK catalyzes mannose phosphorylation. POMT1/2 participate the glycosylation of α-DG Core M1, M2 and M3. POMGNT1 mutations in patients manifest the hallmark of reduced IIH6C4 signal. GnT-Vb/IX mutations are not known to alter IIH6C4 signals. Hence, POMGNT1 but not GnT-Vb/IX can be evaluated by IIH6C4. Dol-P-Man, dolichol phosphate mannose; GlcNAc, N-acetylglucosamine; GalNAc, N-acetylgalactosamine; Rbo5P, ribitol-5-phosphate; Xyl, xylose; GlcA, glucuronic acid; Gal, galactose. (C) IIH6C4 fluorescence flow cytometry is compatible with the HAP1 platform. 1-hr room temperature staining was performed for both primary and secondary antibodies. The *DAG1*-KO HAP1 carries a 1-bp deletion (c.205Gdel). *FKRP*-KO HAP1 was established from a single clone of CRISPR RNP nucleofected cells and carries a 1-bp insertion (c.181Adup). FITC-A: IIH6C4; Pacific Blue-A: Viobility 405/452 Fixable Dye. (D) 1-hr staining is not sensitive enough for a high-throughput FACS-based variant characterization, as the separation between positive signal and null signal is unsatisfactory. (E) Optimized 4 °C overnight staining increased the assay sensitivity. Both staining protocols used the same samples: Lenti-*DAG1 FKRP*-KO HAP1 rescued by a mixture of Lenti-EF1a-*FKRP* and Lenti-EF1a-*FKRP*(c.826C>A). (F and G) *GOI*-KO HAP1 monoclonal cell lines were isolated from pooled CRISPR RNP nucleofected cells. *FKRP*-KO HAP1 carries a 1-bp frameshifting insertion (c.181Adup). *LARGE1*-KO HAP1 carries a 94-bp frameshifting deletion (c.121_214del). The mutations were validated with Sanger sequencing. (H) α-DG overexpression was achieved with Lenti-*DAG1*. (I) RT-PCR experiment for *FKRP*-KO Lenti-*DAG1* monoclonal lines (FLmc#1 and FLmc#2). The same amount of RNA molecules was used for each reaction. Bands in lane 2-5 were the PCR products of the *DAG1* cDNA primers (225 bp). Bands in lanes 1&6 were 200-bp ladder bands. FLmc#1 was chosen as the cell line platform for downstream experiments. (J) RT-qPCR experiment for *LARGE1*-KO Lenti-*DAG1* monoclonal lines. *HPRT1* primers were used as the housekeeping control for the ddCt quantification. The expression level was normalized to the *LARGE1*-KO sample. qPCR triplicates were set for each condition. LLmc#1 was chosen as the cell line platform for downstream experiments. (K and L) These results were from experiments in the early development stage. The platform cell line is the *FKRP*-KO cell line without Lenti-*DAG1* transduction. Initially, we constructed Lenti-EF1a-*FKRP*-P2A-BSD. We later abandoned this design as the over-expression failed to generate the expected separation and the P2A fusion impeded our ability to study nonsense variants in an unbiased manner. Interestingly, the cells treated with the lentivirus carrying the p.Trp26Ter nonsense mutation unexpectedly survived the blasticidin drug selection, suggesting the existence of a genetic mechanism that could rescue this nonsense mutation, at least in the context of using this construct. (K) A mini-library comprising 7 *FKRP* variants and the WT sequence was employed in a FACS experiment. The over-expression masked the deleterious effects of the pathogenic variants, leading to a lack of significant enrichment differences of those variants between the high-glycosylation and low-glycosylation populations. (The enrichment was quantified by Sanger sequencing peaks; the enrichment of the WT was calculated based on the enrichments of other variants.) (L) Western blot was performed for individually transduced samples as well as the Mix8 sample. The lentiviral expression was excessively high compared to the endogenous *FKRP* expression in WT HAP1 (not detectable by WB). A smaller band (between 35 kDa and 55 kDa) was detected in the p.Trp26Ter sample, supporting the hypothesis of the existence of a downstream alternative start codon rescuing this mutation (likely, Met144, resulting in a 39.0 kDa product). The WB protocol is described in Method S9.

**Figure S2:**
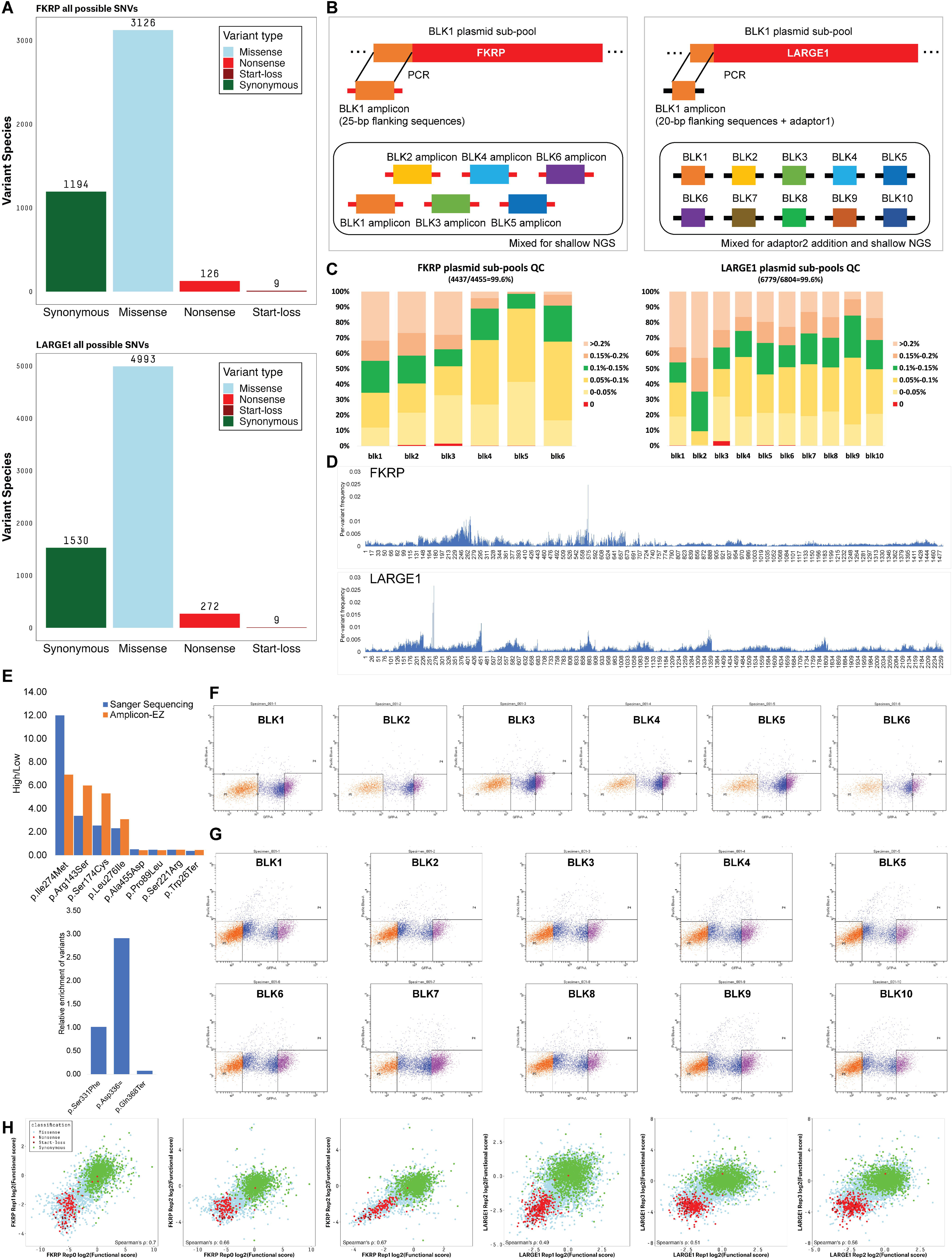
**The implementation of SMuRF** (A) We designed and synthesized oligos for all possible CDS SNVs of *FKRP* and *LARGE1*. Stop-loss variants were intentionally excluded from the design due to their incompatibility with the lentivirus-based saturation mutagenesis strategy. (B) Plasmid pool QC was performed using the Amplicon-EZ service. QC sequencing libraries were constructed using PCR amplification strategies for *FKRP* and *LARGE1*. BLK, block. (C) 99.6% of all possible SNVs were represented in the plasmid pools *FKRP* and *LARGE1*. Color groups indicate the variant representation in the pools. If the variants are evenly represented, we expect a 0.13% (*FKRP* BLK1-5) or a 0.15% (*FKRP* BLK6; *LARGE1* blocks) representation for all variants. (D) Per-variant frequency distribution along the CDS. x-axis represents the nucleotide sites. (E) Mini-libraries (Method S7) were used to optimize the FACS assay to achieve expected separation of variants with known clinical significance; *FKRP* (top), *LARGE1* (bottom). Both Amplicon-EZ NGS and Sanger sequencing were used to quantify the variant enrichment for *FKRP*. Sanger sequencing was used to quantify the variant enrichment for *LARGE1*. Relative enrichment was defined as a ratio of a variant’s representation in the high-glycosylation group to that in the low-glycosylation group. Higher relative enrichment indicates higher variant function. The *LARGE1* sequence was cloned from HEK293T cDNA. HEK293T carries a heterozygous *LARGE1* mutation (c.1848G>A, p.Met616Ile). This variant was removed when building the *LARGE1* SMuRF lentiviral pools but was kept in the experiment depicted here. Sanger sequencing confirmed the frequency of this mutation in these four lentiviral constructs were: WT 35.1%, S331F 96.3%, D336= 2.0%, Q368*=2.0%. (F and G) The high-glycosylation group (top ∼20%) and low-glycosylation group (bottom ∼40%) were gated based on the IIH6C4+FITC signal. Examples show the FACS gating parameters for all blocks of *FKRP* (F) and *LARGE1* (G). 20k events were recorded. Cell debris and multiplets were pre-excluded before the sorting. (H) The functional scores from the biological replicates exhibited good pair-wise correlation. The Spearman’s rank correlation rho is reported.

**Figure S3:**
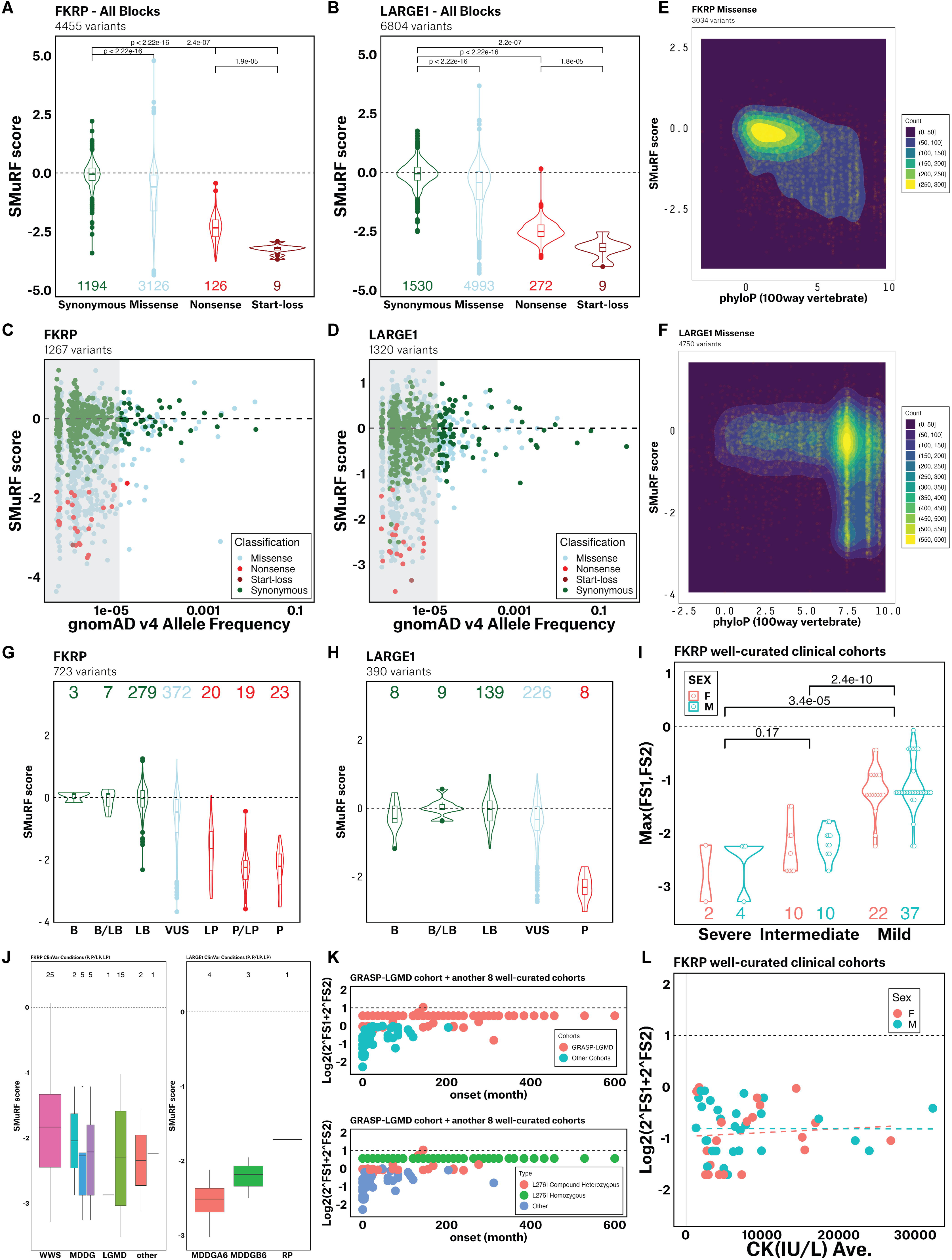
**SMuRF scores were employed to characterize *FKRP* and *LARGE1* variants** (A and B) SMuRF scores align with variant types (A, *FKRP*; B, *LARGE1*): synonymous variants resemble wildtype, nonsense variants consistently have low scores, and start-loss variants exhibit markedly lower scores than nonsense variants. Noteworthy outliers include high-scoring nonsense variants at the end of coding sequences. The box boundaries represent the 25th/75th percentiles, with a horizontal line indicating the median and a vertical line marking an additional 1.5 times interquartile range (IQR) above and below the box boundaries. p-values were calculated using the two-sided Wilcoxon test. Counts of variants were labeled below the boxes. (C and D) SMuRF revealed functional constraints based on variants reported in gnomAD v4.0.0 (both genome and exome sequencing data; C, *FKRP*; D, *LARGE1*): Low allele frequency variants had diverse functional scores, while high allele frequency variants converged towards wildtype due to selection pressures (Gray box: AF<1.5e-05). Dashed lines represent WT functional score. Dots were jittered with geom_jitter(width = 0.05, height = 0.05). (E and F) SMuRF scores were compared with the phyloP score calculated from multiple alignments of 100 vertebrate species (E, *FKRP*; F, *LARGE1*). The higher the phyloP score is, the more conserved a genomic site is. The missense variants show clustering around the WT SMuRF score (0) at less conserved sites, while they tend to be more disruptive at more conserved sites. Density was calculated with contour_var = "count" in R. (G and H) SMuRF scores correlate well with clinical classification in ClinVar (G, *FKRP*; H, *LARGE1*). B: Benign; B/LB: Benign/Likely benign; LB: Likely Benign; P: Pathogenic; P/LP: Pathogenic/Likely pathogenic; LP: Likely pathogenic; VUS: Uncertain significance. Counts of variants were labeled above the boxes. Dashed lines represent WT functional score. (I) The higher SMuRF score of the two scores of a variant pair is used to represent the variant pair. Scores associated with mild cases were significantly higher than those of intermediate and severe cases. Counts of cases were labeled below the boxes. p-values were calculated using the Wilcoxon test. FS1, the SMuRF functional score of the variant on Allele1; FS2, the SMuRF functional score of the variant on Allele2. (J) The SMuRF scores of variants classified as Pathogenic (P), Pathogenic/Likely Pathogenic (P/LP), and Likely Pathogenic (LP) in ClinVar did not show a significant pattern when grouped according to the reported conditions in ClinVar. (Left, *FKRP*; Right, *LARGE1*). WWS: Walker-Warburg syndrome. MDDG: muscular dystrophy dystroglycanopathies (left to right: type A1, type A5, type B5). LGMD: limb-girdle muscular dystrophy (left to right: not specified; type R9/2I). Other: not specified (left to right: Abnormality of the musculature, Cardiovascular phenotype). RP: retinitis pigmentosa. Counts of cases were labeled above the boxes. (K) Variability in onset time among patients with homozygous p.Leu276Ile mutation.Data from The Genetic Resolution and Assessments Solving Phenotypes in LGMD consortium (GRASP LGMD) were integrated with the 8 well-curated cohorts. The GRASP LGMD cohort mainly includes patients with homozygous p.Leu276Ile mutation and patients with compound heterozygous mutations (with p.Leu276Ile on one allele). Patients with homozygous p.Leu276Ile mutation exhibited a diverse range of onset ages, suggesting that the severity of dystroglycanopathy in these individuals might be influenced by other factors, like genetic modifiers, in addition to the *FKRP* mutation. (L) Creatine kinase (CK) values did not show a significant correlation with SMuRF scores. Spearman’s rank correlation rho: -0.04 (all data), -0.09 (male), 0.03 (female). Light grey rectangle indicates the normal range in healthy people (24.00 - 204.00 IU/L).

**Figure S4:**
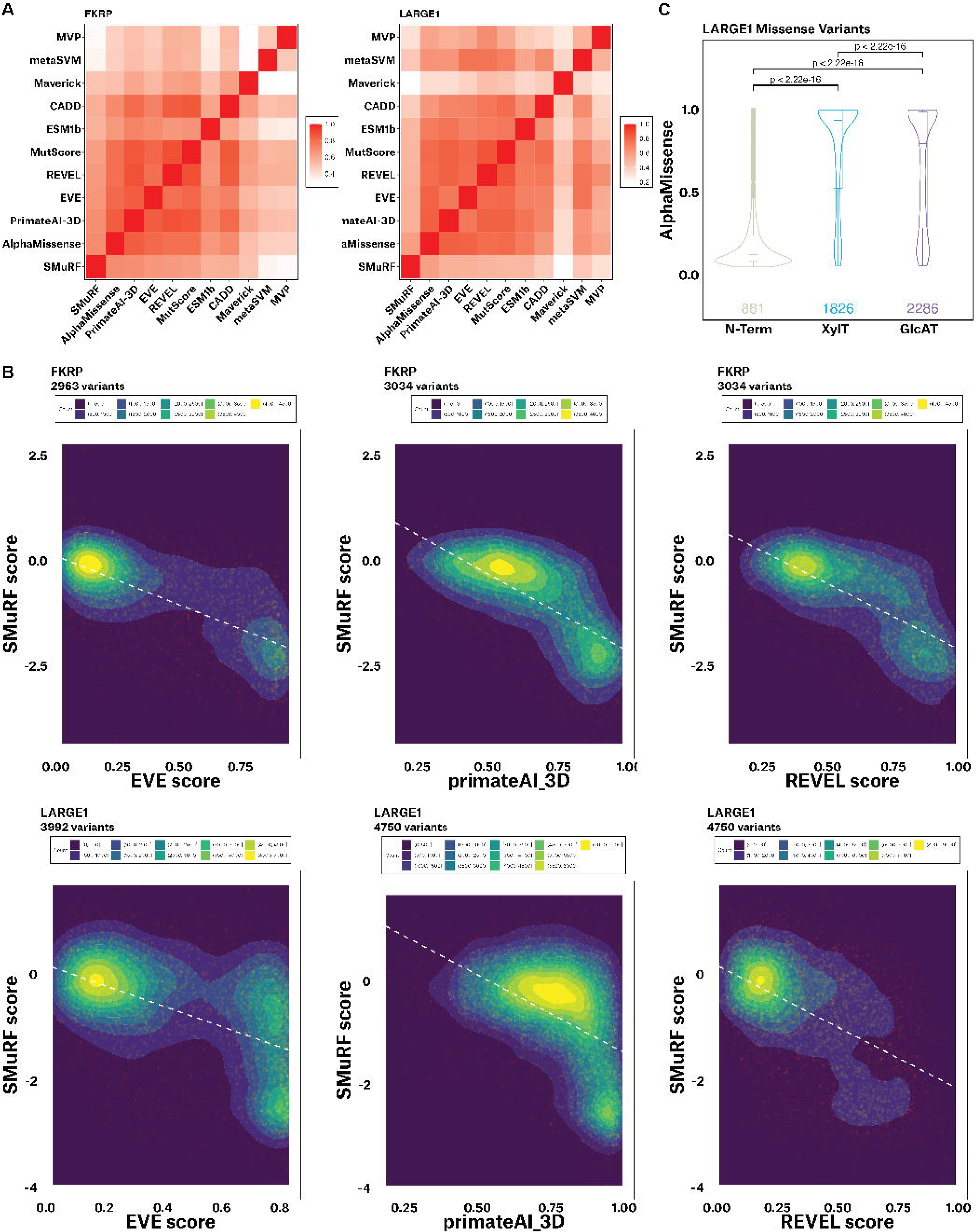
**Comparing SMuRF scores with the predictor scores.** (A) Top-ranked computational predictors correlate well with each other. Legend indicates the absolute Spearman’s rho. (B) Among the predictors examined, those based on evolutionary conservation exhibit high Spearman’s coefficients with SMuRF, such as EVE (*FKRP*: -0.67; *LARGE1*: -0.51) and PrimateAI-3D (*FKRP*: -0.69; *LARGE1*: -0.49). REVEL demonstrates the best performance according to the ROC curves and exhibits high Spearman’s coefficient with SMuRF (*FKRP*: -0.65; *LARGE1*: -0.47). White dashed lines represent linear regression. Density was calculated with contour_var = "count" in R. (C) AlphaMissense also exhibits functional differences between LARGE1 domains. Box plots depict the 25th/75th percentiles (box boundaries), median (horizontal line), and an additional 1.5 times IQR (vertical line) above and below the box boundaries. p-values were calculated using the Wilcoxon test. Counts of variants were labeled below the boxes.

**Figure S5:**
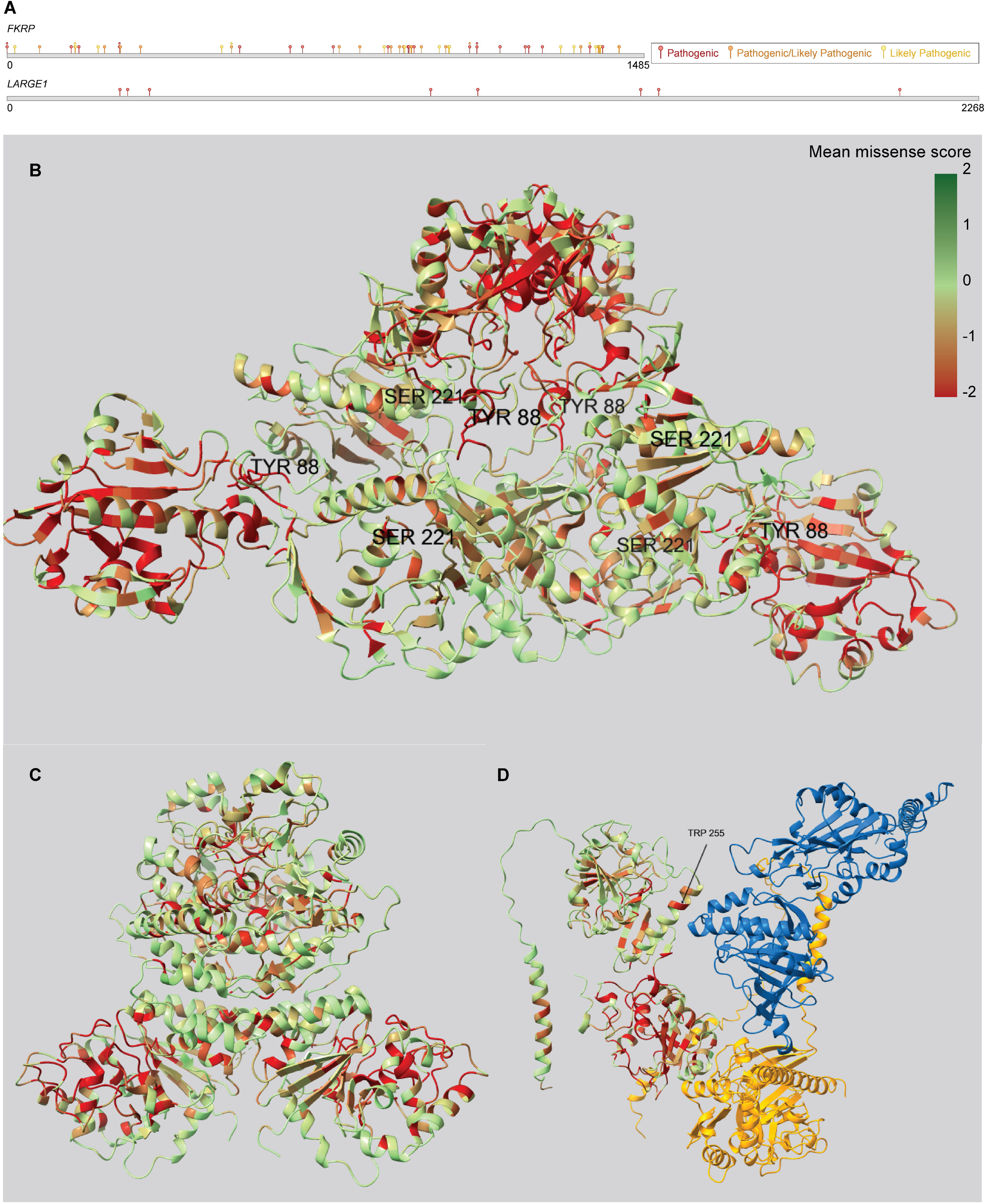
**SMuRF highlights the critical structural regions.** (A) The currently known disease-related SNVs in *FKRP* and *LARGE1* are dispersed throughout their entire sequences. Pathogenic, Pathogenic/Likely pathogenic, and Likely pathogenic variants in ClinVar (April 20, 2023) were labeled at their respective sites on the cDNA sequences. (B) The SMuRF scores of SNV-generated single amino acid substitutions were projected onto the FKRP tetramer (PDB 6KAM). p.Tyr88 (mean SMuRF=-2.69) and p.Ser221 (mean SMuRF=-1.00) are both situated at the subunit-subunit interface involved in FKRP tetramerization. (C) The SMuRF scores of SNV-generated single amino acid substitutions were projected to the LARGE1 dimer (PDB 7UI7). (D) The SMuRF scores of SNV-generated single amino acid substitutions were projected to the FKTN-FKRP-RXYLT1 complex predicted with Alphafold2 LocalColabFold (https://github.com/YoshitakaMo/localcolabfold). Blue: FKTN; orange: RXYLT1.

**Figure S6:**
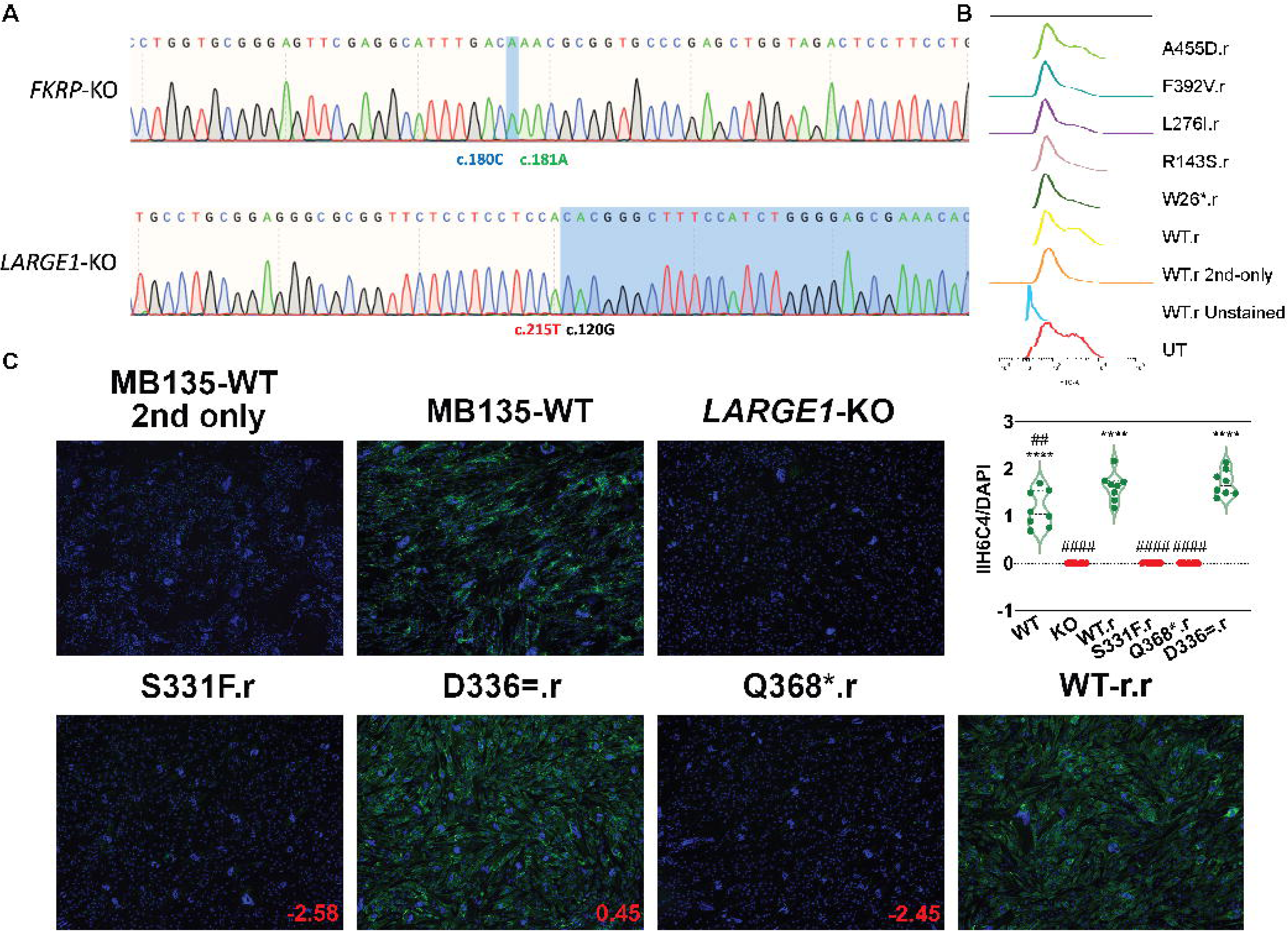
**Additional validations and analyses for SMuRF findings.** (A) Homozygous *GOI*-KO MB135 monoclonal cell lines were isolated from pooled CRISPR RNP nucleofected cells. *FKRP*-KO MB135 carries a 1-bp frameshifting insertion (c.181Adup). *LARGE1*-KO MB135 carries a 94-bp frameshifting deletion (c.121_214del). The mutations were validated with Sanger sequencing. These mutations are the same as the mutations in the *GOI*-KO HAP1 lines. (B) MB135 is not compatible with the flow cytometric assay due to high background noise. Monoclonal Lenti-*DAG1 FKRP*-KO MB135 myoblasts were used in this IIH6C4 flow cytometric experiment. High background noise was observed in both the control sample where no Lenti-*FKRP* rescue was performed (UT), and the control sample where the Lenti-*FKRP*-rescued cells were stained with secondary antibody but without the IIH6C4 primary antibody (WT.r 2nd-only). No significant differences were observed among different variants. (Monoclonal *FKRP*-KO MB135 myoblasts were also tested and showed the same pattern). (C) Validation of individual *LARGE1* variants using an IIH6C4 IF assay. “.r” denotes lentiviral transduction of an individual variant (Same lentivirus as used in Figure S2E). Green: IIH6C4, α-DG the glycosylation level. SMuRF scores are indicated in red. Immunofluorescence intensity was quantified using integrated density (IntDen) of IIH6C4 relative to DAPI by ImageJ. Analyses were conducted on 8 representative images from 2 of 3 independent experiments. ****p < 0.0001 (compared with LARGE1-KO group). ##p < 0.01, ####p < 0.0001 (compared with WT.r group). Multiple comparisons between groups were performed using analysis of variance (ANOVA) followed by Bonferroni post hoc test through GraphPad Prism 10.2.2.

**Figure S7:**
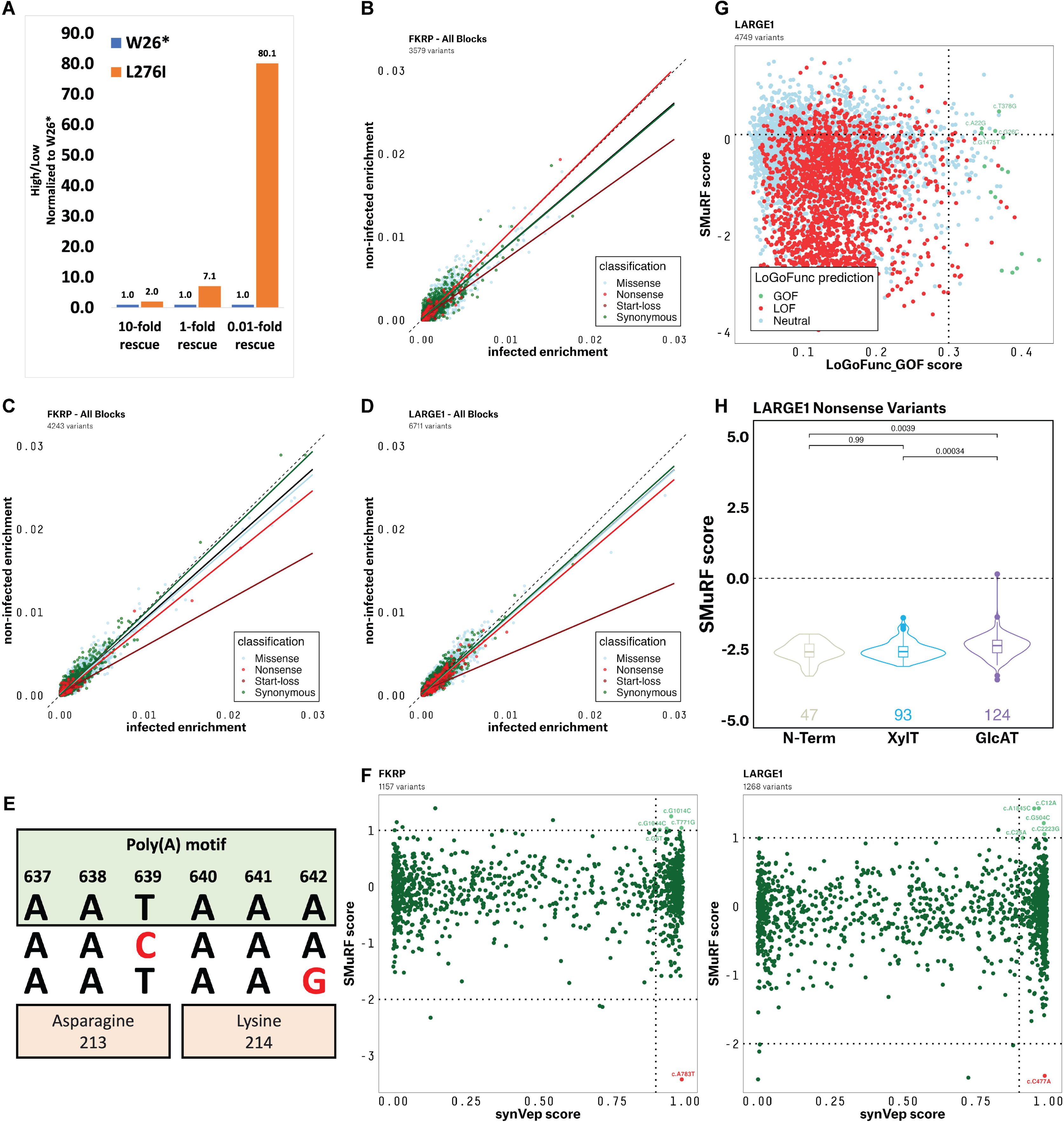
**Considerations for the implementation and further development of SMuRF.** (A) A small-scale FACS experiment utilizing an FKRP mini-library comprising WT, Trp26Ter, and Leu276Ile. y-axis is high/low enrichment normalized to Trp26Ter. Higher number indicates higher function. 1-fold transduction conferred lentiviral *FKRP* expression comparable to the endogenous *FKRP* expression in WT cells. Higher MOI appeared to reduce the functional difference between Trp26Ter and Leu276Ile. (B-D) The VSV-based viral entry assays did not exhibit sufficient sensitivity for large-scale evaluation of variants. Dashed line indicates no enrichment difference between the VSV-infected group and the non-infected group. Black solid line represent linear regression for all variants. Other solid lines represent linear regression for different variant types. B, rVSV-LASV-GPC viral entry assay for *FKRP* (The trend lines of missense variants and all variants overlay with the trend line of synonymous variants). ppVSV-LASV-GPC viral entry assay for *FKRP* (C) and *LARGE1* (D; The trend line of all variants overlays with the trend line of missense variants). The trend lines indicated a tendency where start-loss variants were the most enriched in the infected group, suggesting a greater disruptive effect on α-DG glycosylation. Nonsense variants showed the next highest enrichment, followed by missense variants, while synonymous variants demonstrated the lowest level of enrichment. However, overall, the VSV assays lack sufficient sensitivity for accurate interpretation of variants. (E) Two synonymous variants (red) in *LARGE1*, c.639T>C (p.Asn213=) (SMuRF=0.45) and c.642A>G (p.Lys214=) (SMuRF=0.64), may potentially have gain-of-function (GOF) effects by removing a poly(A) motif from the coding sequence. However, validation using the orthogonal ppVSV assay did not support the hypothesis of these 2 synonymous variants having GOF effects (Figure 6E). (F) Higher synVep scores indicate stronger effects, either loss-of-function effects (LOF) or gain-of-function (GOF) effects. Light green: potential GOF variants. Red: Potential LOF variants. (G) Comparing SMuRF scores with GOF scores generated by LoGoFunc, an ensemble machine learning-based predictor. LoGoFunc classifies the variants into 3 categories: GOF, LOF, or Neutral. LoGoFunc_GOF score is a probability for GOF. (H) Nonsense variants in the GlcAT region of *LARGE1* generally exhibit higher SMuRF scores than those in the XylT region. Box plots depict the 25th/75th percentiles (box boundaries), median (horizontal line), and an additional 1.5 times IQR (vertical line) above and below the box boundaries. p-values were calculated using the two-sided Wilcoxon test. Counts of variants were labeled below the boxes. Dashed lines represent WT functional score.

**Table S6:** Spearman’s rank correlation rho of SMuRF and the computational predictors.

| Gene | AlphaMissense | PrimateAI-3D | EVE | REVEL | MutScore |
| --- | --- | --- | --- | --- | --- |
| *FKRP* | -0.6978703 | -0.6903724 | -0.6659107 | -0.6507694 | -0.6342445 |
| *LARGE1* | -0.5407195 | -0.4947909 | -0.5132787 | -0.4741103 | -0.4700939 |

| Gene | ESM1b | CADD | MAVERICK | metaSVM | MVP |
| --- | --- | --- | --- | --- | --- |
| *FKRP* | 0.5655441 | -0.5813337 | -0.5043554 | -0.33925 | -0.3255476 |
| *LARGE1* | 0.4767206 | -0.4000735 | -0.2382434 | -0.4111915 | -0.2704415 |

